# Geographic Generalization in Airborne RGB Deep Learning Tree Detection

**DOI:** 10.1101/790071

**Authors:** Ben. G. Weinstein, Sergio Marconi, Stephanie A. Bohlman, Alina Zare, Ethan P. White

**Affiliations:** Department of Wildlife Ecology and Conservation, University of Florida, Gainesville, Florida, USA; School of Forest Resources and Conservation, University of Florida, Gainesville, Florida, USA; Department of Electrical and Computer Engineering, University of Florida, Gainesville, Florida, USA

**Keywords:** Tree Crown Detection, RGB Deep learning, object detection, Airborne LiDAR

## Abstract

Tree detection is a fundamental task in remote sensing for forestry and ecosystem ecology applications. While many individual tree segmentation algorithms have been proposed, the development and testing of these algorithms is typically site specific, with few methods evaluated against data from multiple forest types simultaneously. This makes it difficult to determine the generalization of proposed approaches, and limits tree detection at broad scales. Using data from the National Ecological Observatory Network we extend a recently developed semi-supervised deep learning algorithm to include data from a range of forest types, determine whether information from one forest can be used for tree detection in other forests, and explore the potential for building a universal tree detection algorithm. We find that the deep learning approach works well for overstory tree detection across forest conditions, outperforming conventional LIDAR-only methods in all forest types. Performance was best in open oak woodlands and worst in alpine forests. When models were fit to one forest type and used to predict another, performance generally decreased, with better performance when forests were more similar in structure. However, when models were pretrained on data from other sites and then fine-tuned using a small amount of hand-labeled data from the evaluation site, they performed similarly to local site models. Most importantly, a universal model fit to data from all sites simultaneously performed as well or better than individual models trained for each local site. This result suggests that RGB tree detection models that can be applied to a wide array of forest types at broad scales should be possible.

## 1. Introduction

Tree detection is a critical step in remote sensing of forested landscapes. Identifying individual crowns in airborne imagery allows ecologists, foresters, and land managers to increase the extent of sampling compared to terrestrial surveys. While many LIDAR-based tree segmentation algorithms have been proposed (Aubry-Kientz et al., 2019), the field has been slow to adopt automated methods due to concerns over accuracy, transferability and transparency (Vaglio Laurin et al., 2019). As a result, existing methods are rarely evaluated on multiple forests simultaneously, making it unclear how they will perform in the novel contexts required for large scale application. The availability of LIDAR data can also be limiting for large scale application. In contrast, RGB imagery is more widely available, but relatively few RGB algorithms have been proposed (González-Jaramillo et al., 2019) due, in part, to challenges with closed canopies and the diverse appearance of trees across forest types.

Current tree segmentation approaches are primarily based on user-defined algorithms that describe the appearance of trees in a hierarchical sequence of rules. These rule-based approaches rely on combinations of shape features (Gomes et al., 2018), template matching (Dai et al., 2018), network analysis (Williams et al., 2019), and watershed routines (Silva et al., 2016) that are applied to either LIDAR point clouds or RGB photogrammetric imagery (Brieger et al., 2019). By describing the parameters that define an individual tree, unsupervised algorithms attempt to match these rules when predicting unlabeled data. This can make applying these algorithms across different forest types challenging because the rules describing a tree vary depending on the type of forest, leading to overfitting for a particular geographic area. For example, some methods use allometric relationships between crown area and tree height to improve algorithm performance (Coomes et al., 2017; Williams et al., 2019), but these relationships vary with forest type and species. Recent attempts to mitigate this variation have used approaches that choose from a pool of potential tree shapes (Gomes et al., 2018). However, the need to define the full pool of possible tree shapes before analyzing each new site will be prohibitive over large geographic areas that incorporate diverse assemblages. As a result of these limitations, most tree detection algorithms have been applied and tested on similar forest types with little exploration of how the algorithms generalize to other natural settings. Therefore, despite the intense work in airborne tree detection over the last decade (Coomes et al., 2017; Heinzel and Huber, 2018; Jakubowski et al., 2013; Li et al., 2012; Williams et al., 2019), there remains no clear consensus on best practices (Aubry-Kientz et al., 2019).

Within the field of computer vision, there has been a broad shift away from rule-based approaches (i.e., user-designed features) towards approaches that learn features from data using deep learning neural networks (Agarwal et al., 2018). There have been few attempts to use learned features in tree detection (Dai et al., 2018) due to the need for a large amounts of labeled training data, which is often difficult or impossible to collect in ecological contexts. Overall, generalization of deep learning algorithms across applications in airborne remote sensing remains a challenging task (Zhu et al., 2017). A typical neural network has millions of parameters and is therefore at risk of overfitting when using small datasets. Given the diversity of trees, finding general features will require a combination of large training datasets and algorithmic approaches that allow the neural networks to learn the combination of features that characterize trees across forest types.

Weinstein (et al. 2019) recently developed a deep learning approach for tree detection using RGB data that has the potential to address these requirements for identifying trees across forest types. The semi-supervised method first uses unsupervised LIDAR-based tree detection (e.g., Silva et al. 2016) to generate millions of labeled trees that are used to pretraining of the neural network. This pretraining stage is followed by retraining the network based on a small number of high-quality hand-annotations. This addresses the need for large training data by generating millions of annotations of moderate quality for model pretraining and the method has been shown to perform well on a single oak woodland site. Due to its deep learning architecture, this method has the potential to learn general features of trees across forest types, but this remains an untested possibility.

Here we build on Weinstein et al. (2019) explore the potential of this tree detection method to generalize across sites by evaluating its performance on a range of forest types, assessing the transferability of tree features across forest types, and exploring the possibility of building a single unified tree detection model. Our aim is to test a deep learning approach 1) for identifying trees in four different forest types when trained on that forest type (‘within-site’); 2) for identifying trees when trained on data from other forest types (‘cross-site’); 3) for combining pretraining data from other sites with hand-annotated data from a new site (‘transfer learning’); and 4) for comparing the performance of a within-site model to a universal model fit to data on all forest types simultaneously (‘universal’). We also explore the sensitivity of the self-supervised method to the number of hand annotations, to determine the amount of time-intensive work needed to produce accurate results. By answering these questions, we will improve our understanding of the potential for universal tree detection methods and potentially advance RGB-based tree detection from algorithm development to large scale application for better understanding forests at scale.

## Methods

### Data Collection and Site Descriptions

The aerial remote sensing data products were provided by the National Ecology Observation Network (NEON) Airborne Observation Platform. We used the NEON 2018 “classified LiDAR point cloud” data product (NEON ID: DP1.30003.001) and the “orthorectified camera mosaic” (NEON ID: DP1.30010.001). The LiDAR data consist of 3D spatial point coordinates with an average of 4-6 points/m^2^. These data provide high resolution information about crown shape and height. The RGB data are a 1km x 1km mosaic of individual images with a cell size of 0.1 meters. All data are publicly available on the NEON Data Portal (http://data.neonscience.org/). For hand-annotations, we selected two 1km x 1km RGB tiles and used the program RectLabel (https://rectlabel.com/) to draw bounding boxes around each visible tree. For a count of tree annotations per site, see Table 1. All code for this project is available on GitHub (https://github.com/weecology/DeepLiDAR) and archived on Zenodo, and all annotations are available as part of the forthcoming NEON Tree Benchmark (https://github.com/weecology/NeonTreeEvaluation).

**Table 1.** The number of tree annotations used for semi-supervised pretraining, retraining and evaluation. Pretraining annotations are generated automatically using a LiDAR-based unsupervised algorithm. Training and evaluation annotations were hand-drawn.

| Forest Type | Pretraining Annotations | Training Annotations | Evaluation Annotations |
| --- | --- | --- | --- |
| Oak Woodland | 550,905 | 2,533 | 293 |
| Mixed Pine | 2,522,855 | 3,405 | 747 |
| Alpine | 3,121,036 | 9,730 | 1,699 |
| Eastern Deciduous | 3,131,283 | 1,231 | 489 |

We selected four sites from the NEON network to capture a range of canopy complexity and forest types. The ‘Oak Woodland’ is the San Joaquin Experimental Range, California. The site contains live oak (*Quercus agrifolia*), blue oak (*Quercus douglasii*) and foothill pine (*Pinus sabiniana*) forest. The majority of the site has a single-story canopy with mixed understory of herbaceous vegetation. The “Mixed Pine” site is Teakettle Canyon, California (37.00583, -119.00602) which contains red fir (*Abies magnifica*) and white fir (*Abies concolor*), jeffrey pine (*Pinus jeffreyi*) and Lodgepole Pine (*Pinus contorta*). The “Alpine” site is Niwot Ridge Mountain Research Station, Colorado (40.05425, -105.58237). This high elevation site (3000m) is near treeline with clusters of subalpine fir (*Abies lasciocarpa*) and englemann spruce (*Picea engelmanii*). Finally, the “Eastern Deciduous” site is the Mountain Lake Biological Station, Virginia (37.37828, -80.52484). Here the dense canopy is dominated by red maple (*Acer rubrum*) and white oak (*Quercus alba*). Each site presents its own challenges, with broad flat-topped trees in the Oak Woodland, tight clusters of trees in the Mixed Pine forest, thin conifers in the Alpine forest, and completely connected crowns in the Eastern Deciduous forest.

For each site, we manually annotated training tiles using the program RectLabel (Table 1). Training tiles were selected at random from the NEON data portal. At higher tree density sites, we cropped the 1km^2^ tiles to create more tractable sizes for hand-annotation. To enforce a minimum size threshold for tree annotations, we compared the hand-annotations to a LiDAR canopy height model and removed any trees less than 3m in height. The resulting annotations were compared to the LiDAR point cloud for further assessment. No attempt was made to delineate understory trees that were not visible in the RGB imagery.

For model evaluation, we used the NEON “tower” plots, which are a set of 40x40m plots placed throughout each site. For the Eastern Deciduous site, it was difficult to determine tree boundaries in both the RGB and LiDAR images. For this site, we overlaid a 1m resolution three-band hyperspectral composite image to highlight differences among co-occurring tree species in the area. The composite image came from NEON’s orthorectified surface reflectance (ID: DP1.30006.001) and contained bands in the infrared (940nm), red (650nm), and blue (430nm) spectrum. This allowed us to more accurately annotate the training and evaluation data in the closed canopy conditions.

### LiDAR tree Detection

We tested three existing unsupervised LiDAR algorithms (Dalponte and Coomes, 2016; Li et al., 2012; Silva et al., 2016), as implemented in the lidR R package (Roussel and David Auty, 2019), as both a comparison to the semi-supervised approach, and as potential algorithms to generate tree labels for the self-supervised portion of the workflow. We selected the best performing method (Silva et al., 2016) to create initial tree predictions in the LiDAR point cloud. This approach uses a canopy height model and an allometry of tree height to crown width to cluster the LiDAR cloud into individual trees. We used a canopy height model of 0.5m horizontal resolution to generate local treetops and an allometry of 90% of crown diameter to height for deciduous forests (Oak Woodland and Eastern Deciduous) and 20% of crown diameter to height for the coniferous forests (Mixed Pine and Alpine). LiDAR algorithms perform segmentation on a per-point basis, so we converted the output to a bounding box that covered the entire set of LiDAR points assigned to each tree to create training data equivalent to the hand annotated bounding boxes.

### Semi-supervised Deep Learning

We used our previously developed self-supervised algorithm for RGB-based tree identification (Weinstein et al. 2019). This method uses the Retinanet one-stage object detector (Gaiser et al 2017) with a Resnet-50 classification backbone, which allows pixel information to be shared at multiple scales, from individual pixels to groups of connected objects. We used a Resnet-50 classification backbone pretrained on the ImageNet dataset (He et al., 2016). Since the entire 1km RGB tile cannot fit into GPU memory, we cut each tile into 40m by 40m windows with an overlap of 5% (n=729). The order of tiles and windows were randomized before training to minimize overfitting among epochs. To reduce potential spatial autocorrelation in tree appearance between evaluation plots and pretraining data, we removed any training tiles within 1km of an evaluation tile. Using the pool of unsupervised LiDAR-based tree predictions, we pretrained the network with a batch size of 20 on 2 Tesla K80 GPU for 5 epochs. To align these unsupervised classifications with the ImageNet pretraining weights, we normalized the RGB channels by subtracted the ImageNet mean from each channel. We then retrained the network using the hand-annotated data for 40 epochs. For more details of this semi-supervised approach see Weinstein et al. (2019).

### Model Evaluation

Using the evaluation plots, we chose two metrics to assess model performance. For comparison with the existing LiDAR-only implementations, we used precision and recall statistics with a bounding box marked as true positive if it had an intersection-over-union (IoU) of greater than 0.5. Intersection-over-union is the ratio of the area of bounding box overlap to the area of bounding box union between the predicted tree crown and the visually annotated crowns in the evaluation data. For each bounding box prediction, the deep learning model reports a confidence score between 0 and 1. To transform these scores into precision and recall statistics, we need to define a threshold of box scores to accept. As we lower the threshold for acceptance, a greater number of trees will be captured, but at the expense of decreased precision. To highlight this relationship, we showed the performance of the deep learning approach across all bounding box probability thresholds between 0 and 1 with an interval of 0.1.

While IoU precision and recall are intuitive statistics, they are reported separately and do not capture differences in bounding box confidence scores. When comparing the different generalization approaches, it is useful to have a single metric to compare. We used the Average Precision (AP) metric commonly used for object detection tasks in computer vision, which is the area under the precision-recall curve computed at the 11 fixed 0.1 intervals between 0 and 1 (Lin et al., 2017).

### Assessing Generalization, Transferability, and Universal Model Fit

To assess generalization among sites, we performed three types of experiments that used different combinations for hand-annotations and pretraining data (Figure 2). The first experiment is to use pretraining and hand-annotated data to predict the evaluation data from the same site (‘within-site’). The next setup is to use the pretraining data and hand-annotated from the same site to predict the evaluation data from a different site (‘cross-site’). For example, using each of the within-site models, we can test the ability for a model to predict tree conditions in each of the other geographic sites, creating a matrix of cross-site predictions. To assess generalization without local pretraining data, we tested a model training using pretraining data from all other sites, but hand annotations from the same site as the evaluation data (‘transfer-learning’). For example, the transfer learning model for Oak Woodland used the hand-annotations from Oak Woodland, but the pretraining data for Alpine, Mixed Pine, and Eastern Deciduous. Finally, to test the potential for a universal model, we tested a model pretrained on all sites, followed by retraining on all hand-annotations. We then compared this model with each of the within-site model to test whether the addition of data from other sites improved predictions of trees from the same site.

**Figure 1.**
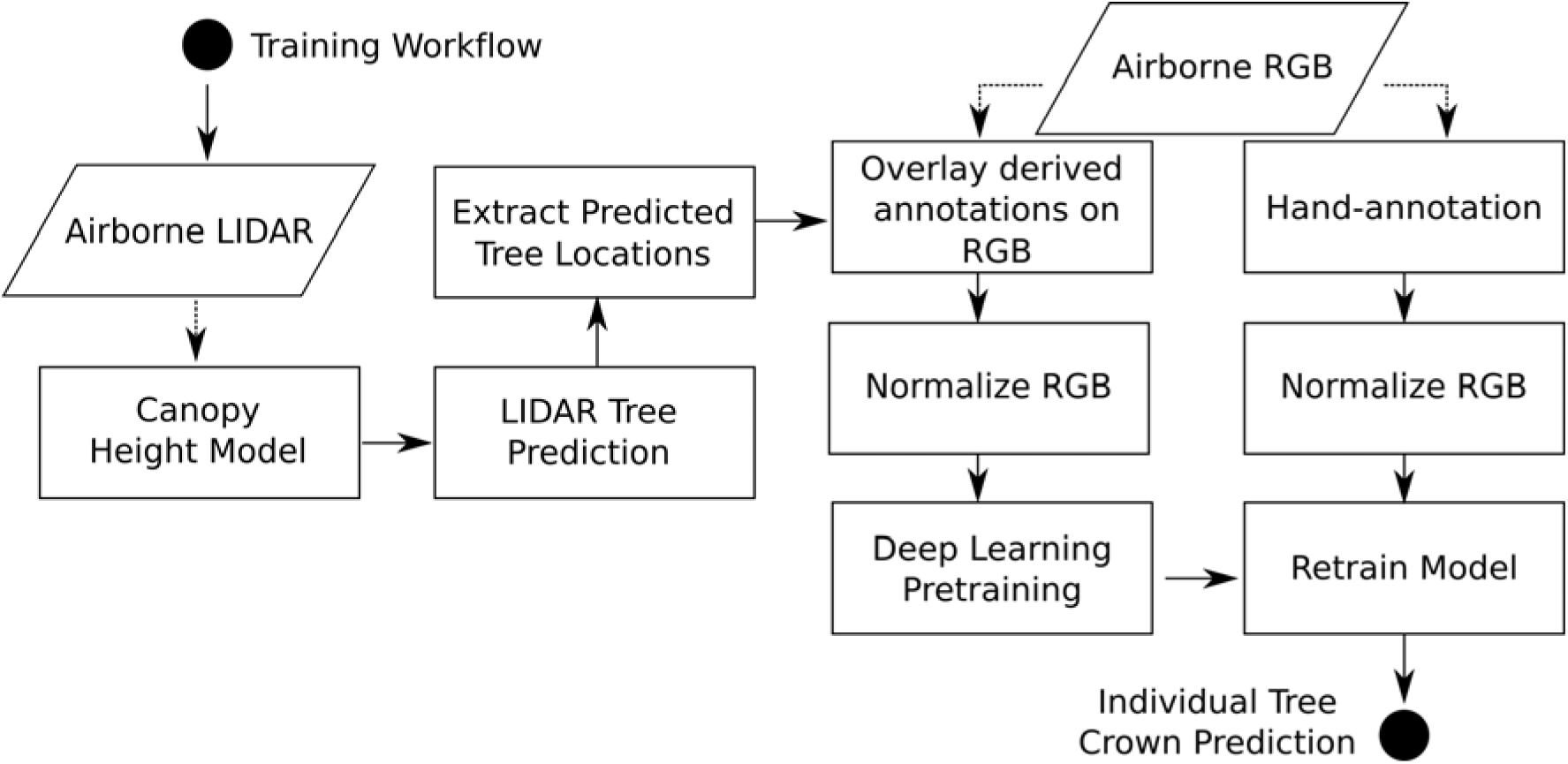
Conceptual workflow of proposed approach for airborne detection of individual tree crowns. Pretraining data is generated by overlaying predicted trees from a LIDAR-based unsupervised algorithm on to RGB imagery. These RGB images are used to pretrain a deep learning neural network. The resulting model is retrained based on RGB hand-annotations.

**Figure 2.**
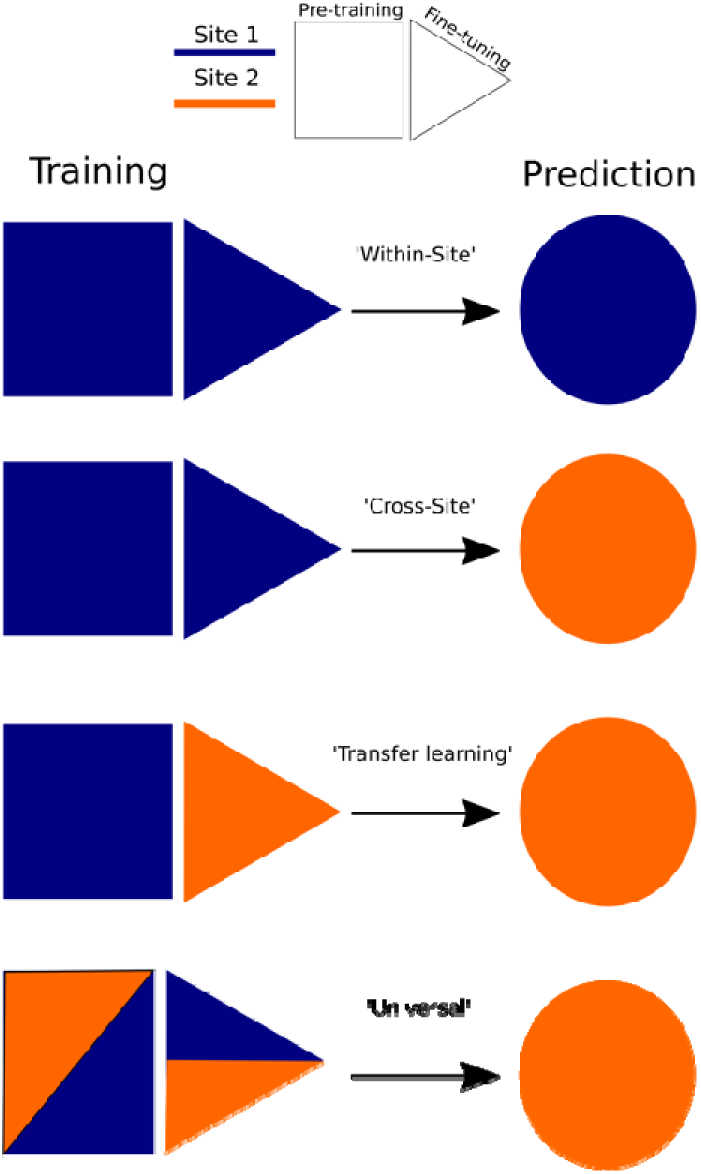
Approaches to geographic generalization in semi-supervised model training: 1) ‘Within-site’ training in which training data from site 1 is used to predict site 1; 2) ‘Cross-site’ training in which training data from site 1 is used to predict site 2; 2) ‘Transfer learning’ in which a model is first trained on site 1 data, followed by finetuning on site 2 training data, and 3) ‘Universal’ model in which training data from both site 1 and site 2 are used to predict evaluation data from site 2.

### Sensitivity to the Number of Hand-annotations

Collecting a sufficient number of training samples will often be a bottleneck in developing supervised methods in airborne imagery. It is therefore useful to test the number of local training samples needed to achieve peak performance. We performed a sensitivity study by training models using different proportions of training data. We selected 5%, 25%, 50% and 75% of the total hand-annotations to compare to the full dataset for the within-site results for each site. We reran this experiment five times to account for the random subsampling of annotations. In addition, we ran the evaluation plots for the pretraining model only (i.e. 0% hand-annotated data) to assess whether the addition of hand-annotated data improved the within-site pretraining.

## Results

Based on the highest performing probability cutoff (Figure 4), within-site predictions ranged from 0.60 recall and 0.75 precision in Mixed Pine to 0.34 recall and 0.55 precision in Alpine. The Oak Woodland and Mixed Pine sites consistently performed better than the Eastern Deciduous and Alpine sites. Visual inspection of the results showed that the vast majority of false positives were positively identified trees, but whose crown boundaries were either too large or too small for the intersection-over-union score of 0.5. Repeated training runs for each model showed relatively little variance, despite heterogeneity in tree types at all sites (Figure 4).

**Figure 4.**
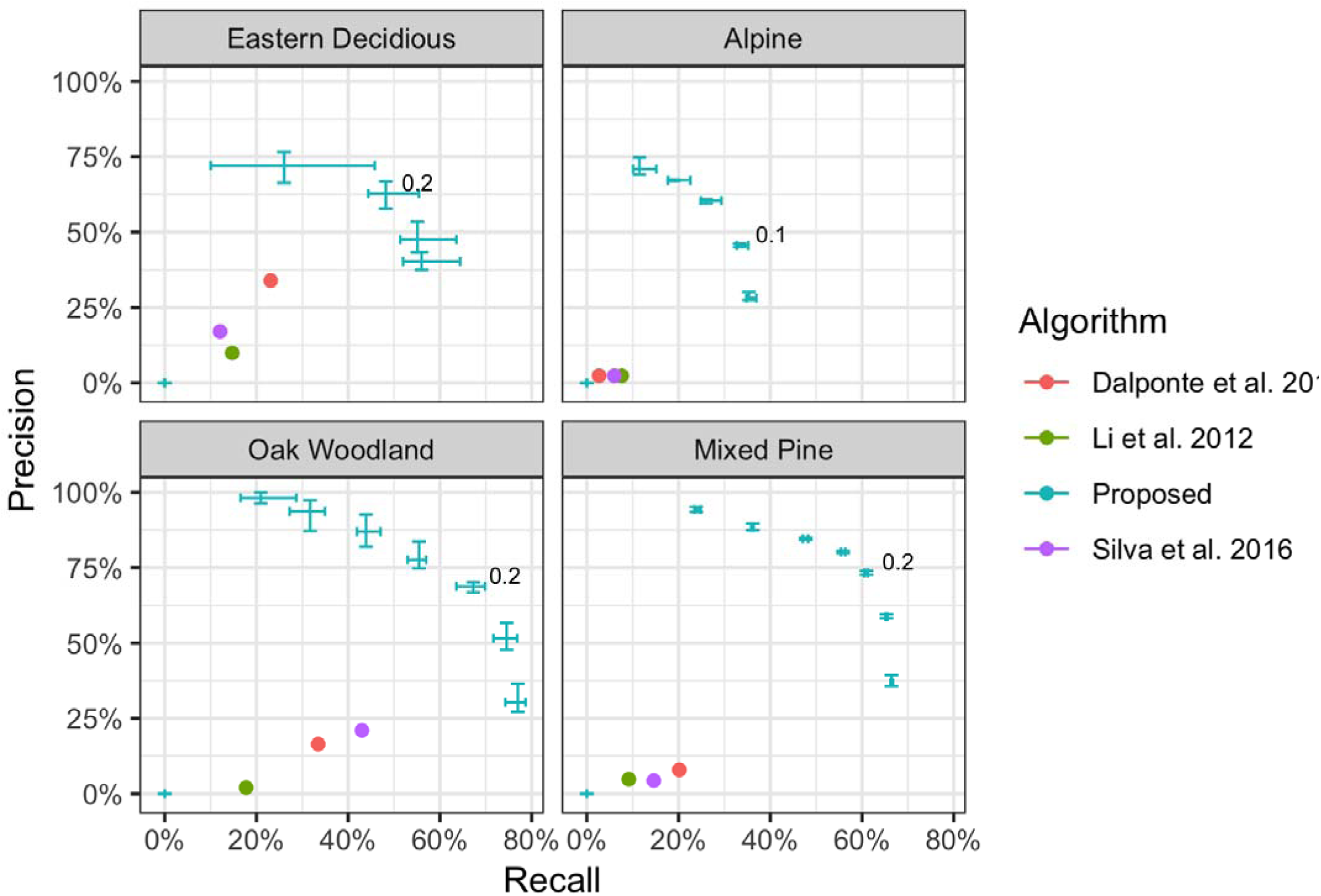
For each site, results of our proposed workflow compared to three existing LiDAR-only implementations from the commonly used lidR package. The proposed approach was evaluated at each of the 0.1 probability score intervals between 0-1. The probability threshold of the best performing model in our approach, calculated by f-score, is shown in black. Error bars show the variance in recall and precision based on five runs of hand-annotation training for each site.

The semi-supervised deep learning approach significantly outperformed the available LiDAR-only implementations from the lidR R package at all sites (Roussel and David Auty, 2019) (Figure 4). When comparing model performance with the Silva et al. (2016) algorithm used to generate the pretraining data, the deep learning model was more successful at delineating complex crown boundaries and avoiding clumping together small trees with narrow gaps (Figure 5). For the Oak Woodland site, the deep learning model was better able to capture crown area for the flat-topped canopy and avoided erroneously labeling bushes as trees. For the Eastern Deciduous site, the deep learning model more accurately found trees in the closed canopies, despite strong overlap in bounding box predictions and similarity in neighboring tree appearance. The Alpine site was the worst performing model, and many small trees were missed. This is likely due to the minimum anchor box size in the object detector and the arbitrary cutoff at 3m height for defining a ‘Tree’ class in the Alpine training data. To view predictions overlaid on each of the plots for the within-site models, see supplemental dataset S1.

**Figure 5.**
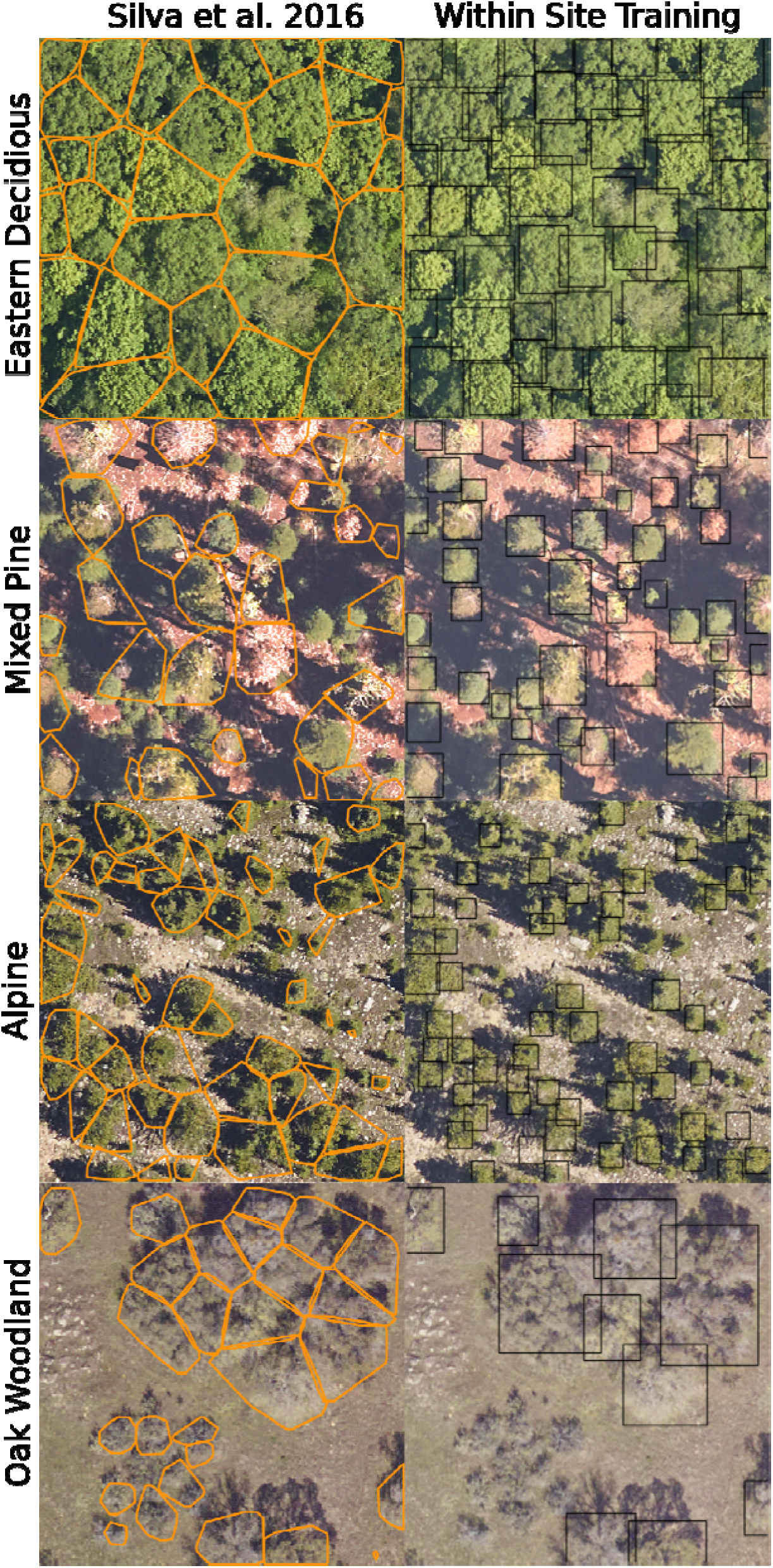
Example predictions for the LiDAR-only pretraining algorithm, and the deep learning detection network trained within-site.

When applying a model fit at one site to make predictions at other sites, we found generalization of the single-site models to be weak (Figure 7 A-E). Tree stems were often correctly identified among sites with similar forest conditions (Coniferous versus Deciduous), but the resulting crown boundaries were rarely accurate (Figure 6 – “Cross-Site”). The one exception was the prediction of Alpine evaluation plots using a model built from the Mixed Pine site. This model outperformed all other cross-site experiments and was superior even to the Alpine within-site model.

**Figure 6.**
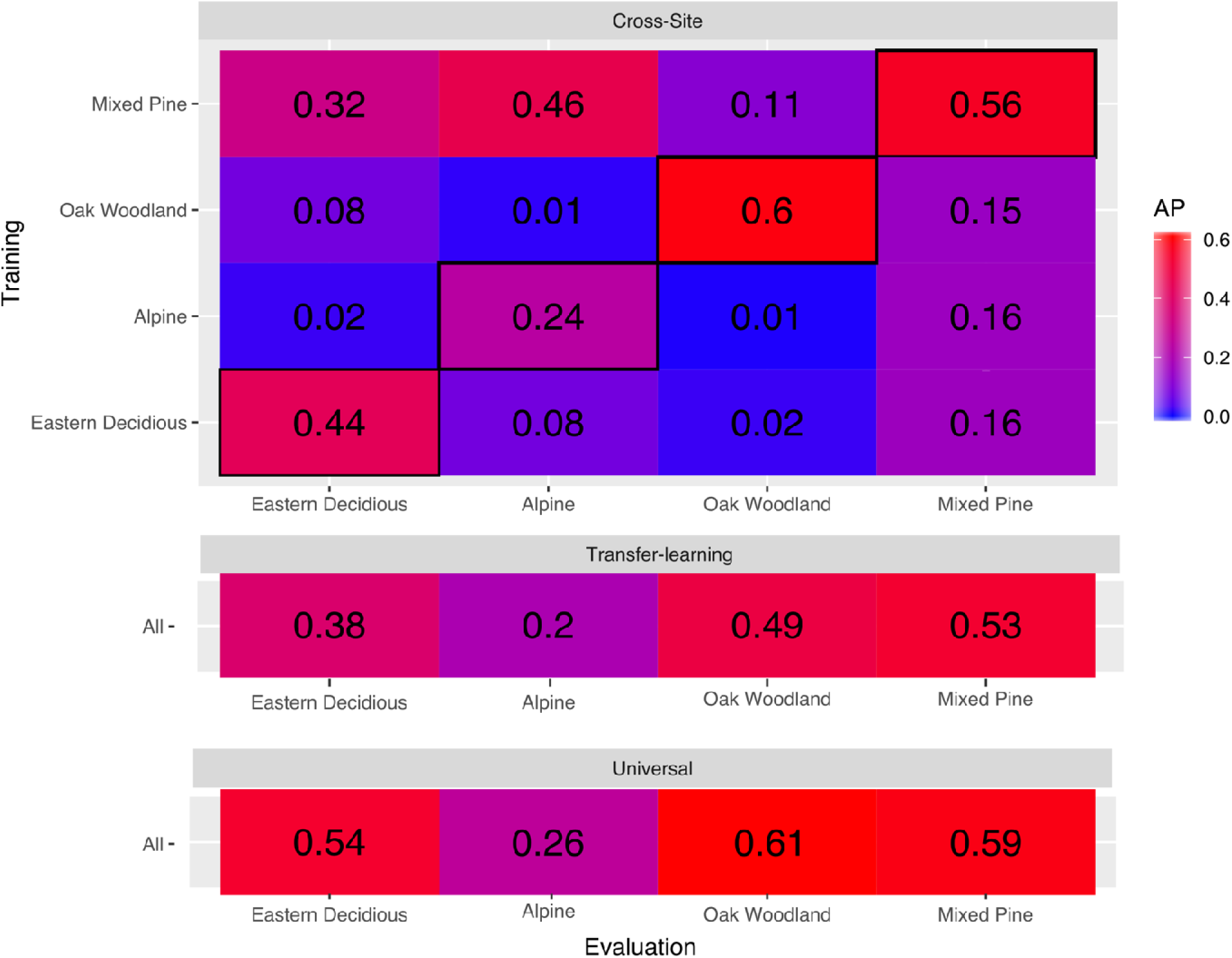
Comparison single-site, cross-site, transfer, and universal model performance based on Average Precision (AP). Single site predictions are on the bolded diagonal of the cross-site section and represent fitting and predicting on the same site. Cross-site predictions are for models trained on the one site (listed on the left side of the results matrix) and evaluated on a second site (listed across the bottom of the results matrix). Transfer learning takes a model pretrained on all sites except the focal site and retrained using the hand-annotations of the evaluation site. The universal model uses pretraining and hand-annotation data from all sites.

**Figure 7.**
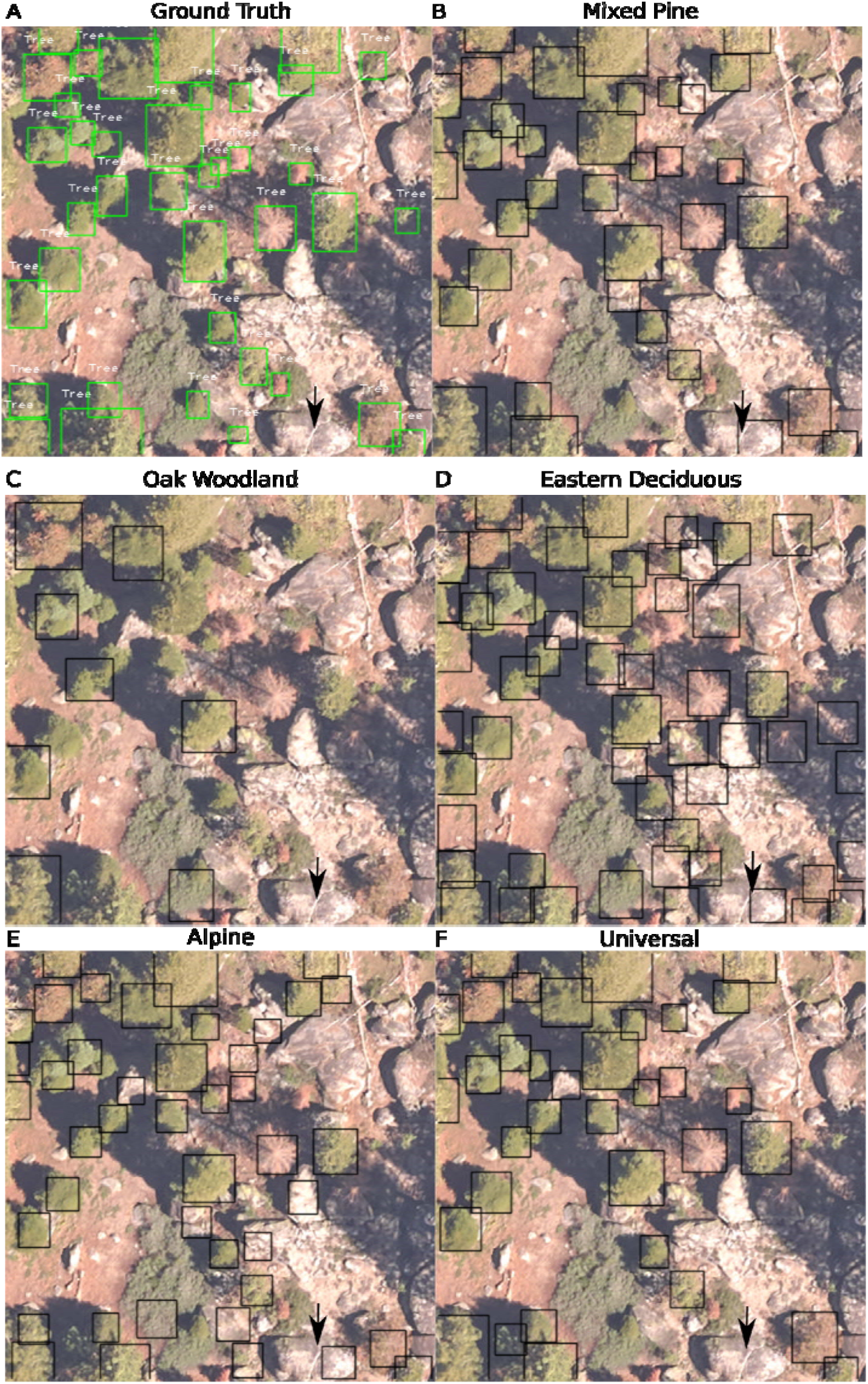
A sample evaluation plots from the Mixed Pine site predicted by a model built from training data from the same site, from each other site, and a universal model. Ground truth boxes are shown in green (A). Individual trees with a predicted probability greater than 15% are shown in black (B-E). The universal model (F) built from all annotations slightly outperformed all other models, including the model trained only from the Mixed Pine site. For example, the boulder in the bottom right corner is incorrectly classified as a tree by the models trained from Mixed Pine, Alpine, and Eastern Deciduous sites, but is correctly ignored in the Oak Woodland and Universal models.

**Figure 8.**
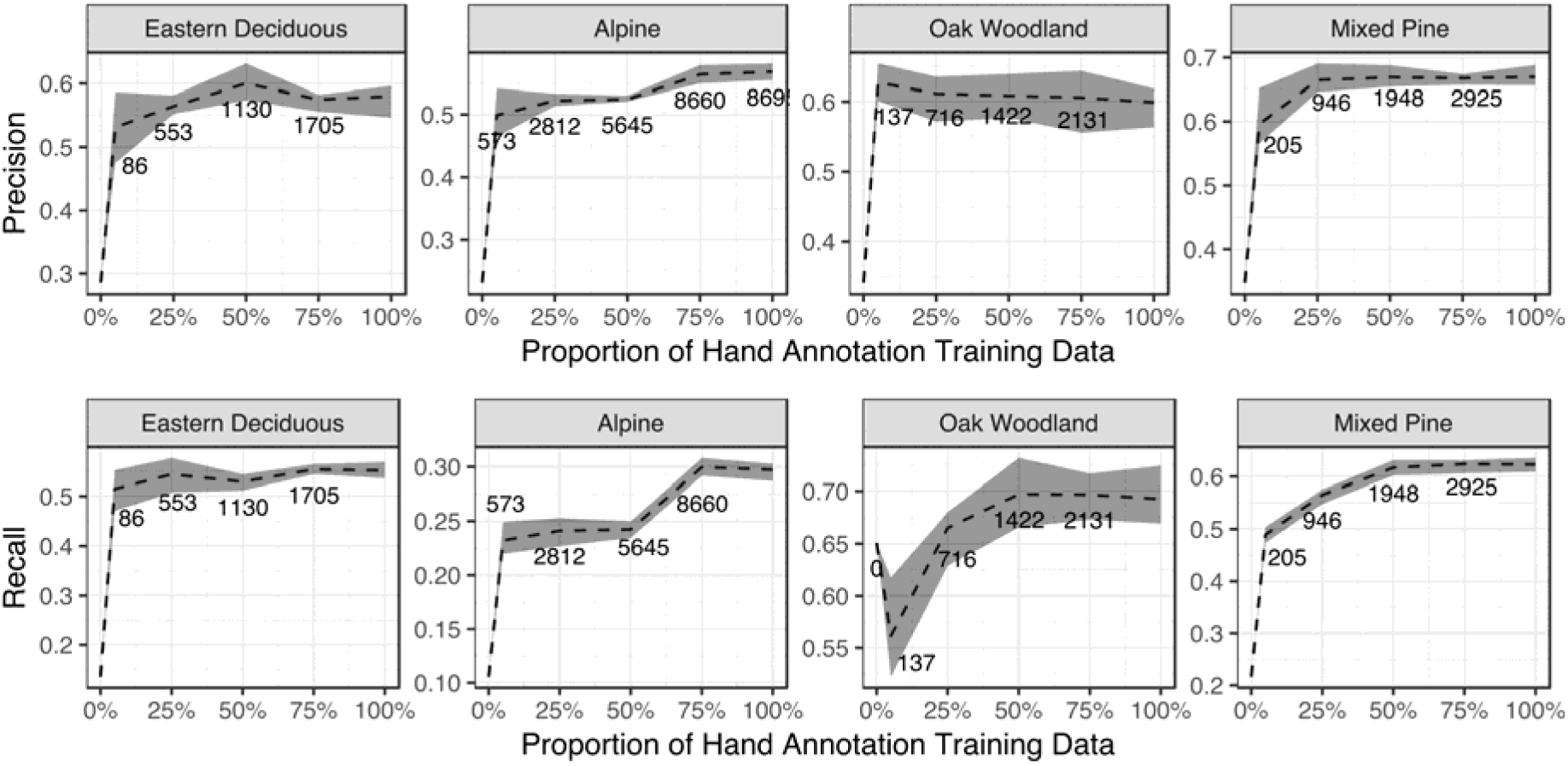
Sensitivity curves of the proportion of hand-annotation training data for each site. Values indicate the number of trees in the training dataset for each cutoff. Shaded area is the range of results from rerunning the analysis five times for each site. Note that due to the random sampling among runs, the exact number of trees will vary slightly. For simplicity, we show the mean number of training trees for each threshold.

Combining local hand-annotated data with unsupervised pretraining data from the other three sites demonstrated good transferability, with performance almost as good as using local pretraining data. The transfer learning experiments performed better than cross-site predictions for every site. This suggests that the pretraining model allows for generalized features that can be fine-tuned to local conditions.

Fitting a single universal model using data from all sites resulted in the best predictions for every individual site. The Average Precision of the Eastern Deciduous site improved from 0.44 to 0.54 (22%), Mixed Pine from 0.56 to 0.59 (5.4%), Alpine from 0.24 to 0.26 (8.3%) and Oak Woodland from 0.6 to 0.61 (1.6%). Visual inspection of predictions for the Mixed Pine site illustrates why the universal model improves performance. In Figure 7B, the within-site model (Mixed Pine) erroneously labels a large boulder in the bottom right hand corner of the image as a tree. This error was made in all other cross-site models, except for Oak Woodland. In the Universal model, this error was not made, suggesting that the universal model learned information about the background from other sites to improve predictions.

Assessment of the number of hand-annotations needed to improve model performance, indicated that while some hand-annotated data was important at all sites, the number of hand-annotated trees needed to improve model performance was typically relatively small. For example, the recall in the Mixed Pine site was < 0.2 with no hand-annotated data and was over 0.6 using approximately 2000 hand labeled crowns. Only minimal gains in performance occurred using up to an additional 2000 hand-annotated crowns. Overall, the shape of the ablation curves suggest that the model is fairly robust and needs only approximately 1000 crowns in most cases to create a model close to full performance. The exception is the Alpine model, which improved by more than 30% after 3,000 crowns. In general, the precision was more robust than recall, suggesting that the hand annotations mostly improve the predictions of crown boundaries rather than additional tree locations.

## Discussion

Airborne tree detection promises to unlock ecological and forestry data at unprecedented spatial extents compared to traditional ground surveys. To turn remote sensing data into ecological information, there is a need for a unified tree detection model that can be applied to a broad array of forest conditions. Using a semi-supervised deep learning approach, we trained individual tree detection models for four geographic sites and studied the transferability of tree features among forest types. Our results show significant improvements over commonly used LiDAR-only implementations. Despite challenging conditions including overlapping canopies and a range of acquisition environments, the proposed approach holds promise for automated tree location and size detection at scale. On average across sites, the universal model correctly identified crown extent with approximately 65% recall and 70% precision. The remaining false positives were almost always correctly detected individual trees, but whose crown boundaries did not meet the intersection over union score of greater than 0.5.

One goal was to assess the proposed crown detection approach in variety of canopy conditions to better understand which factors limit performance. We find performance is best in open environments with large, well-spaced, trees as in the Oak Woodland site. We had anticipated the performance of the algorithm would be worst at the closed canopy Eastern Deciduous site. However, it was at the Alpine site that the algorithm had the poorest performance, suggesting that short dense trees, rather than complex, interconnected tree boundaries are the biggest challenge. One possible explanation is that the trees in the Alpine site are more sensitive to the resolution of the RGB image due to their small size. Since we use an evaluation metric of intersection-over-union of 0.5, a difference of one pixel is inconsequential for large trees but may push small trees under the threshold for predicted positive.

One of the advantages of deep learning approaches to tree detection is the potential to learn cross-site features. We conducted three types of generalization experiments to assess the transferability among forest types. The first was to use models trained from one site to predict an unseen site. Prediction to unseen conditions is a challenging task in computer vision, especially when the sites were specifically chosen to represent distinct forest types. Overall, we saw a significant decrease in performance between cross-site and within-site models. This means that generalization between two forest types without local training data remains unlikely to provide acceptable results. The one exception was the prediction of the Alpine site, which had superior performance when predicted by the Mixed Pine site, rather than using the Alpine hand annotations. This may stem from the difficulty of hand annotating the small trees that are common in the Alpine site. It is possible that the model was better at transferring the features from the large conifers in Mixed Pine to the smaller conifers in Alpine than a human was in annotating the crown boundaries in Alpine. A second possibility is that the significant heterogeneity in the pretraining data for the Alpine site led to poor results. The LiDAR-based pretraining algorithm did not perform well at this site, with consistent under-segmentation among small trees. It is possible that the superior quality of the pretraining data at the Mixed Pine site allowed for better predictions in the Alpine site, compared to using lower quality data from the same site.

To provide the model with more information on local tree conditions, we conducted transfer learning experiments to assess whether models pretrained at other sites could be used in conjunction with training data from a local to site to fine tune the model to that site. We find that building from existing models of tree detection is a promising avenue towards cross-site generalization. Adding only a small amount of local training data greatly increased performance and nearly recovered performance of the within-site model. The results were mostly logical; combining pretraining from a deciduous site (e.g. Oak Woodland) to predict another deciduous site (e.g. Eastern Deciduous) is better than using pretraining from a coniferous site (e.g. Alpine). This opens up the possibility of regional tree detection models that connect ecotypes based on their dominant canopy structure and species.

The ultimate goal of the proposed approach is to move toward a single unified model that can produce individual tree predictions in a variety of ecosystems. Our analysis shows promising results for a universal model trained from all pretraining and hand annotations from every site. In all sites, a universal model provided equivalent or better predictions than a within-site model, with improvements of up to 20% in one site. Given that the sites were selected to be as different as possible, and encompass a range of tree canopy conditions, this result highlights the ability of convolutional neural networks to learn flexible deep features. We expect that as more sites are included, the universal model will continue to improve. This means that a way forward is to combine pretraining from as many sites as possible. Given that each NEON site has millions of trees, and there are dozens of sites with trees collected annually, there is a possibility of pretraining on continental scale. Further work is needed to know the balance between the number of training images per site and the number of sites to most efficiently train generalized features.

In addition to universal model development, transferring knowledge beyond the NEON sites may be useful for many applied problems. It is currently unknown to what extent features learned from the 0.1 m resolution data used here can be applied to lower resolution satellite data (Karlson et al., 2014) or higher resolution UAV data (Brieger et al., 2019). Cross resolution training has not been fully explored in environmental remote sensing, but Li et al., (2018) recently showed that deep learning networks can learn scale invariant land classifications that can be matched among data sources. Given the ability to collect virtually unlimited pretraining data using our semi-supervised approach, NEON sites can be seen as an ideal training sources for RGB tree models that could then be applied to other data types.

Our semi-supervised deep learning method uses LiDAR-based pretraining and RGB deep learning to perform individual tree segmentation (Weinstein et al. 2019). The NEON Airborne platform also collects hyperspectral information that may improve generalization across sites with similar species composition. Due to foliar and physical properties, tree species often have distinct spectral signatures which may facilitate distinguishing adjacent tree crowns. Hyperspectral features for tree species classification are relatively common (e.g. Maschler et al., 2018), but few papers have focused on integrating hyperspectral data into tree detection alongside other sensors. Hyperspectral data is available for all NEON sites, and we utilized a three-band composite image to assist in annotating the Eastern Deciduous site (Figure 9), illustrating the usefulness of hyperspectral data to distinguish adjacent tree crowns with human vision. Choosing the best way to represent high-dimensional hyperspectral data in conjunction with the LiDAR and RGB data is non-trivial and will be important for improvements in individual tree detection at broad scales.

**Figure 9.**
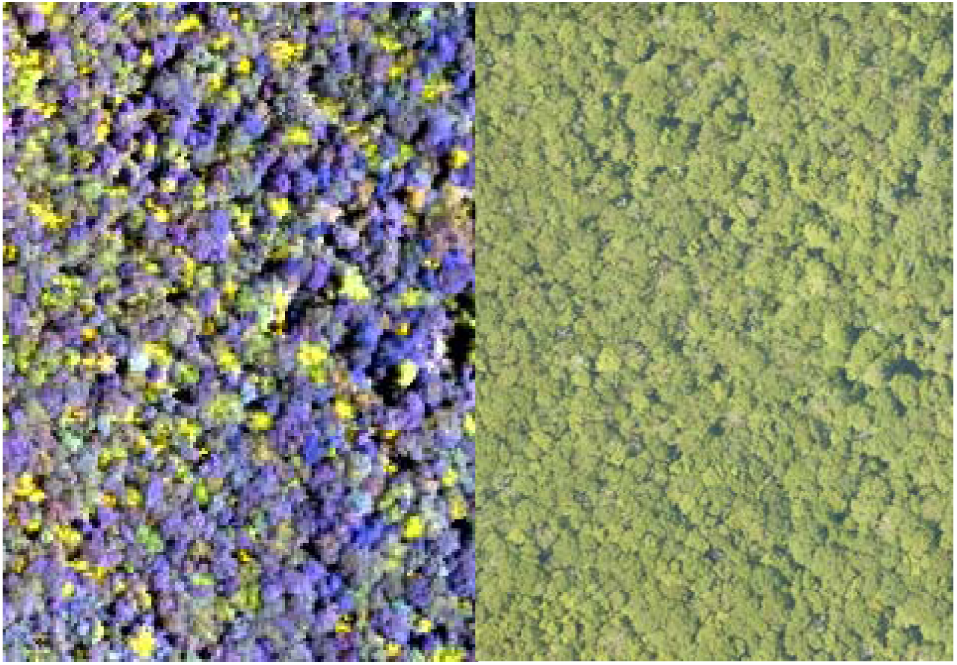
Composite hyperspectral image and corresponding RGB image for the Eastern Deciduous site. The composite image contained near infrared (940nm), red (650nm), and blue (430nm) bands. Forests that are difficult to segment in RGB imagery may be more separable in hyperspectral imagery due to the differing foliar chemical properties of co-occurring trees.

Methods to extract ecological information from airborne sensors are maturing due to advancements in computer vision, data availability and sensor quality. Given our results, what are the strengths and limitations ecologists should consider when adding airborne-derived data? Our results, and prior works (Aubry-Kientz et al., 2019), suggest that small and subcanopy trees will likely be overlooked. We therefore expect that studies in which the results are driven by the upper canopy, sun-exposed trees will benefit the most from remote sensing at broad scales. For example, the total amount of biomass in most forests depends strongly on the largest trees and will be less sensitive to potential non-detections of smaller subcanopy trees (Asner et al., 2012, Stegen et al. 2011, Bastin et al. 2018). The inclusion of RGB data may benefit existing large-scale LiDAR-based studies of tree growth (Caughlin et al., 2016), taxonomy (Féret and Asner, 2012) and disturbance (Garcia Millan and Sanchez-Azofeifa, 2018), since improved individual segmentation will lead to a more accurate matching of individual trees to metadata on taxonomy and health status. Studies of post-landscape disturbance, such as post-fire, will be aided by the broader perspective of airborne data, as well having significantly reduced risk compared to field surveys in recently burned forests. Most disturbances, such as fire and windstorms, alter the size distribution of forests, including large trees, and thus our approach can provide valuable, detailed landscape scale information about disturbance intensity and impacts (Kamoske et al., 2019). To address these questions, we envision a future in which airborne data on tree locations and sizes are a complement to local field surveys in broadening the scale of sampling in complex landscapes.

## Acknowledgements

This research was supported by the Gordon and Betty Moore Foundation’s Data-Driven Discovery Initiative through grant GBMF4563 to E.P. White and by the National Science Foundation through grant 1926542 to E.P. White, S.A. Bohlman, A. Zaire, D.Z. Wang, and A. Singh. This work was supported by the USDA National Institute of Food and Agriculture, McIntire Stennis project 1007080.

