## Supplementary material for "Geographic Generalization in Airborne RGB Deep Learning Tree Detection": S1

Supplementary Materials S1

This appendix contains images of predicted plots for all sites. Models were trained from training data from the same site. These plots form the basis of the analysis for the precision recall curves found in the main article (Figure 4).

Eastern Deciduous p2-10
Alpine p11-22
Oak Woodland: p23-55
Mixed Pine: p55-73

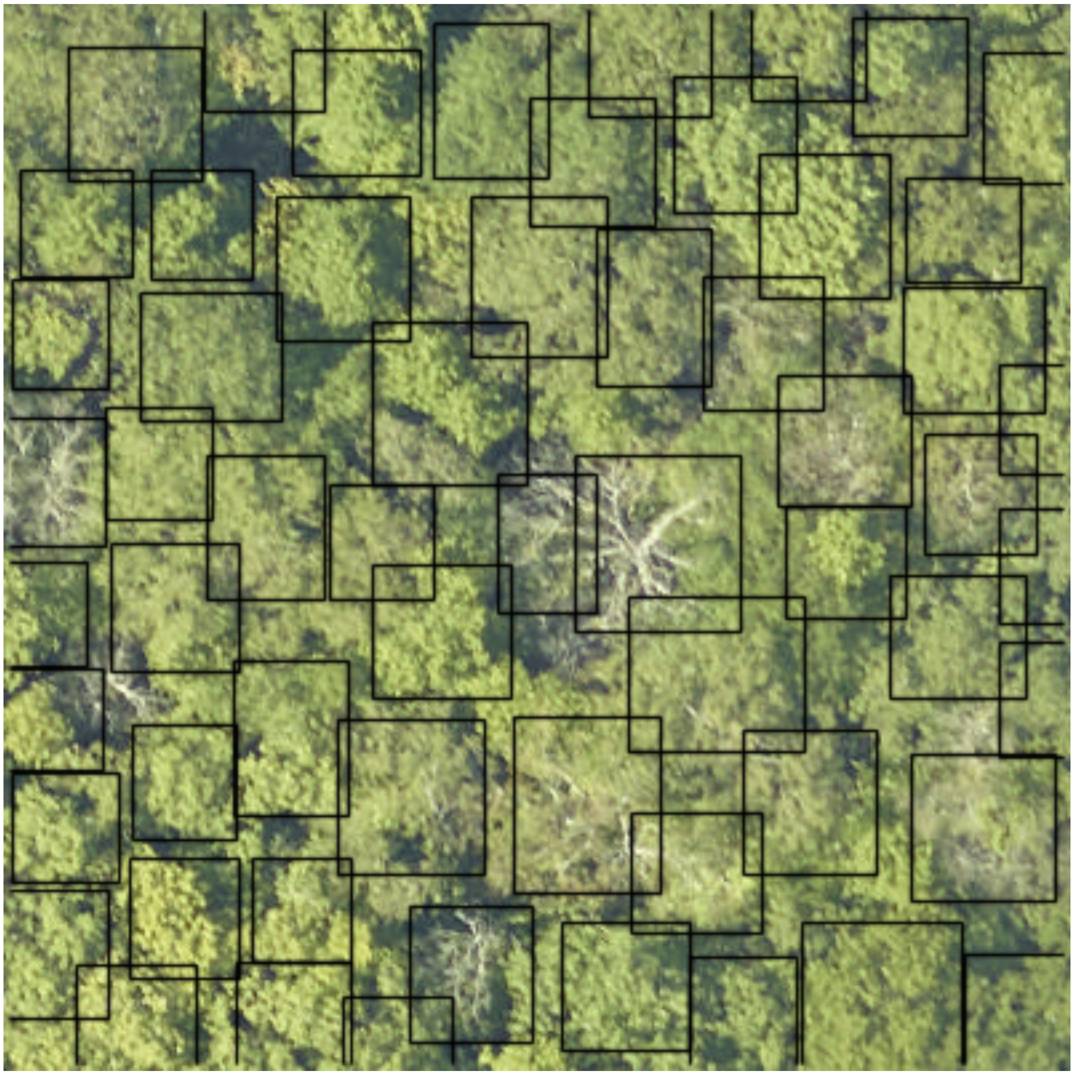

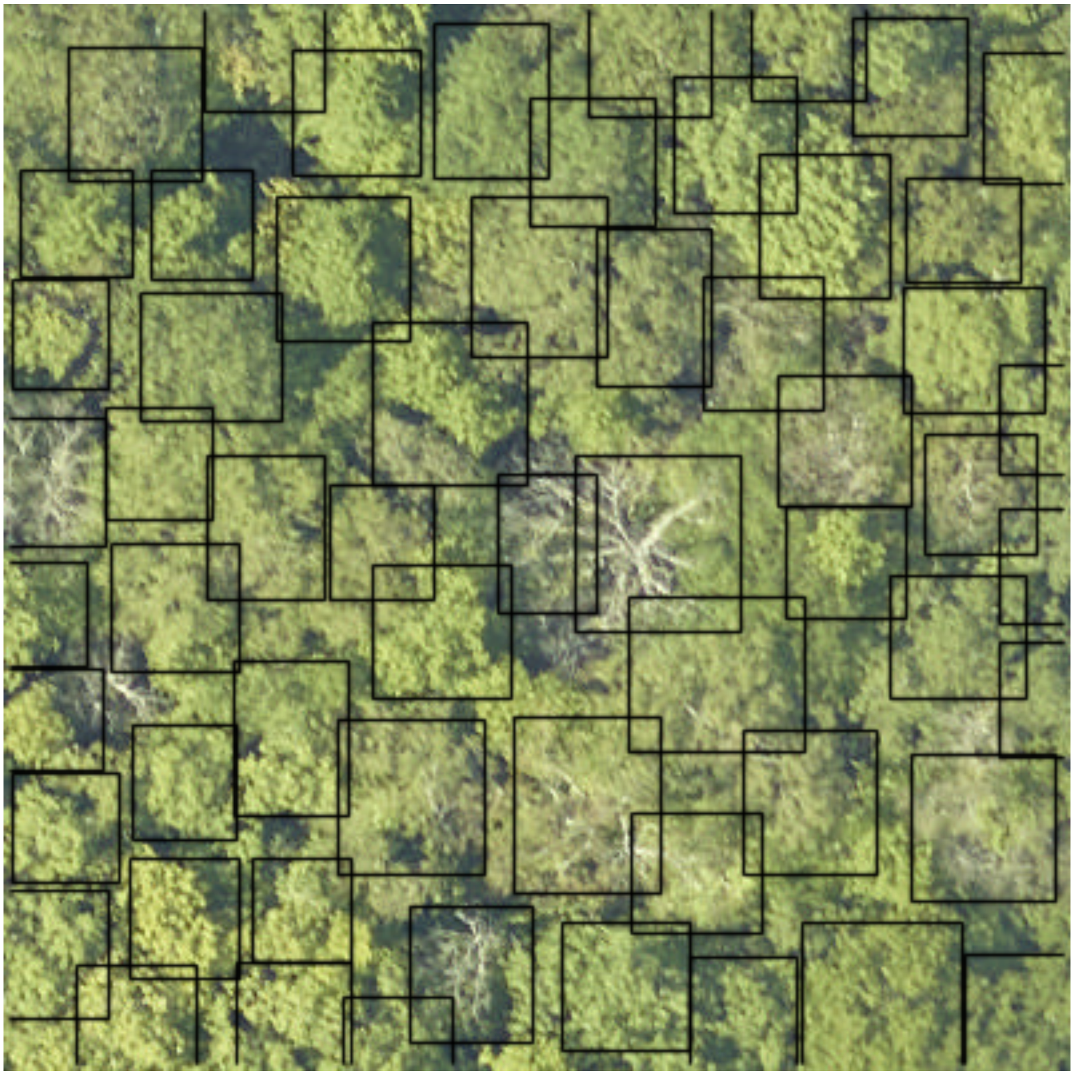

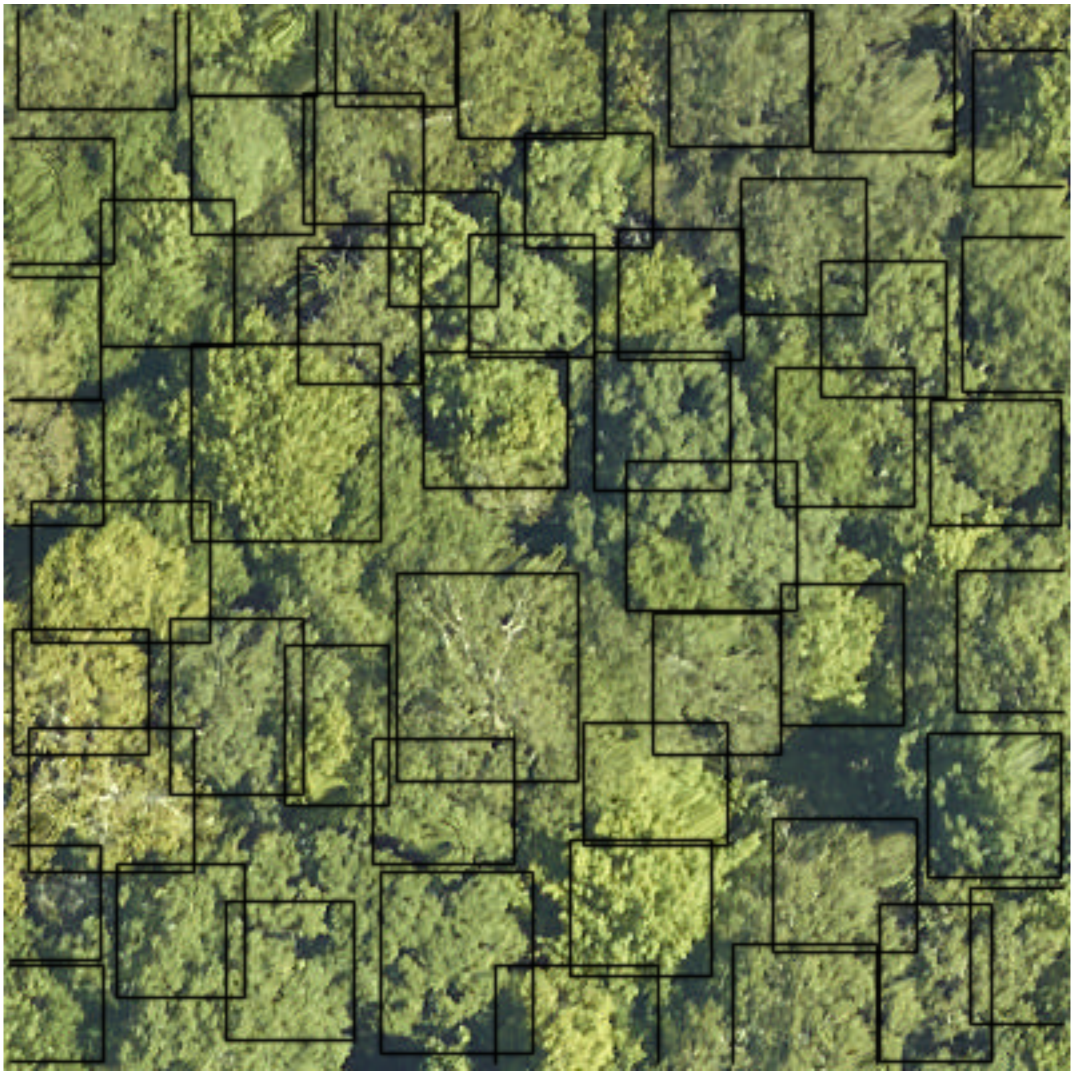

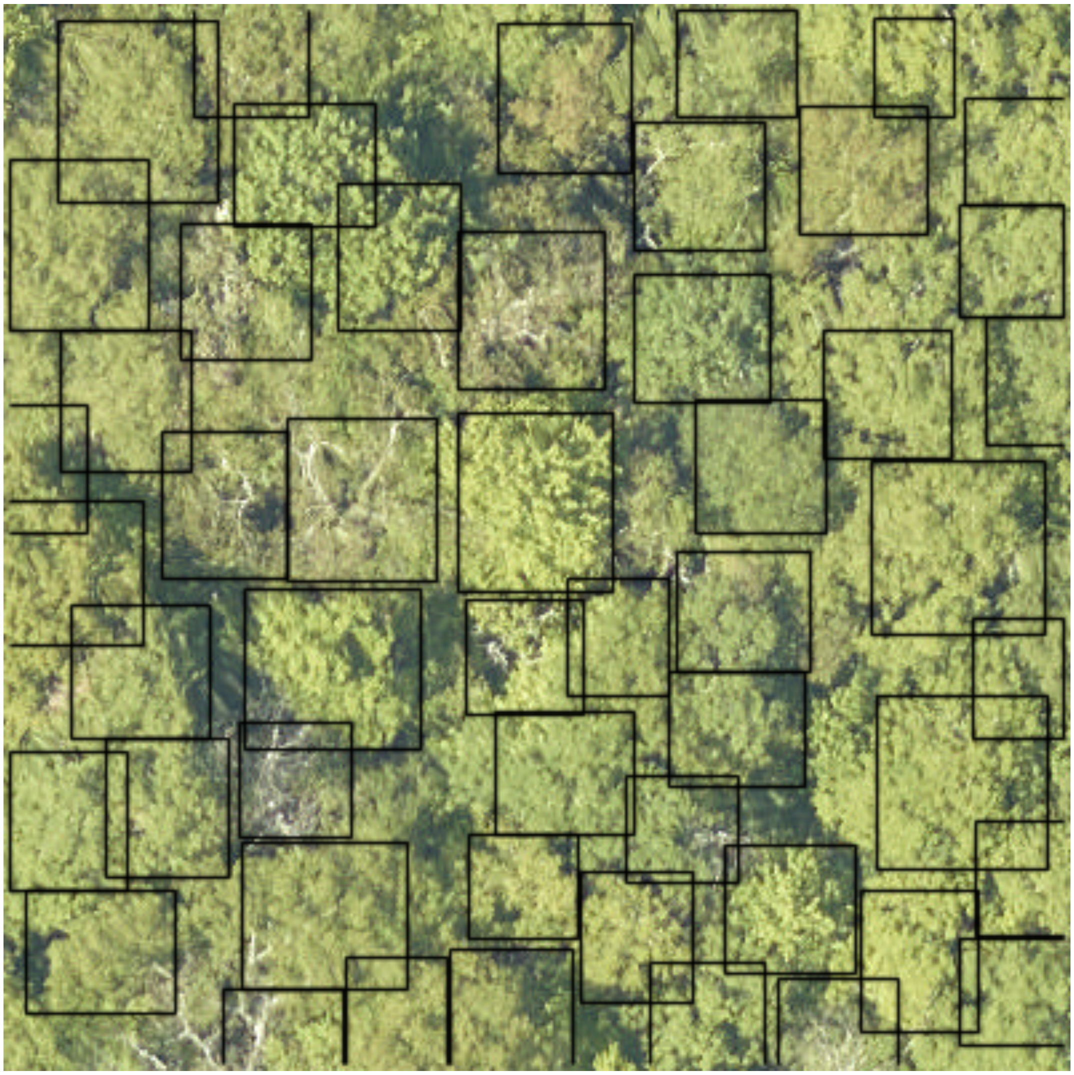

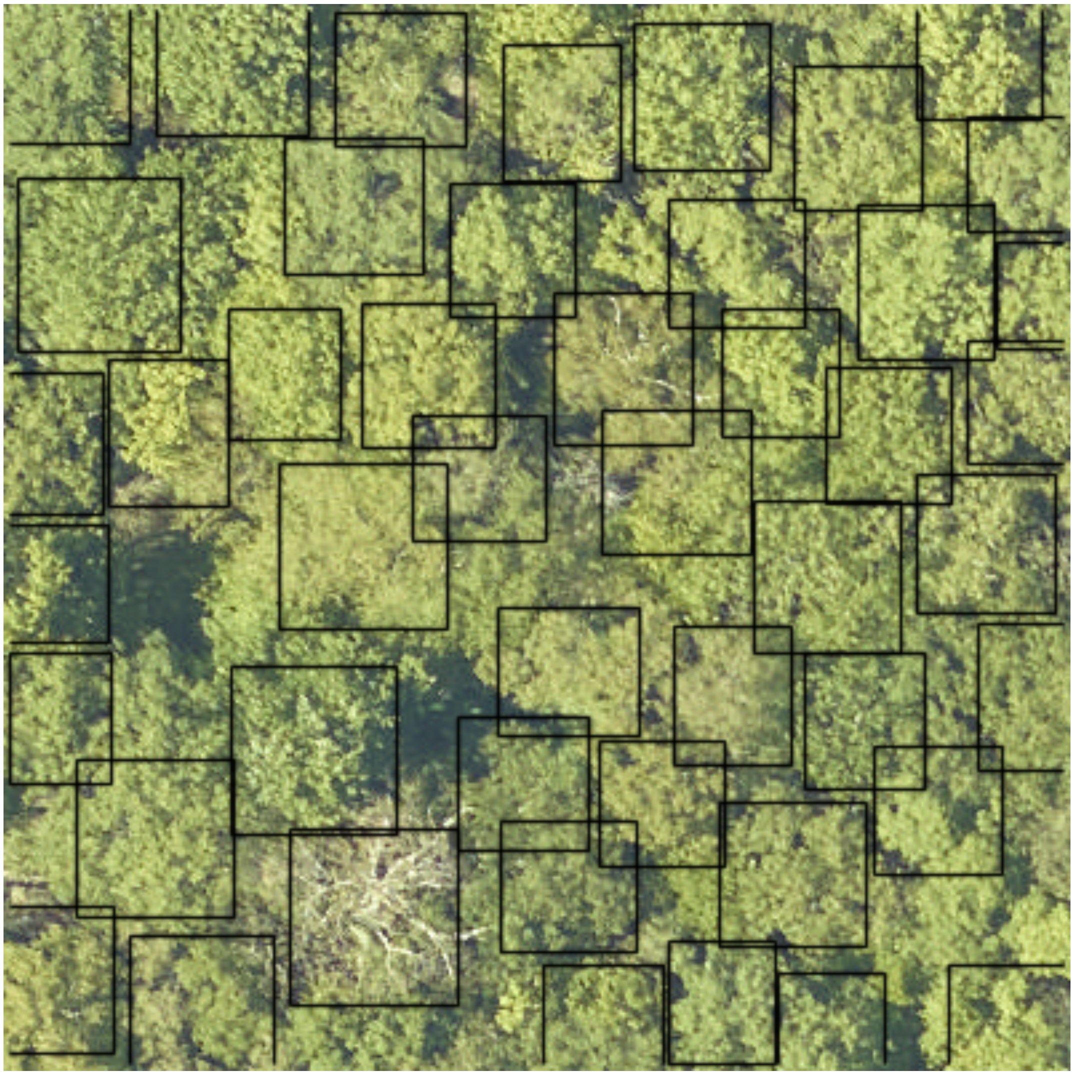

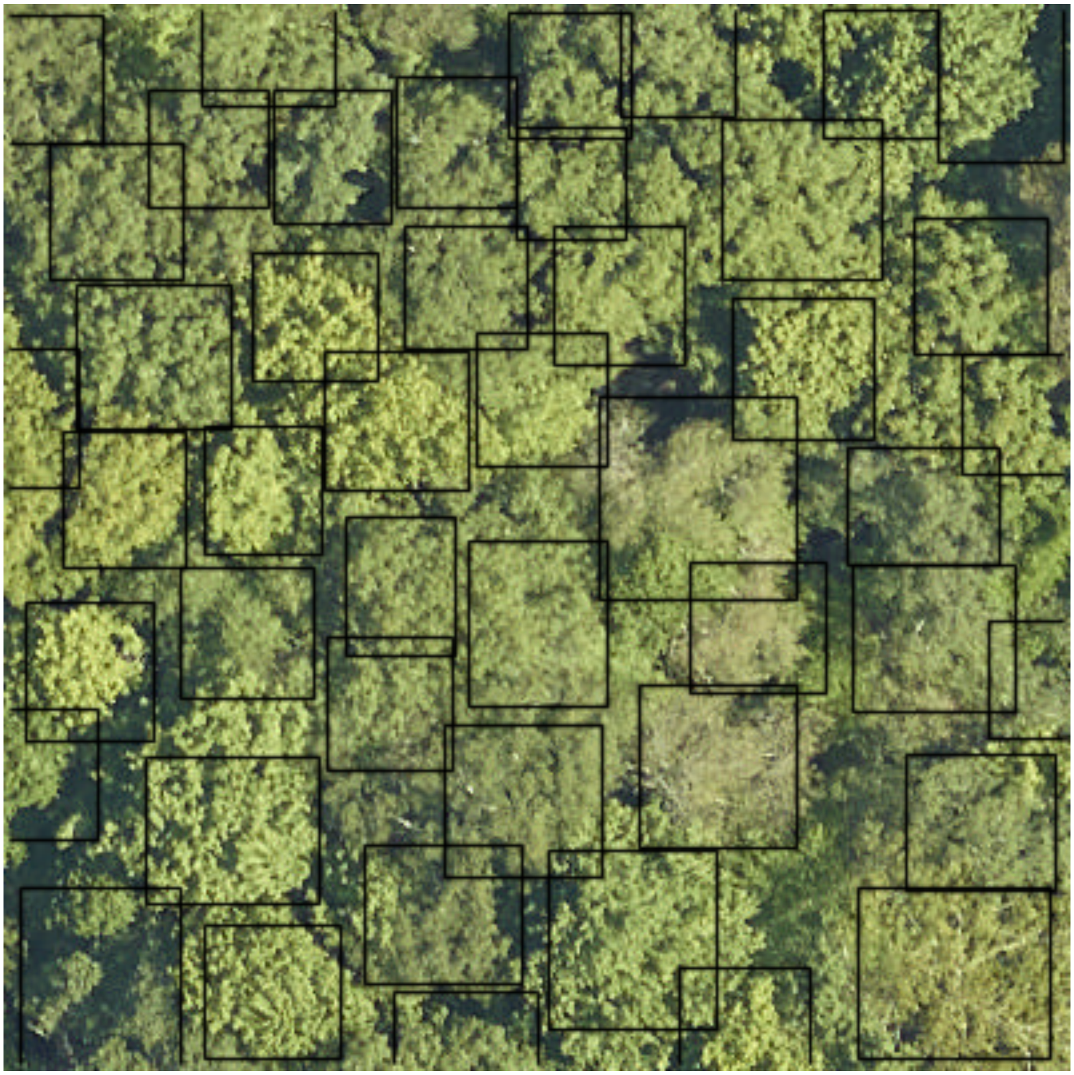

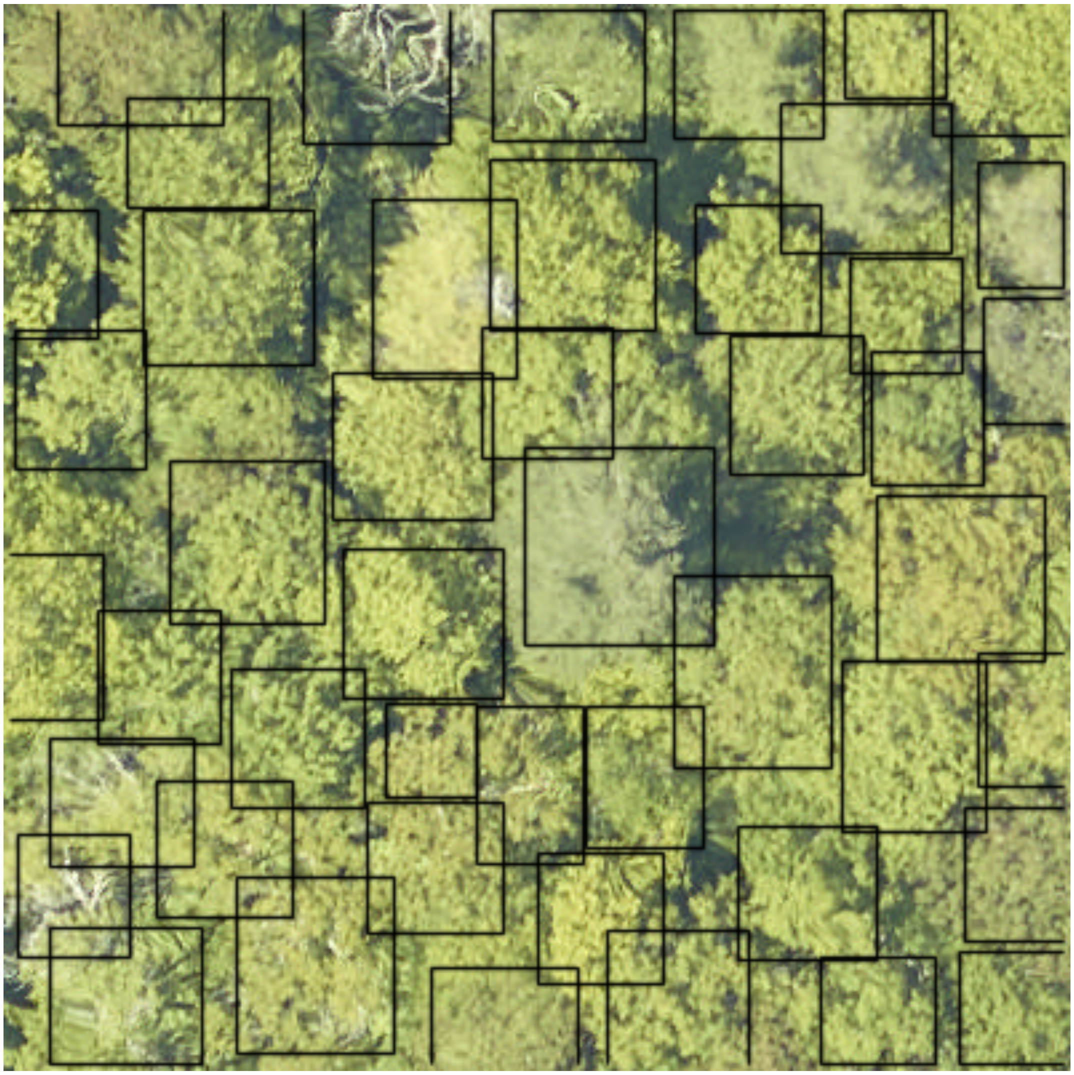

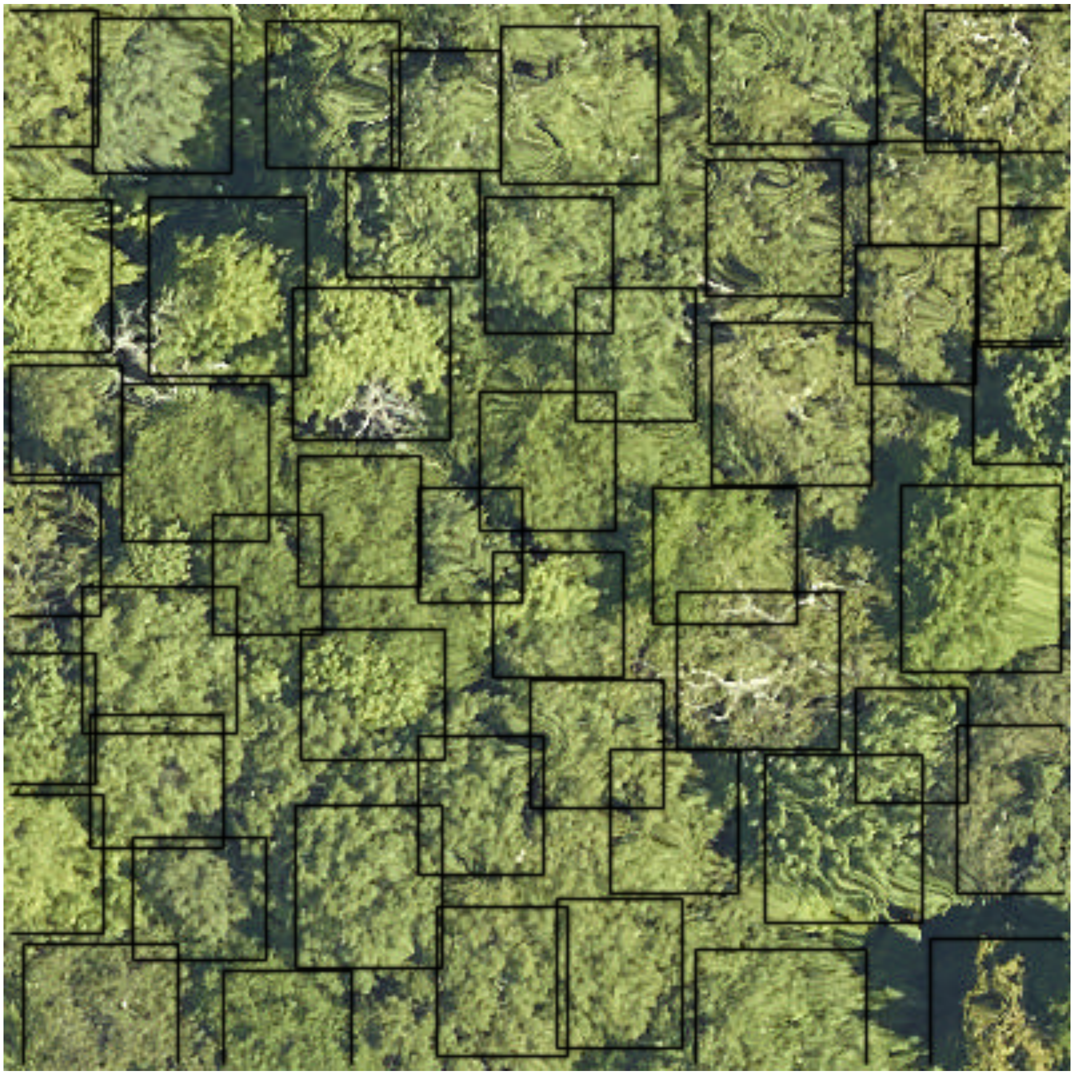

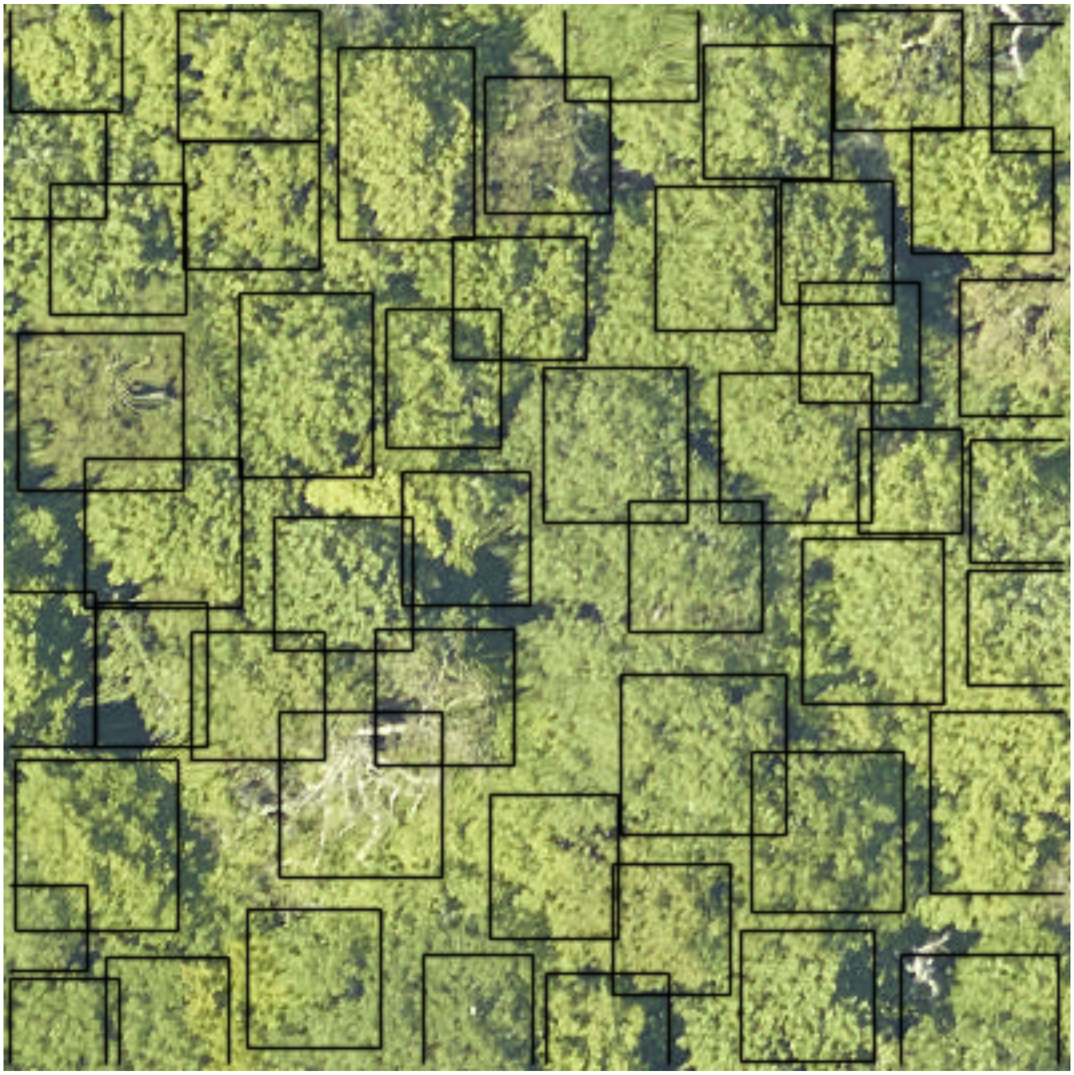

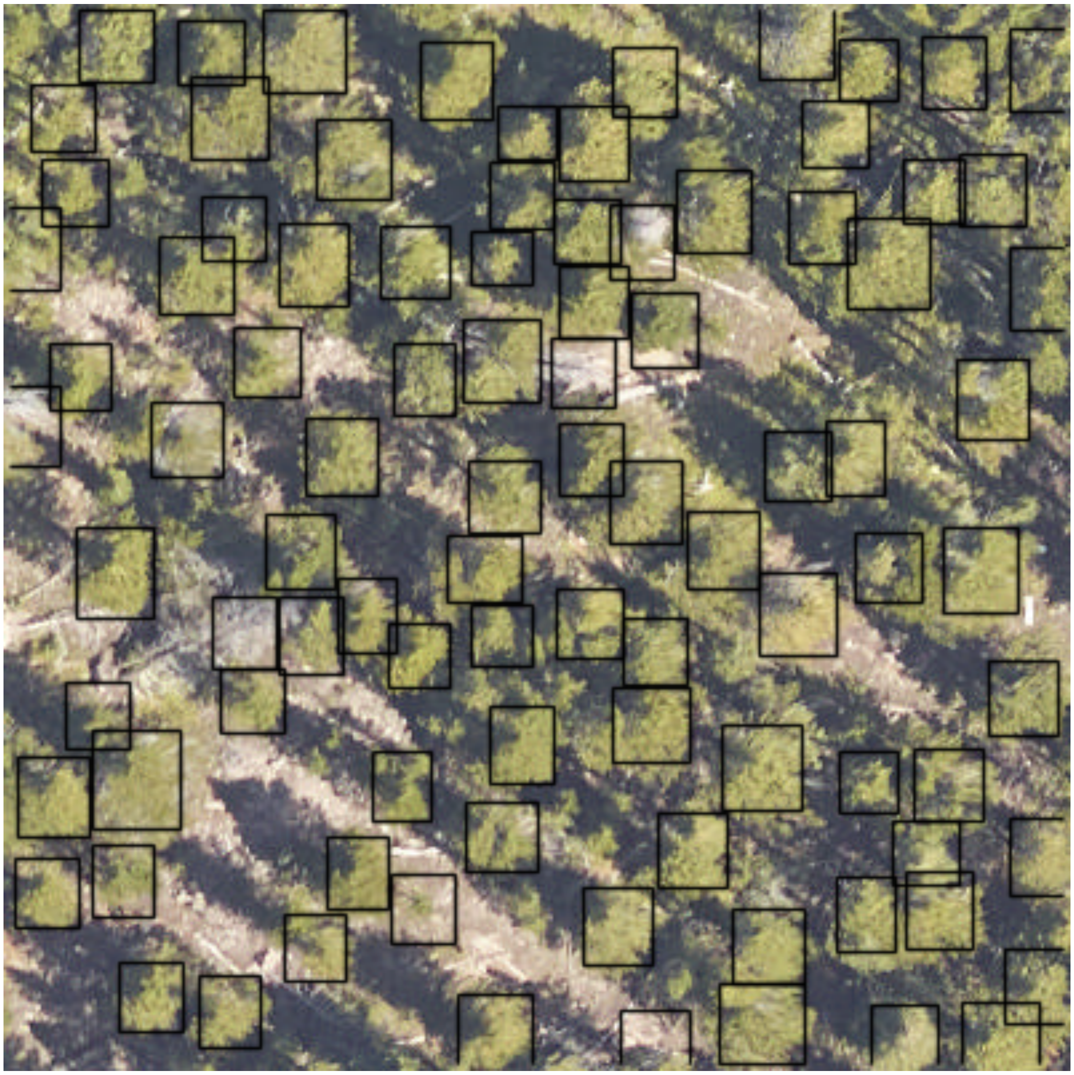

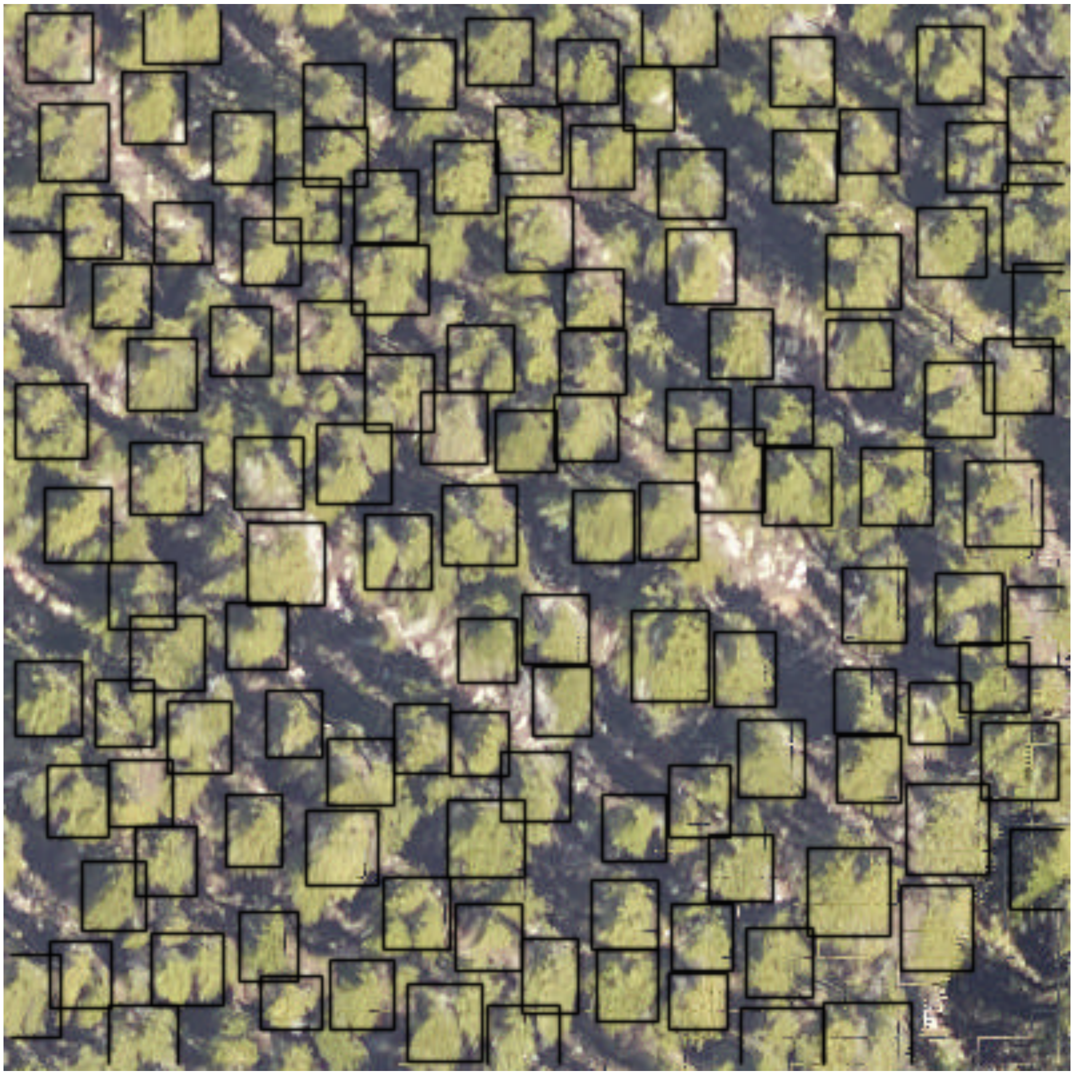

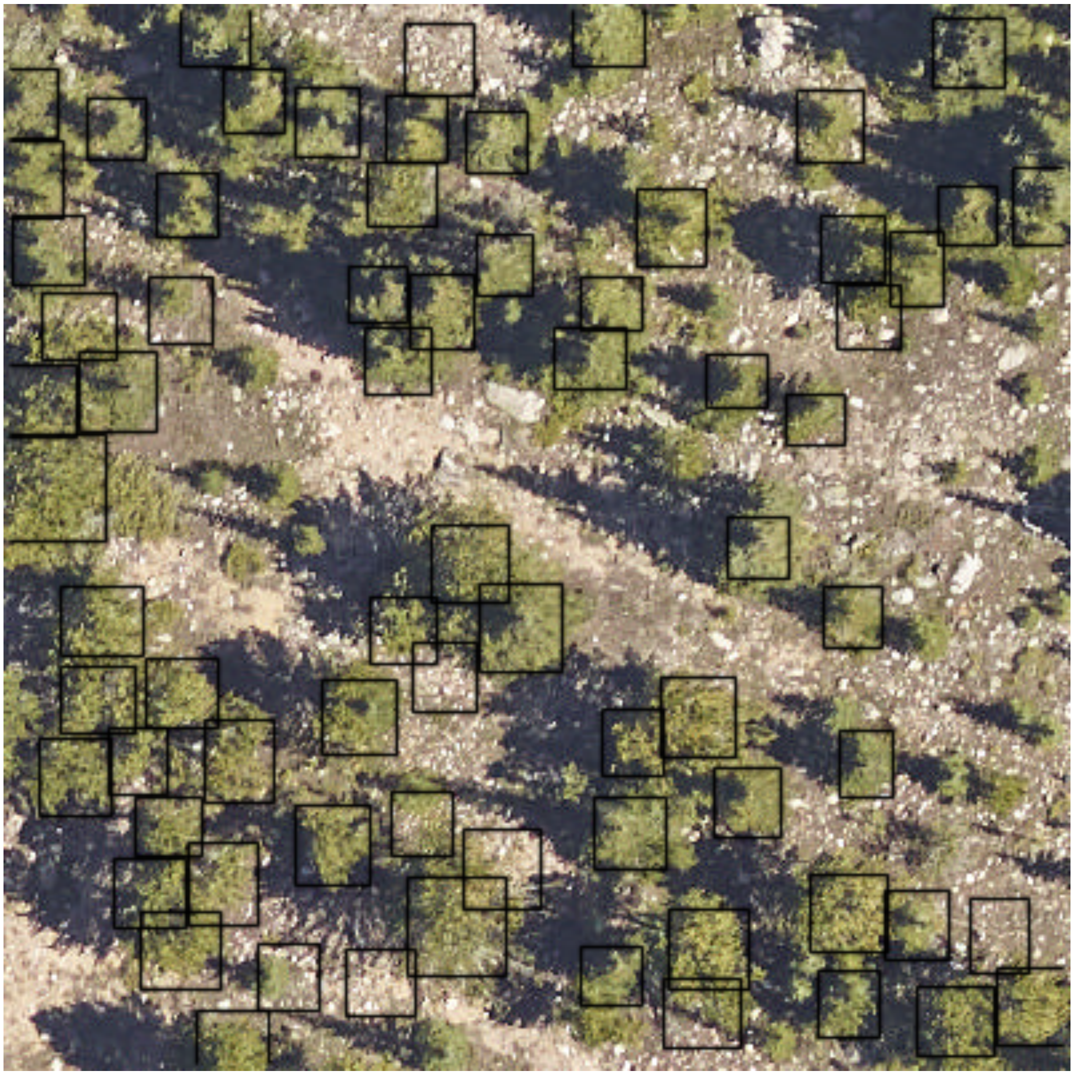

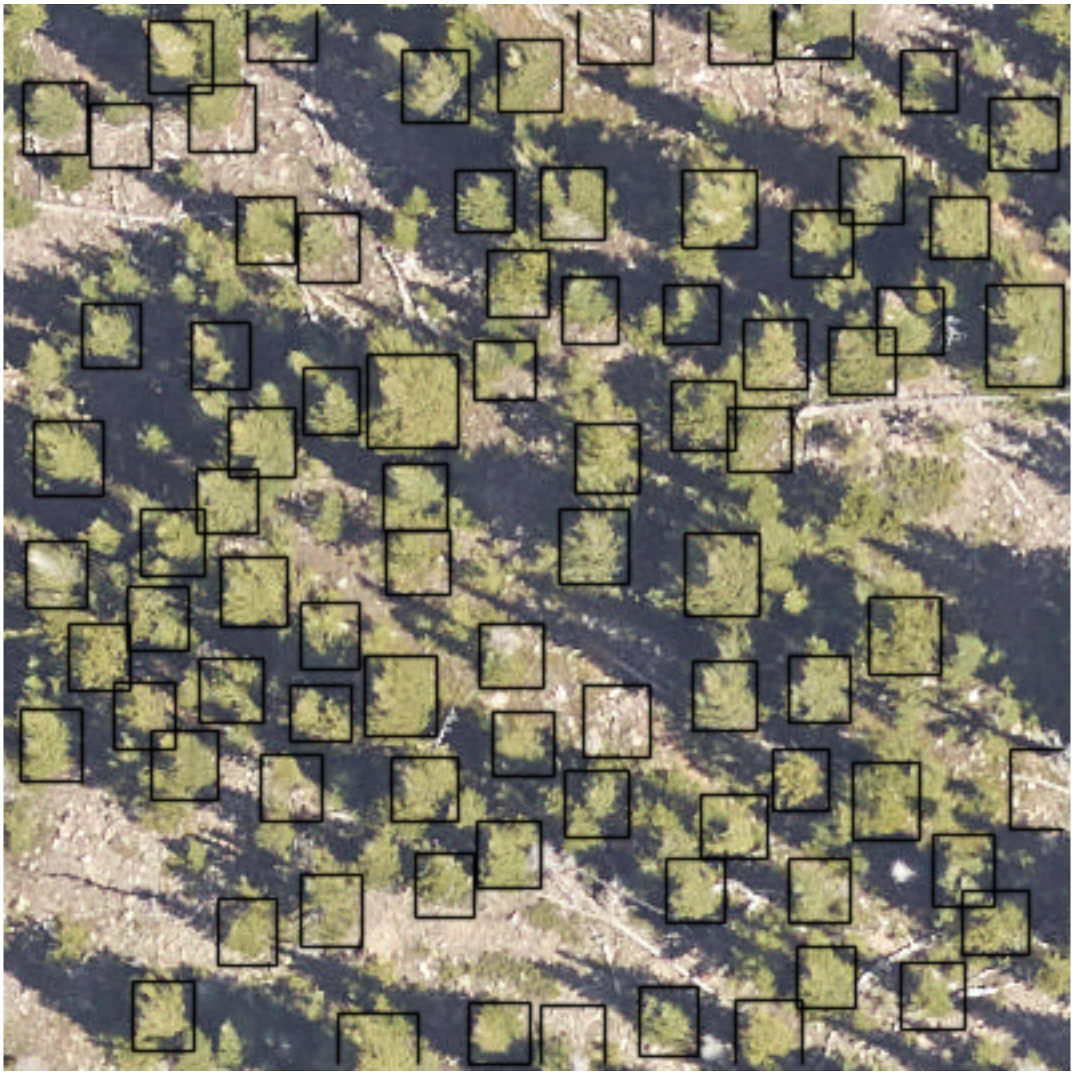

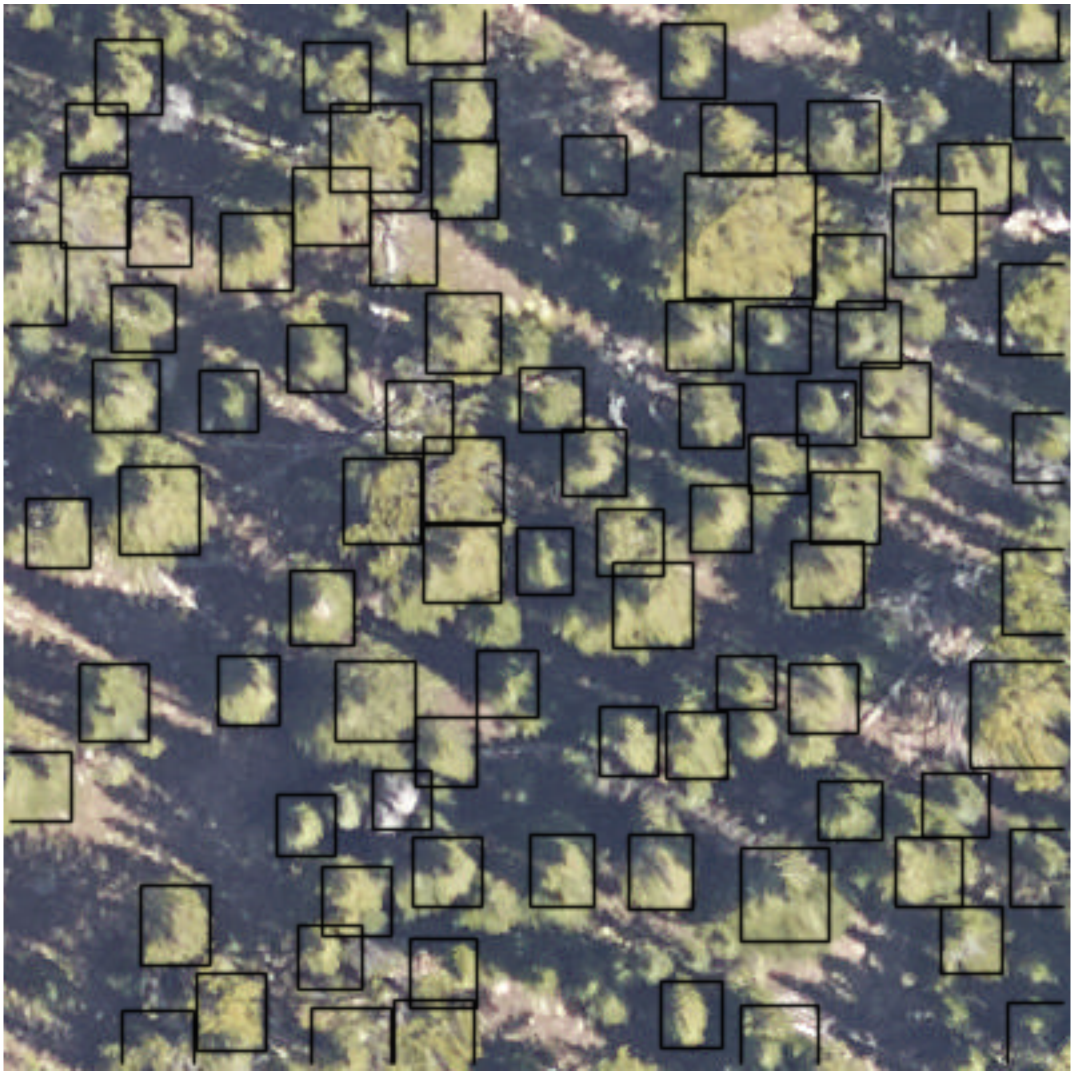

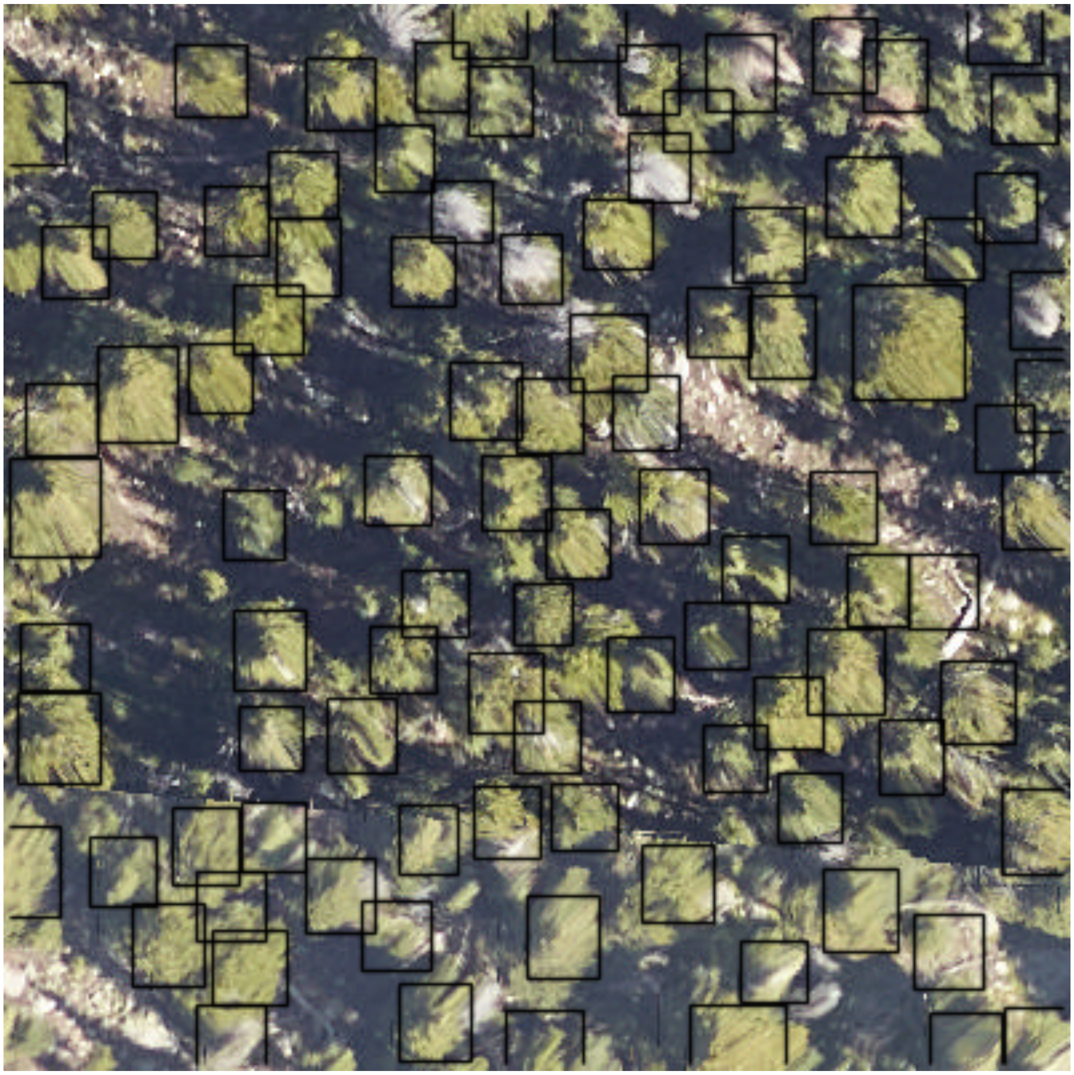

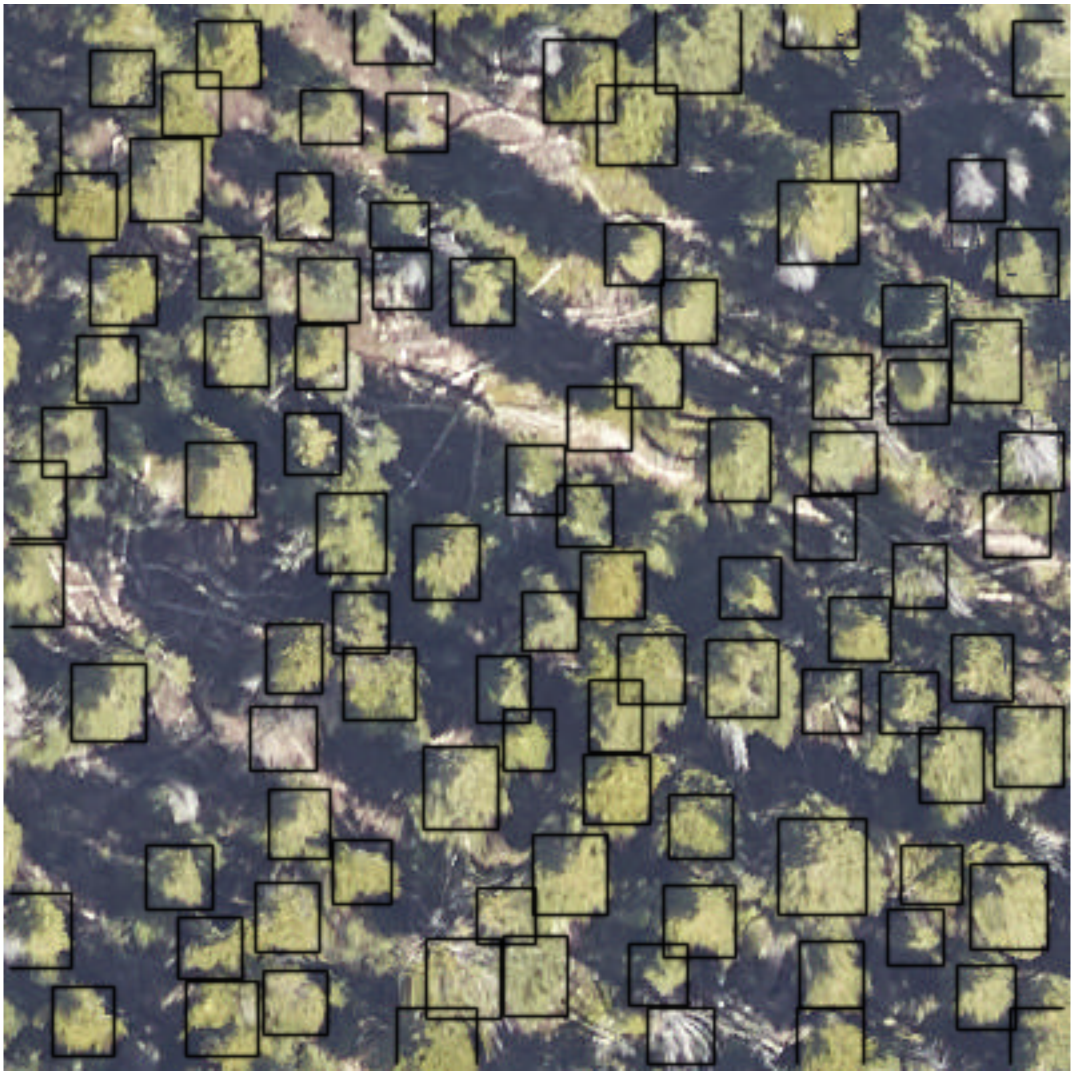

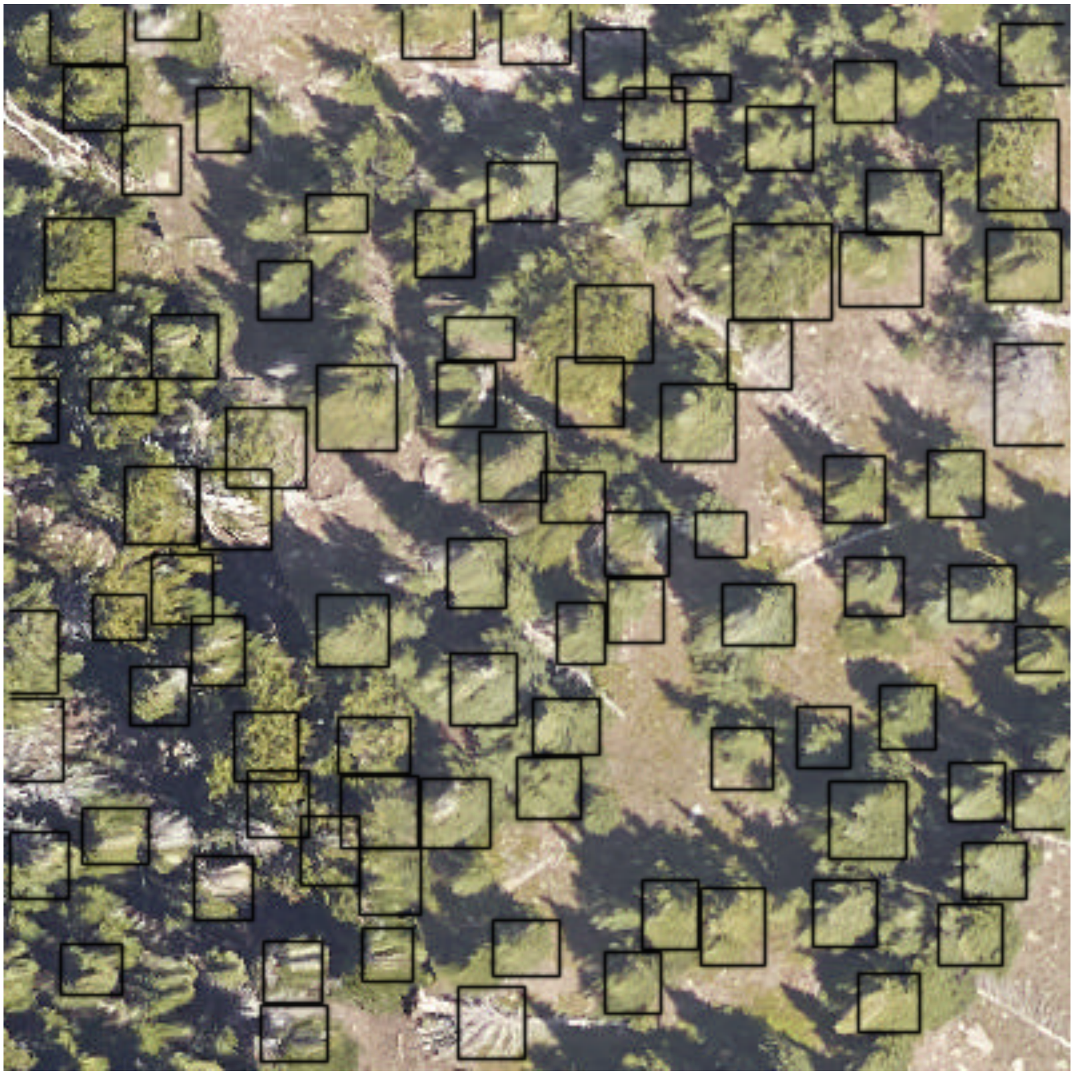

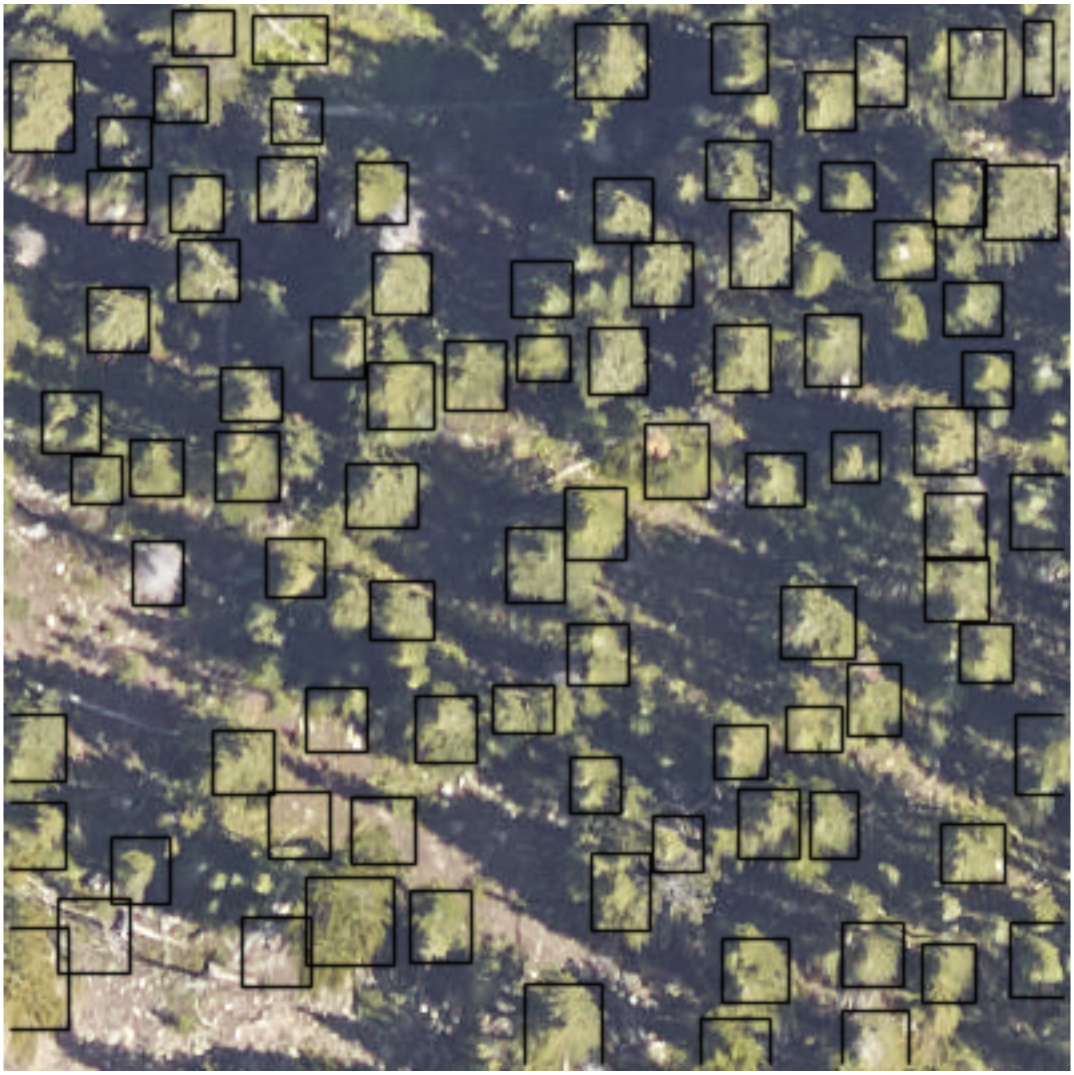

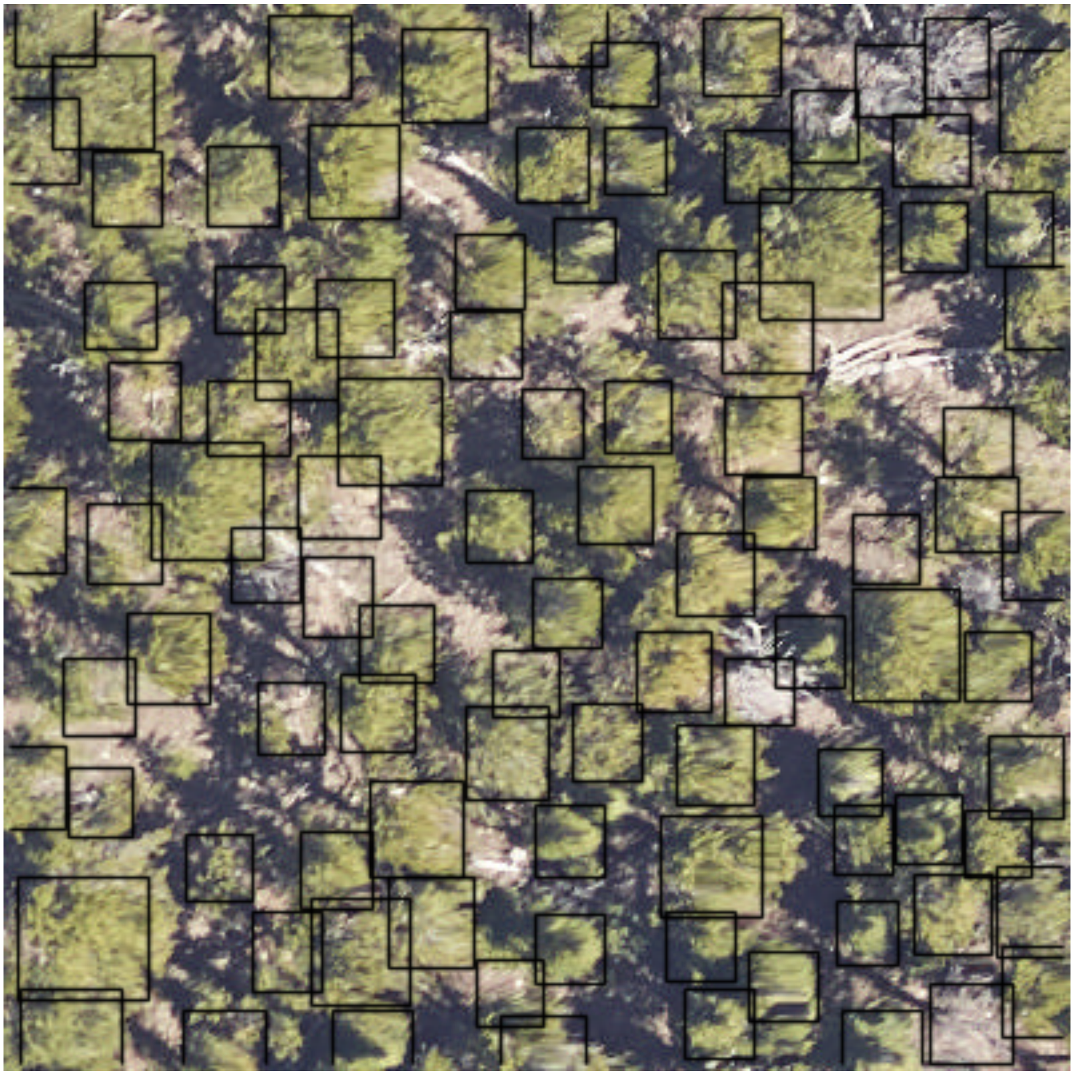

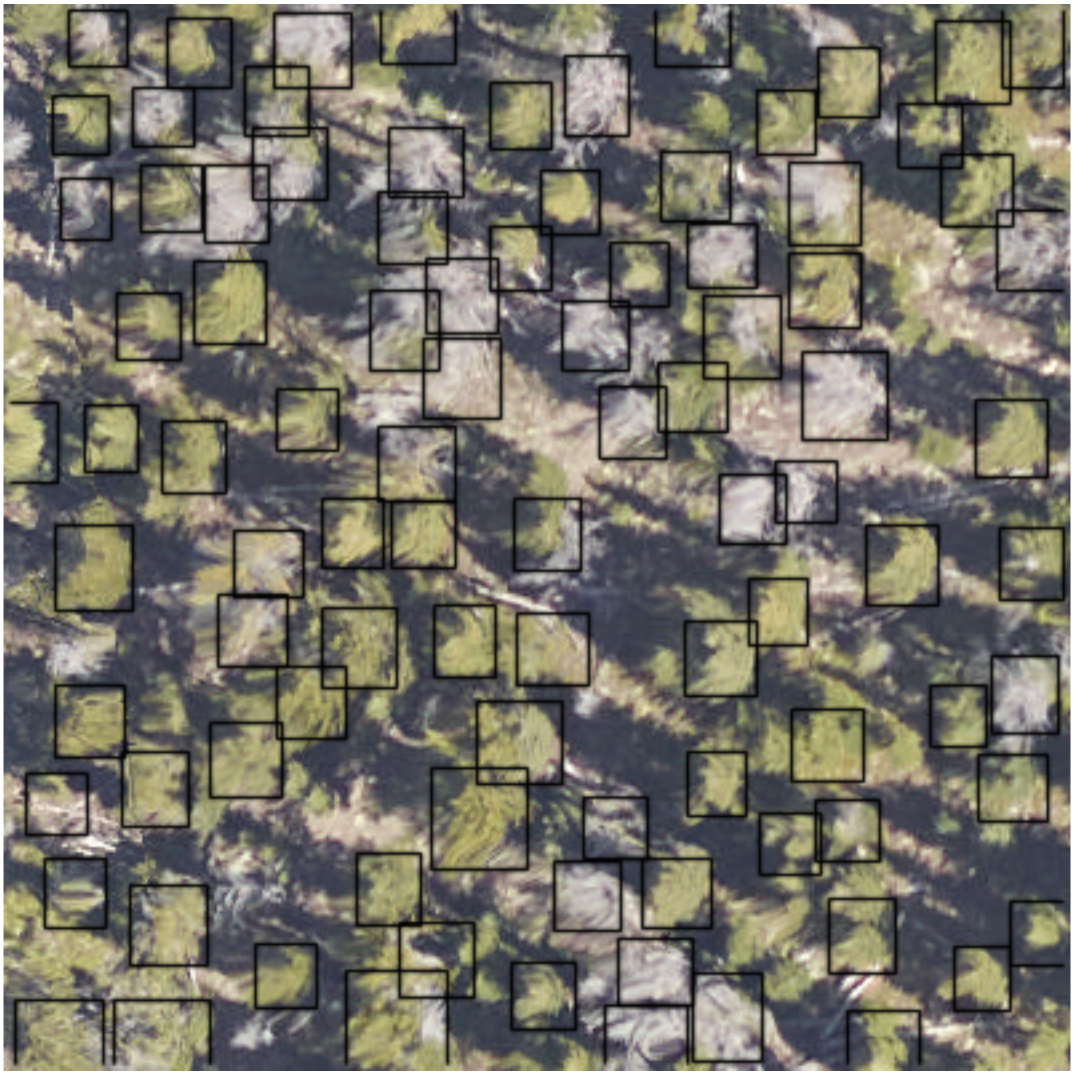

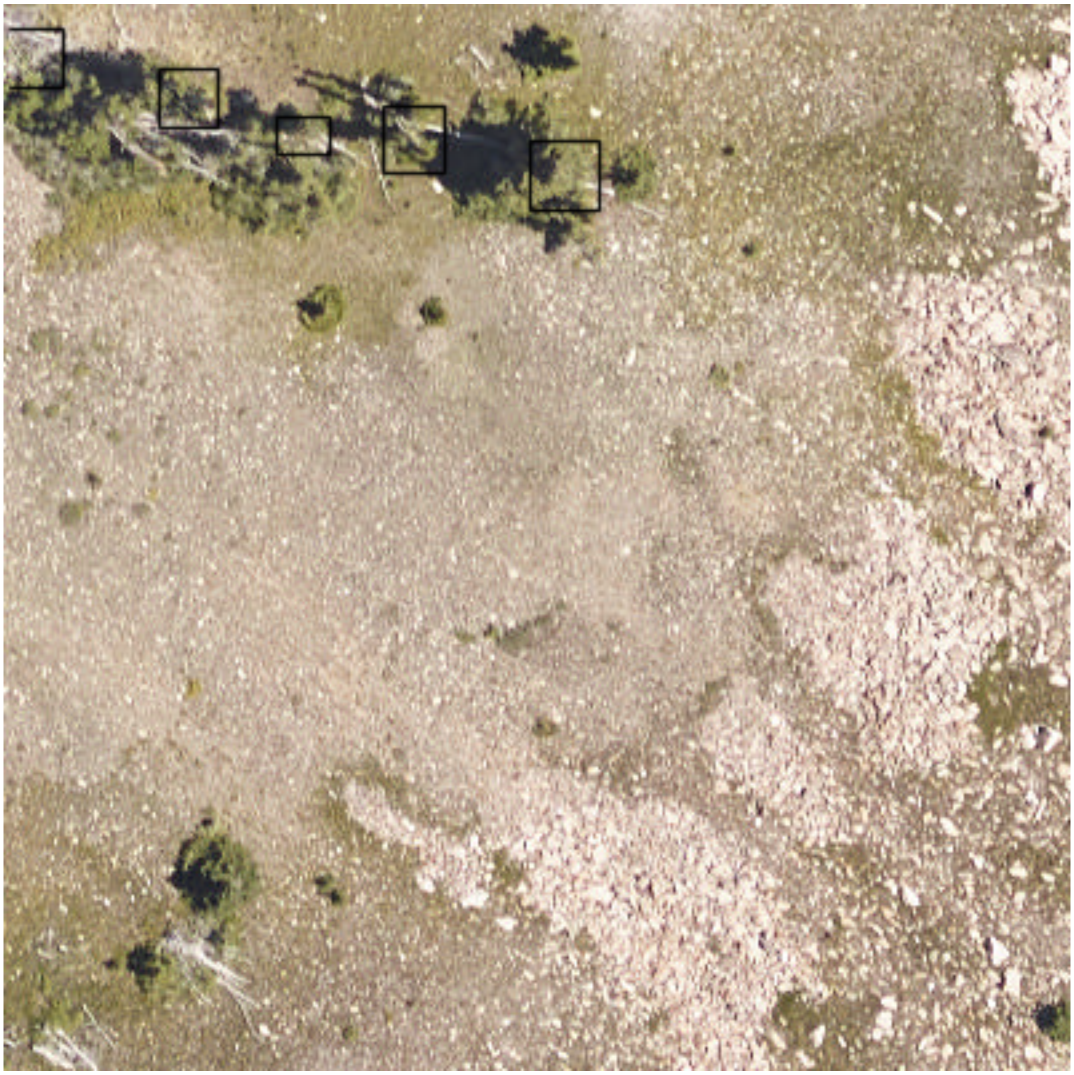

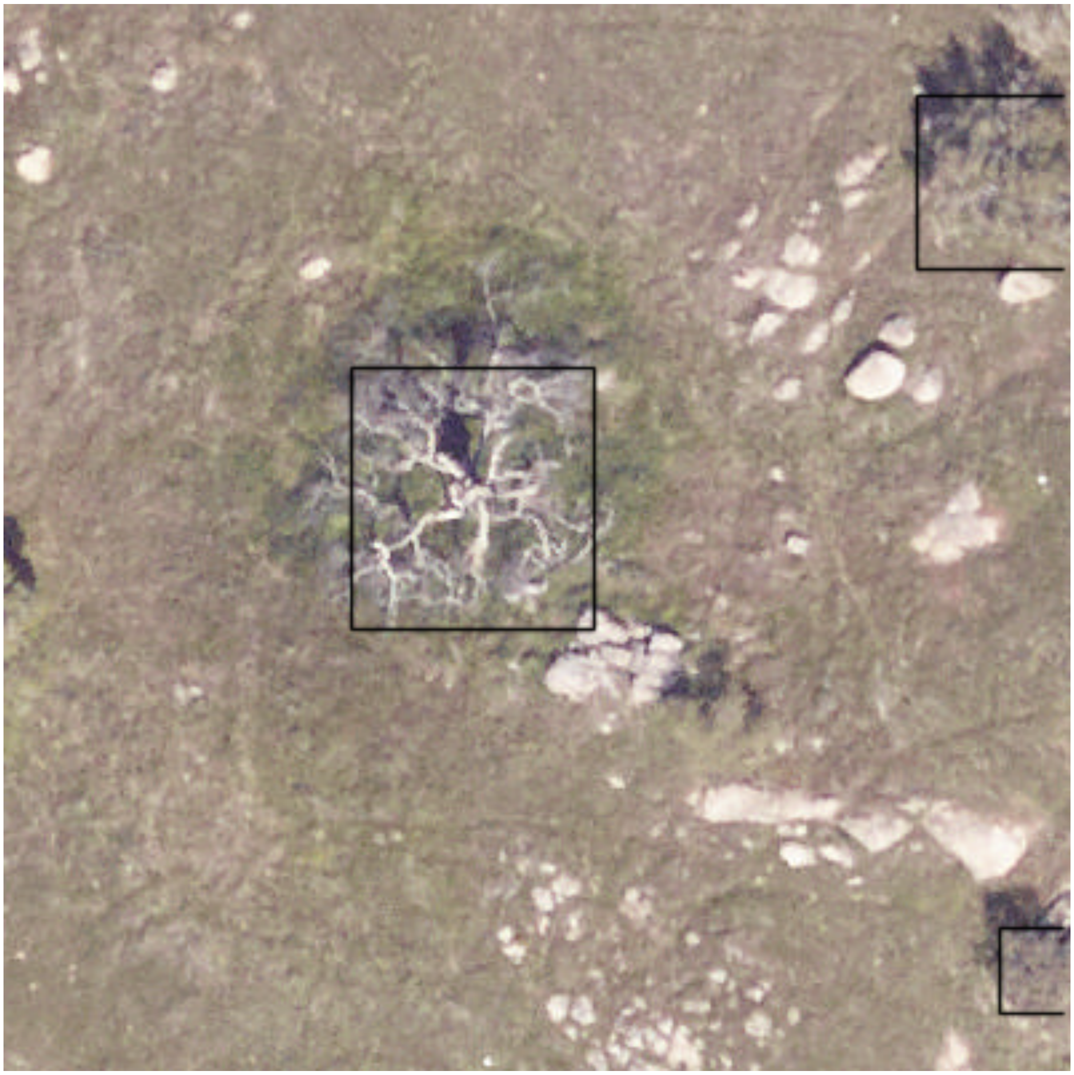

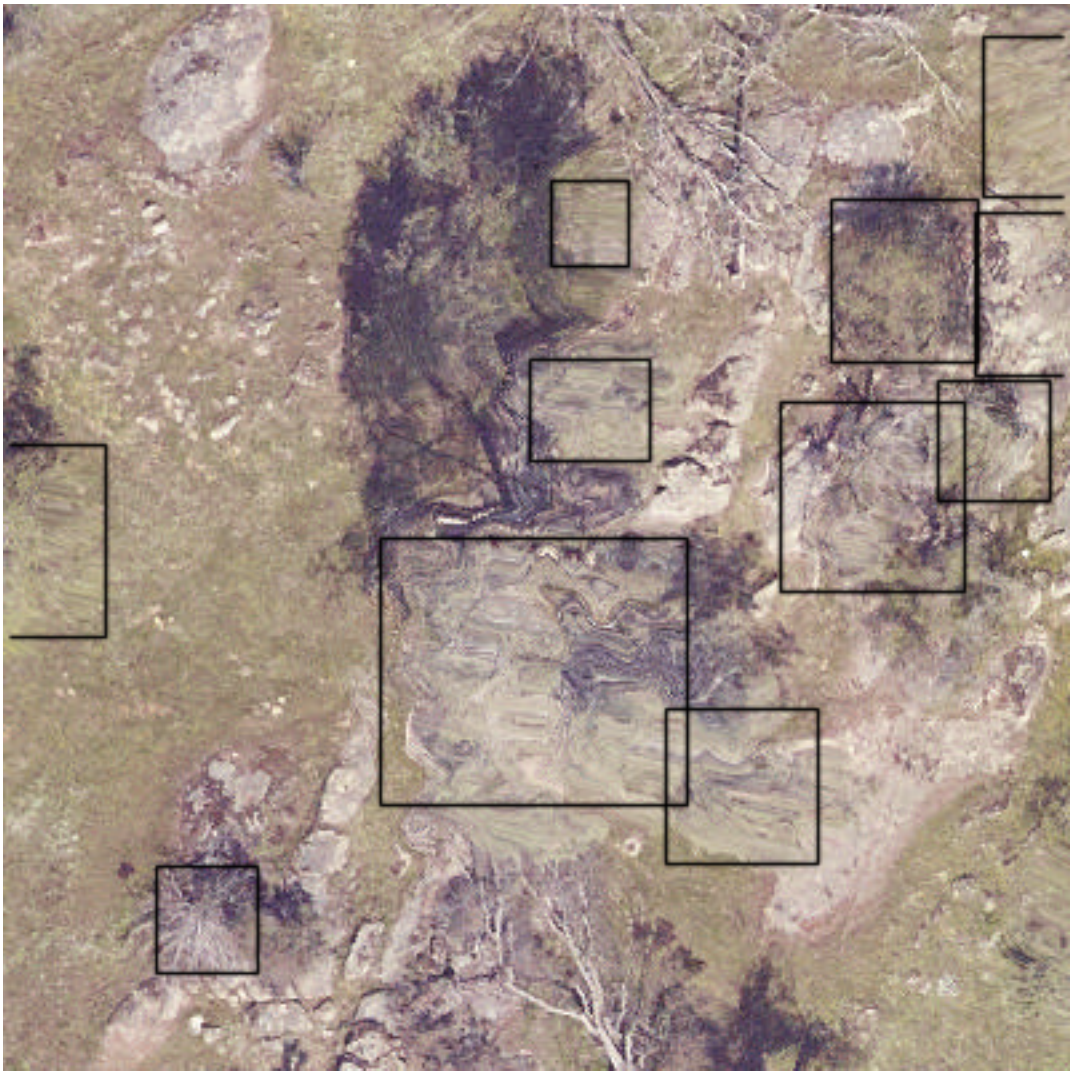

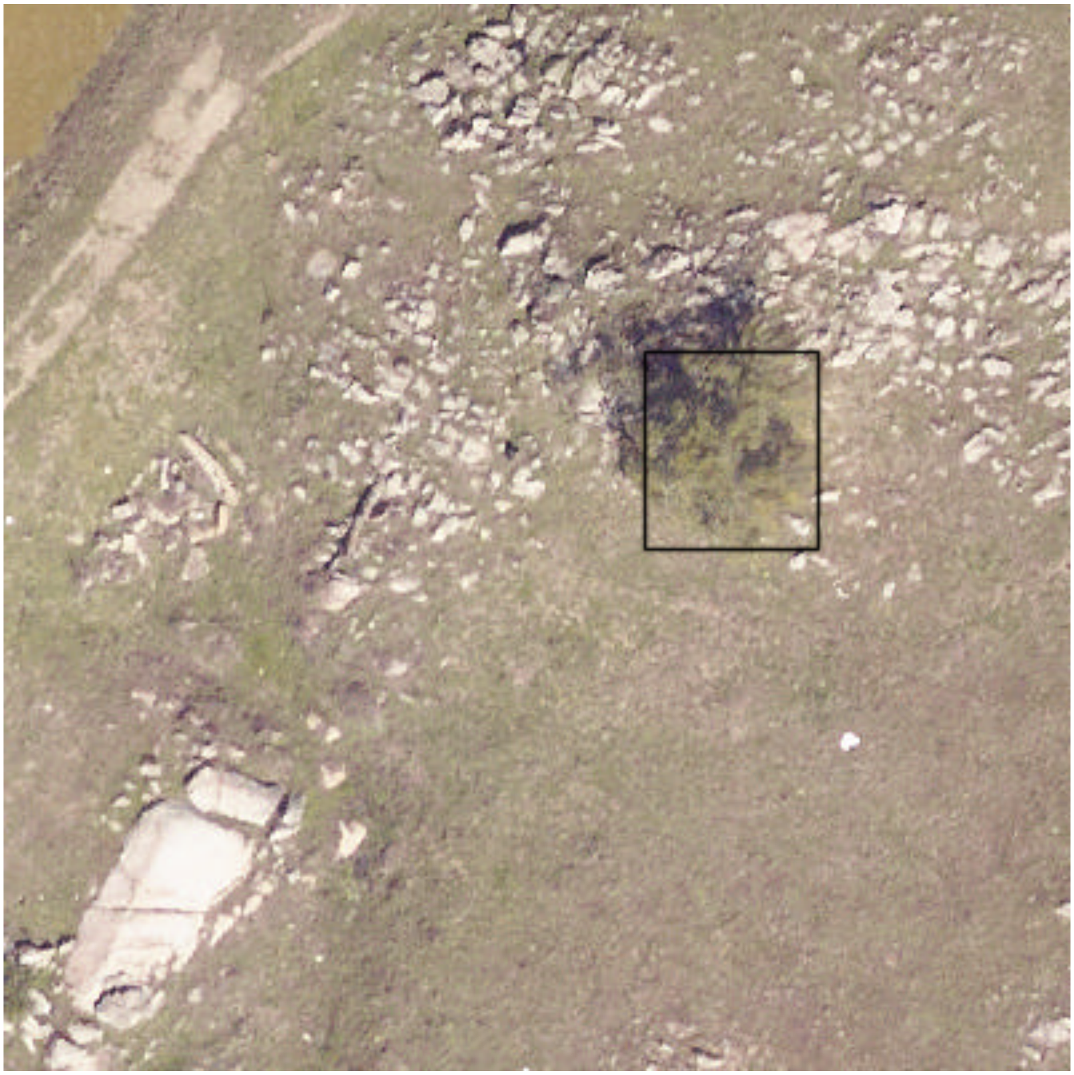

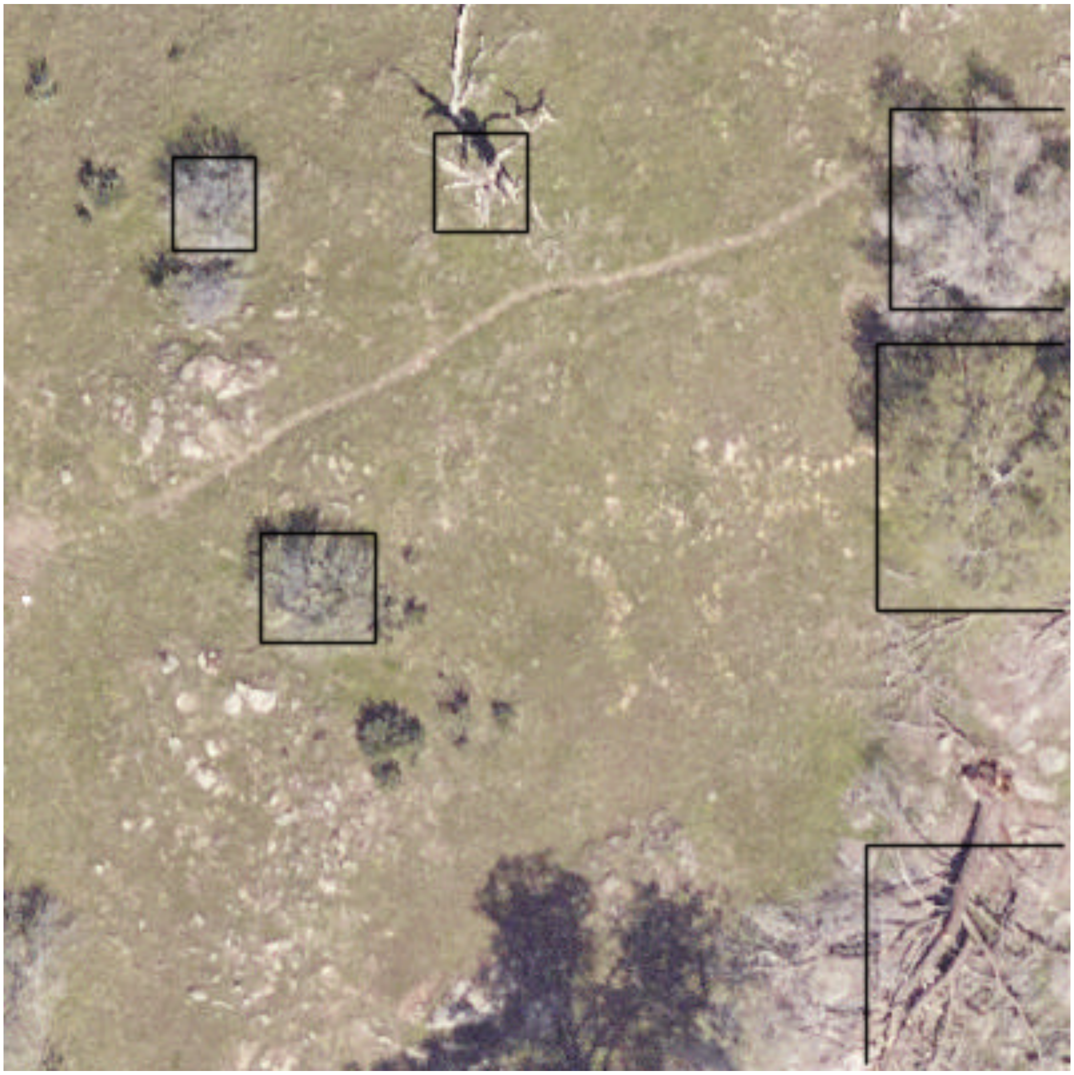

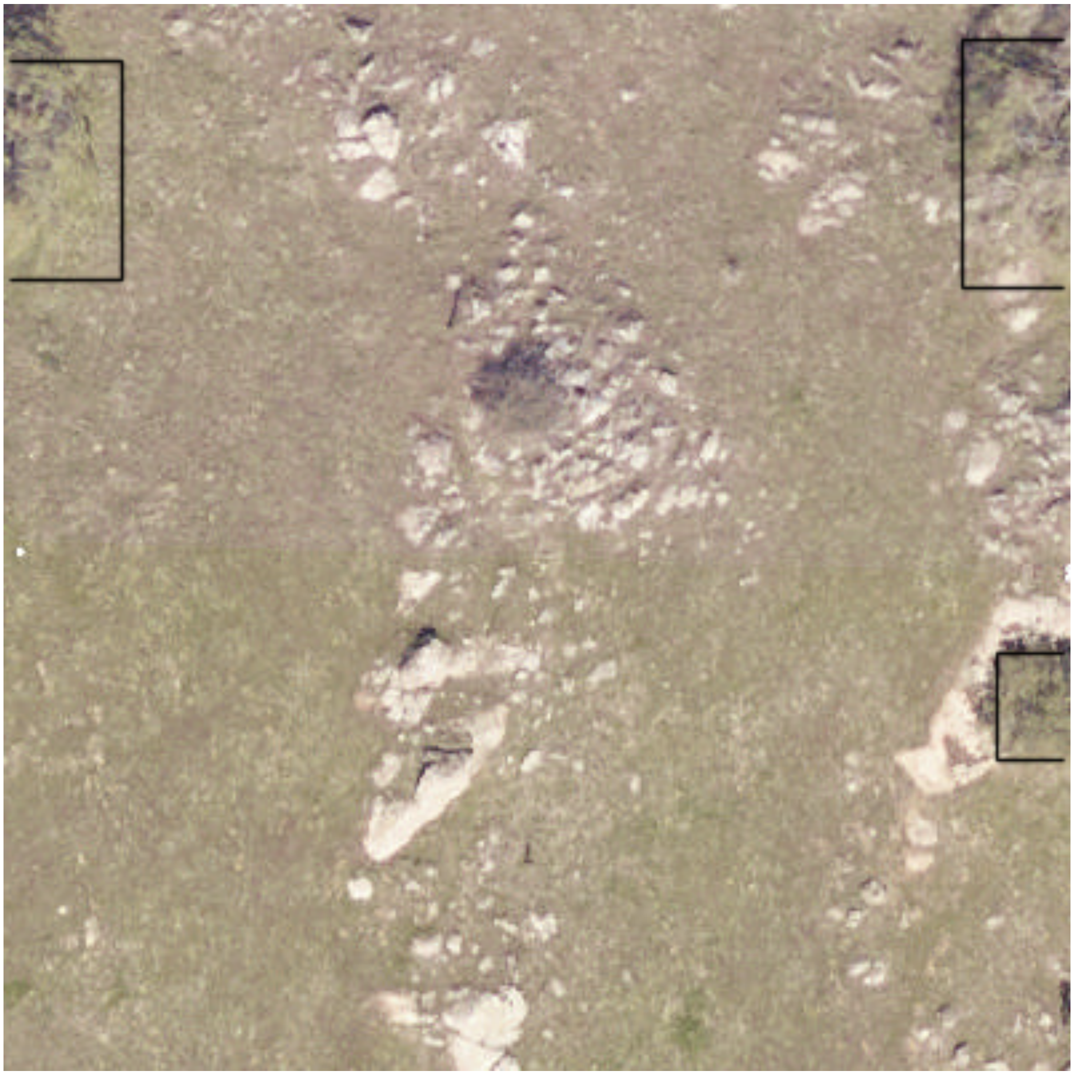

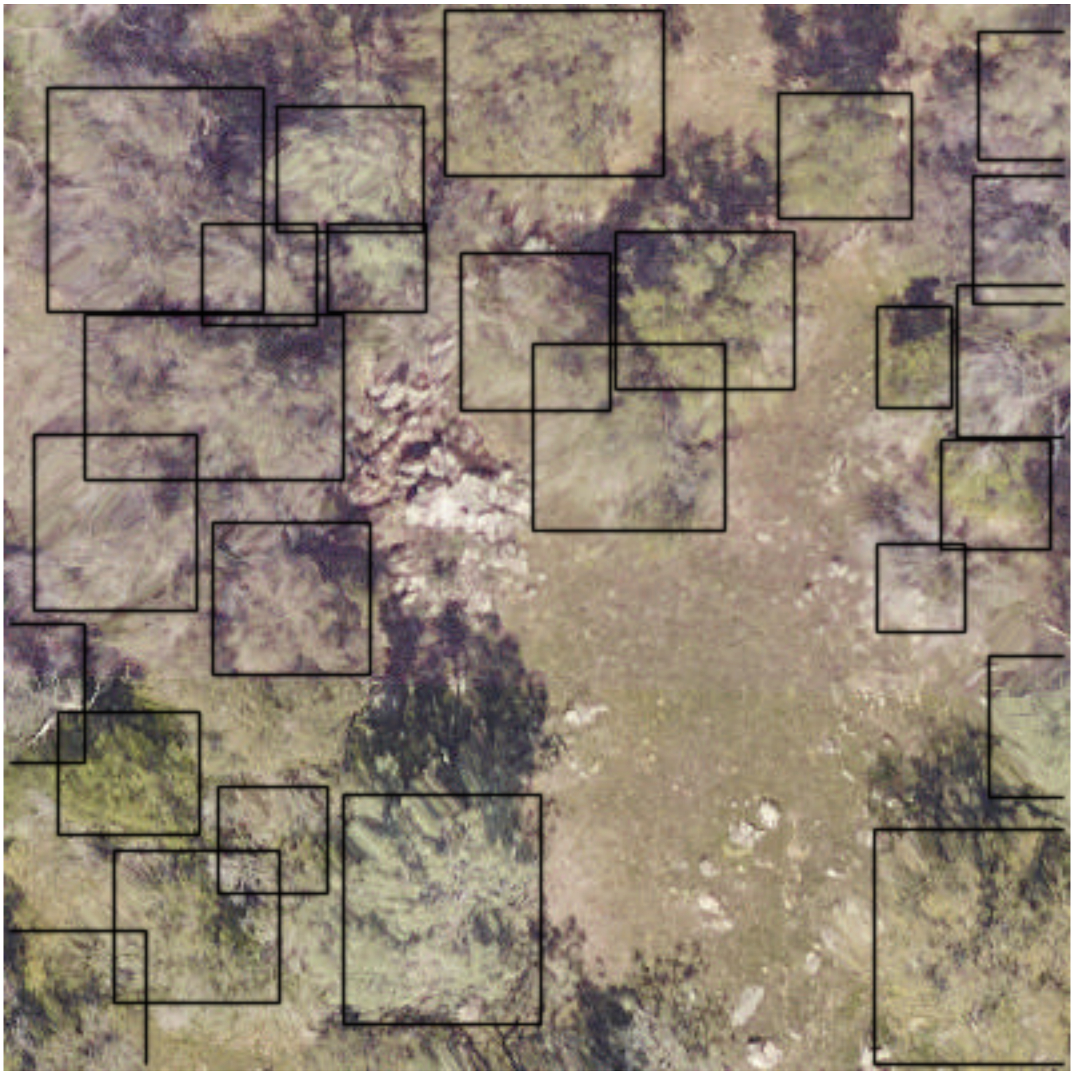

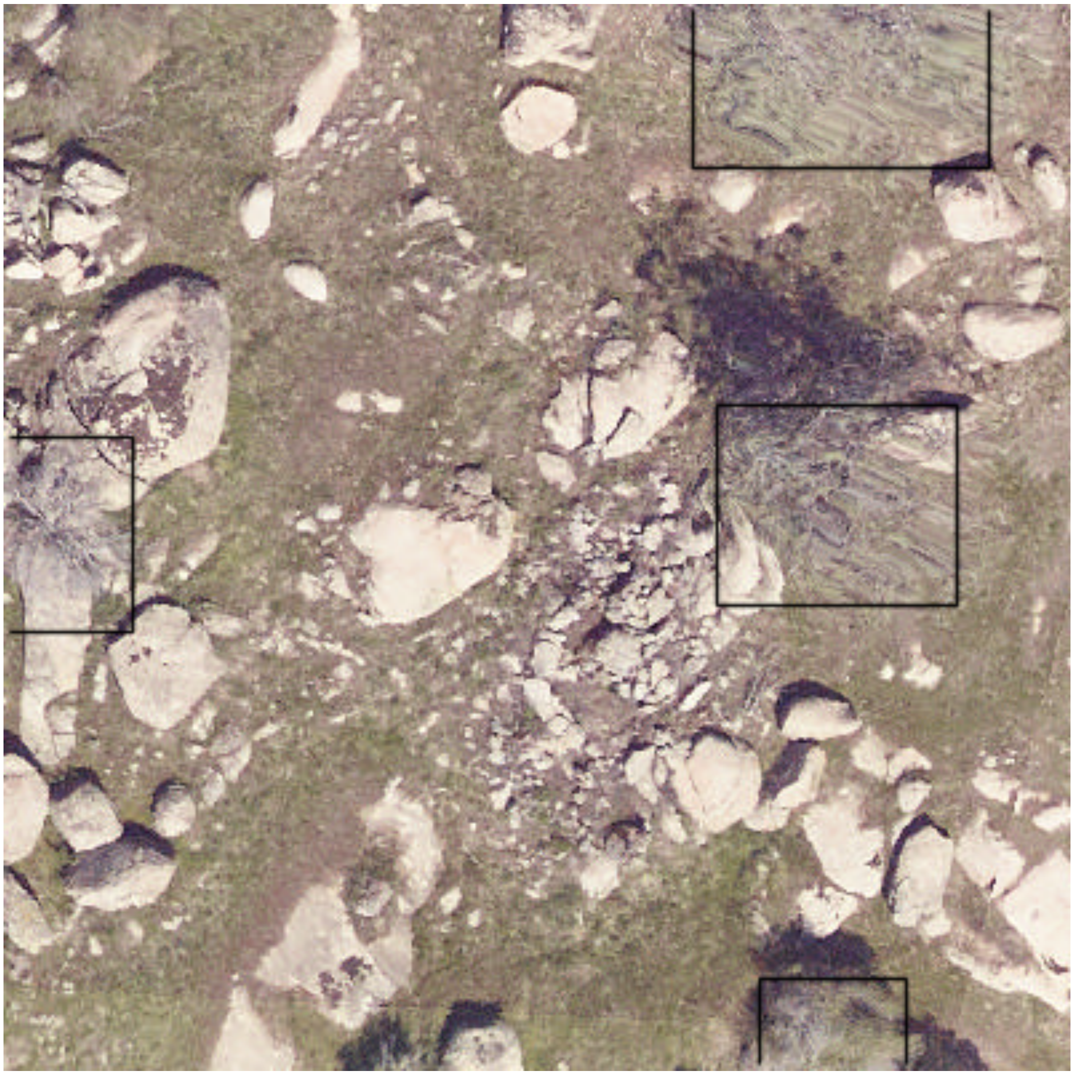

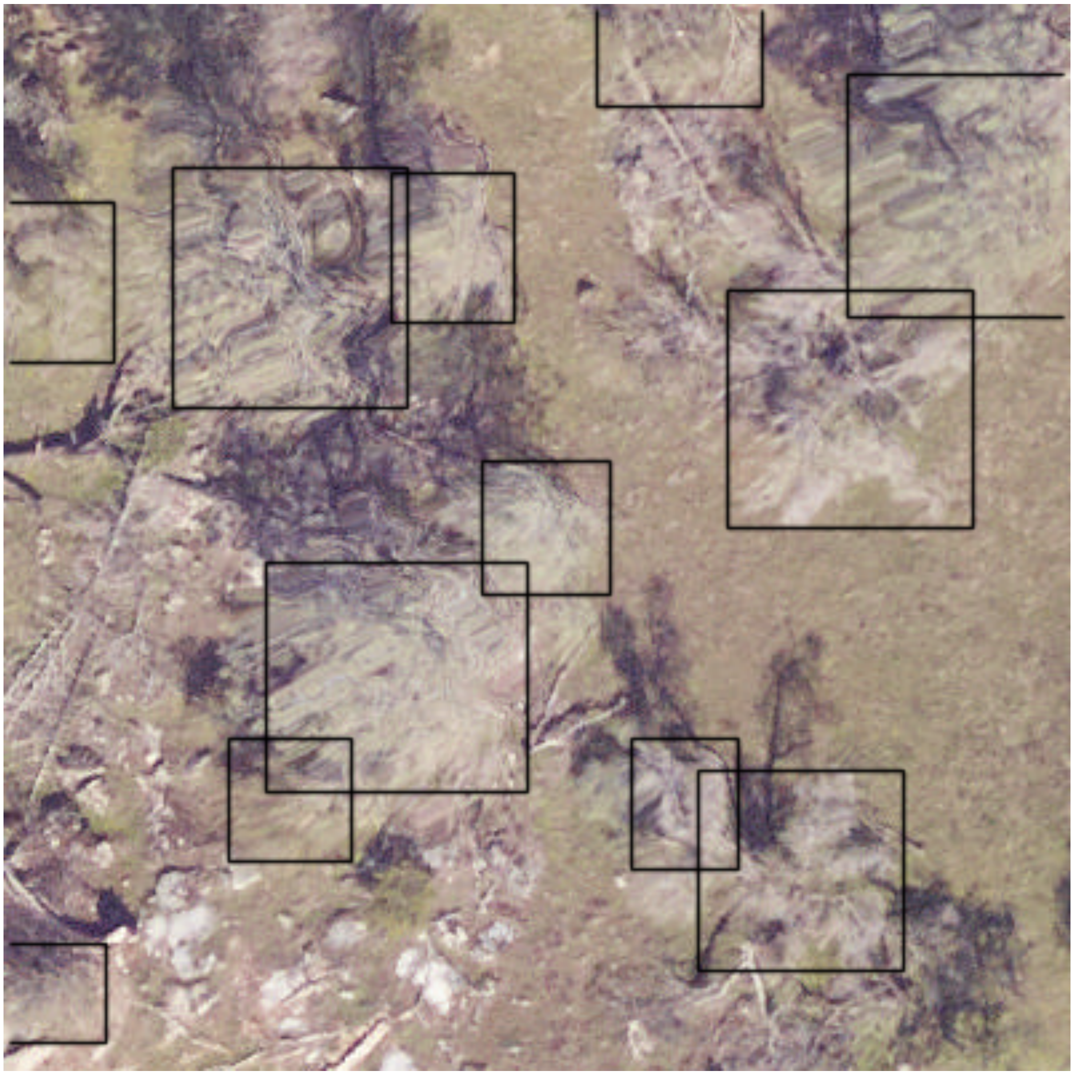

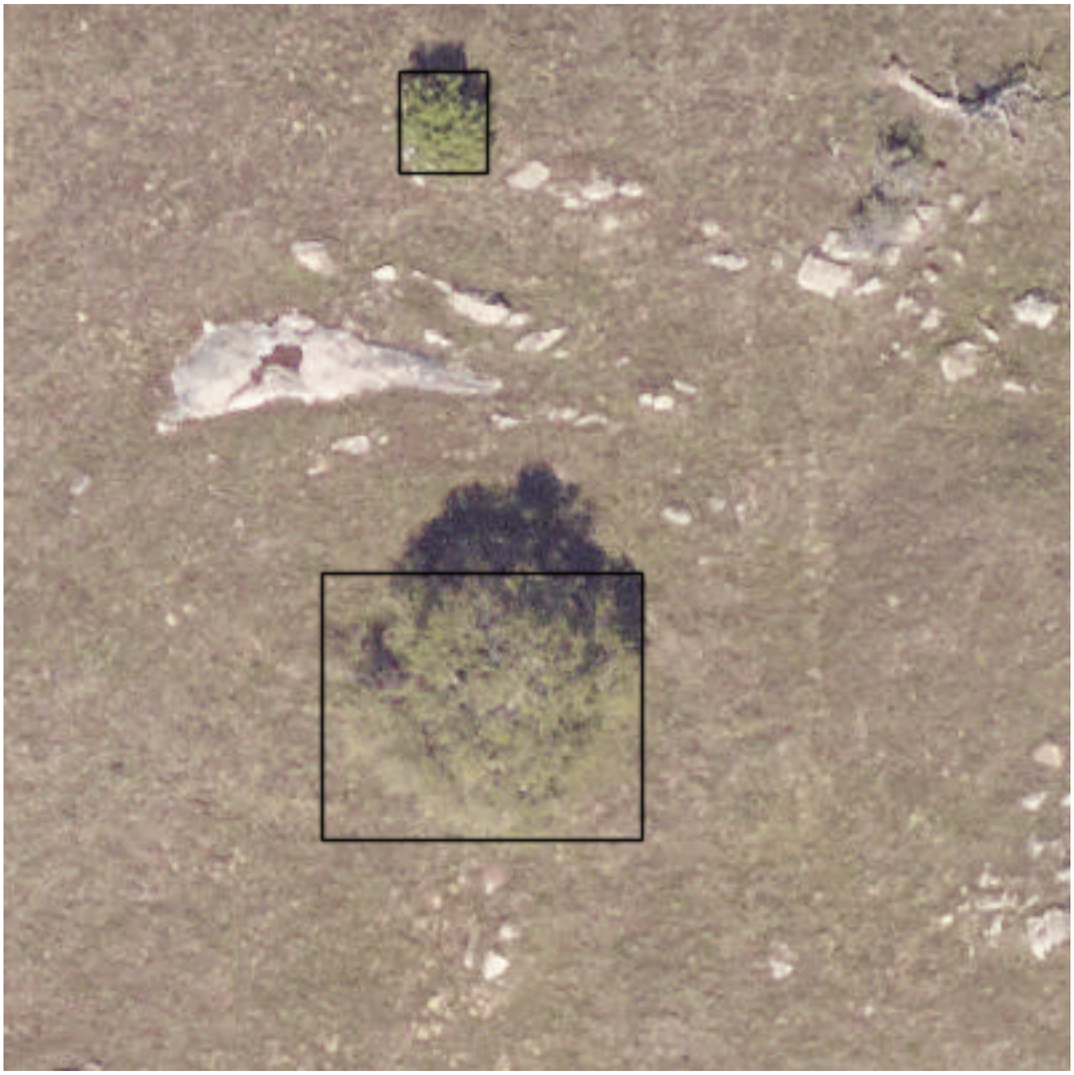

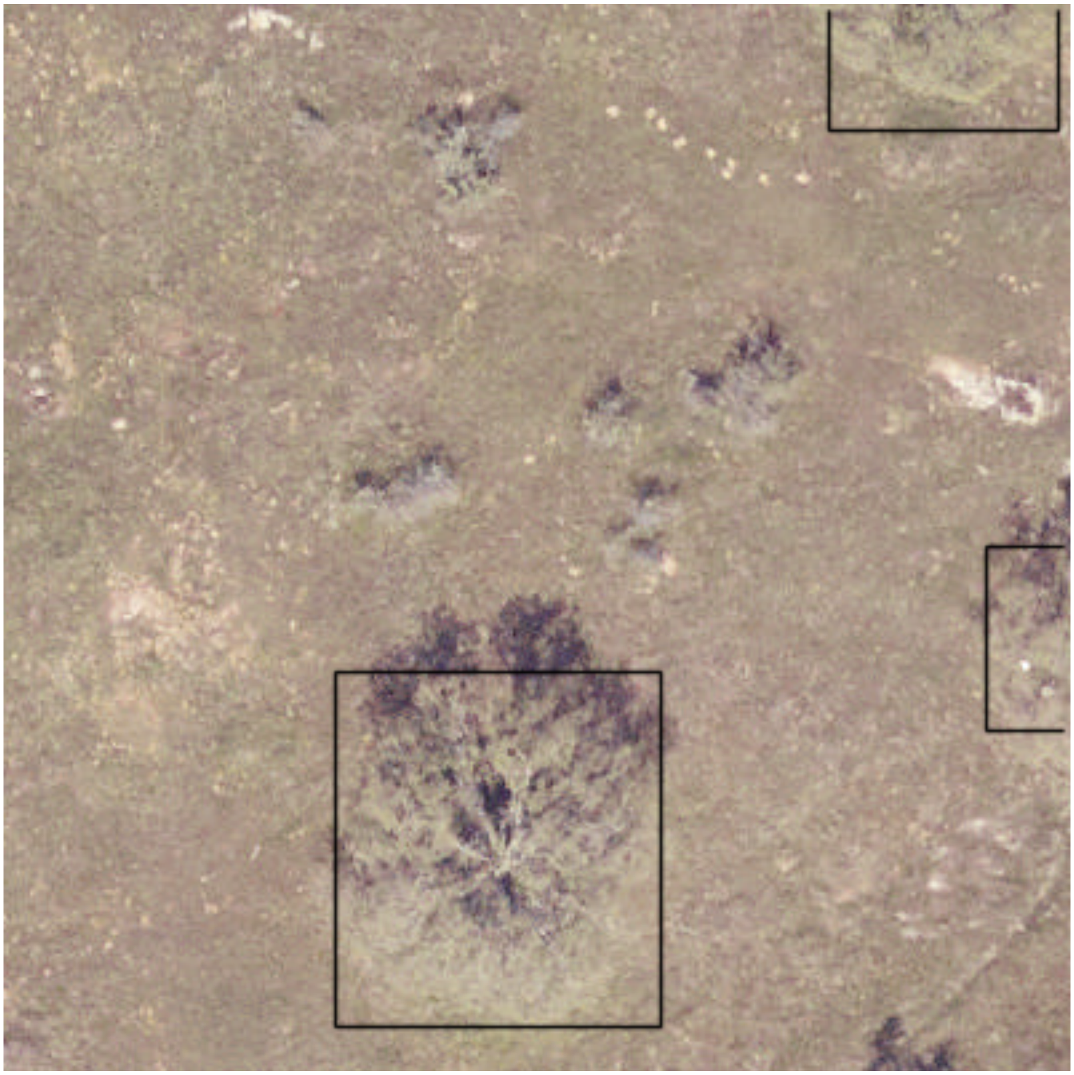

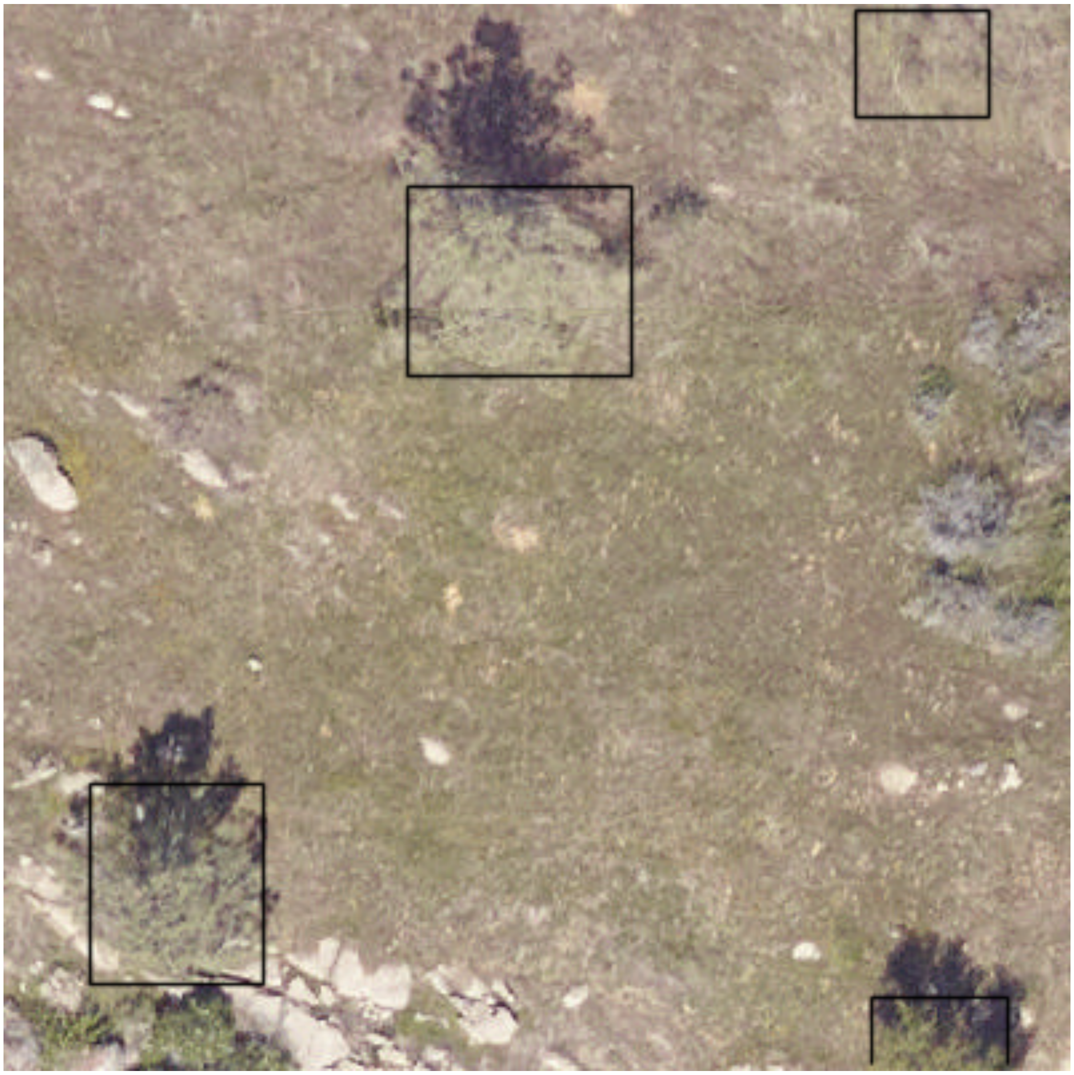

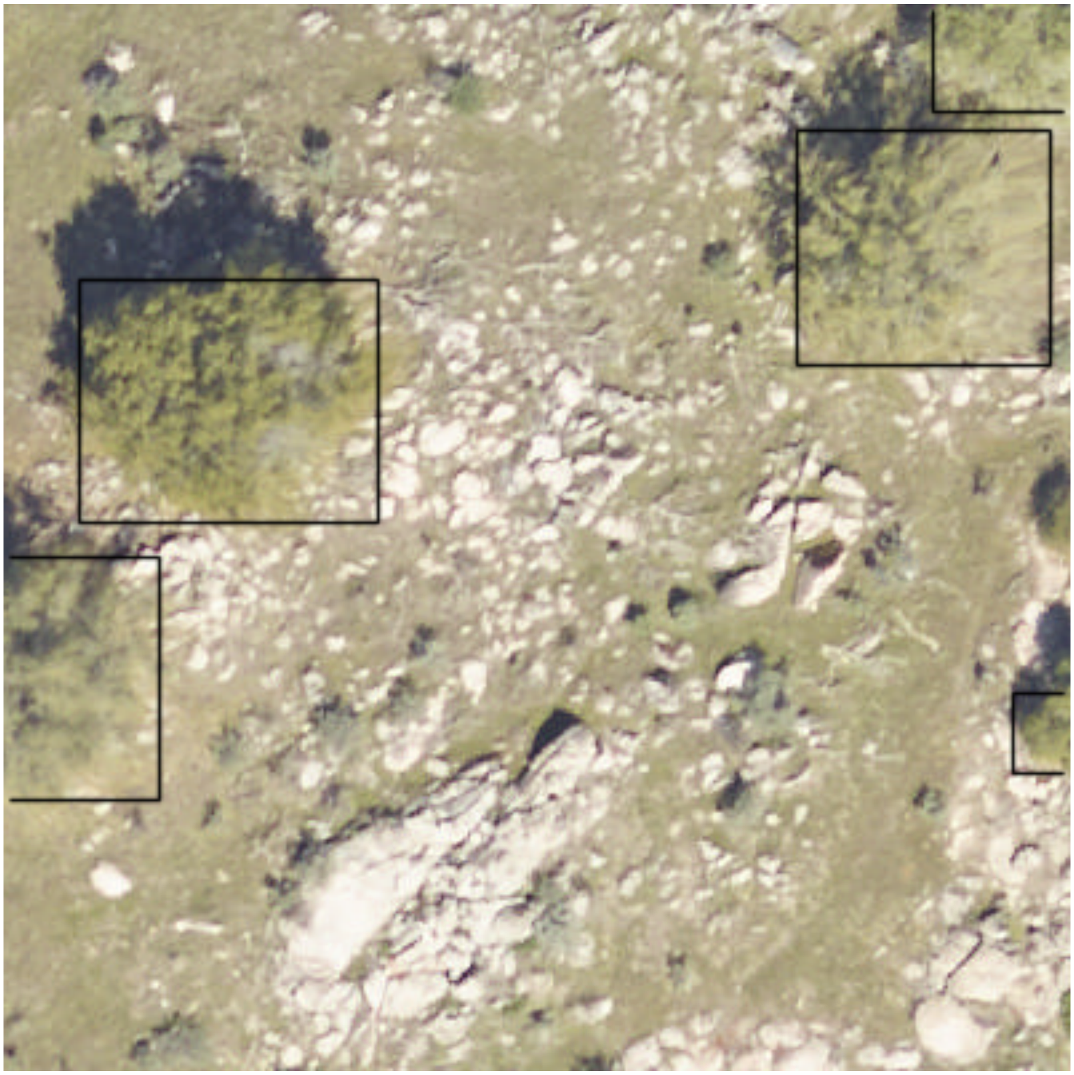

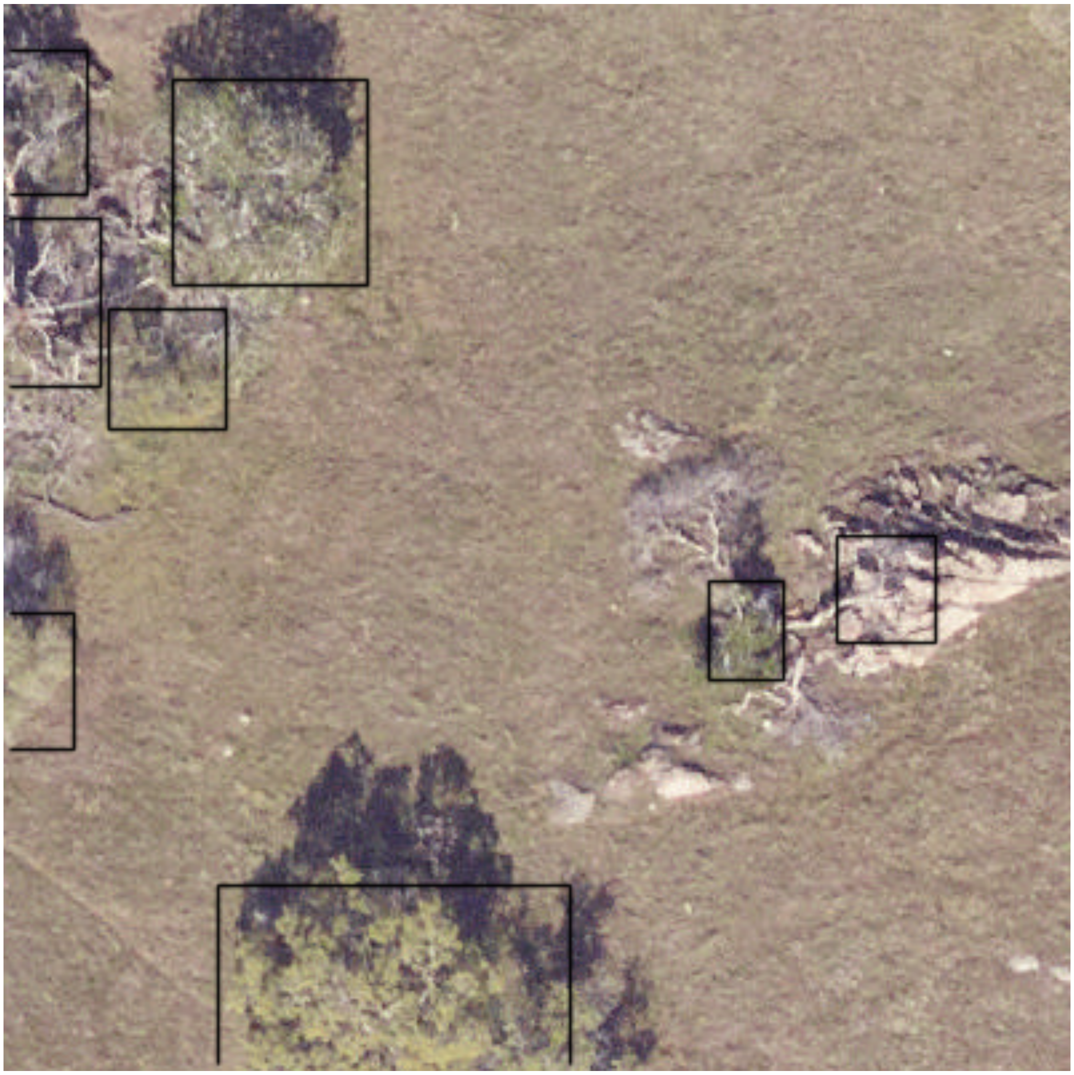

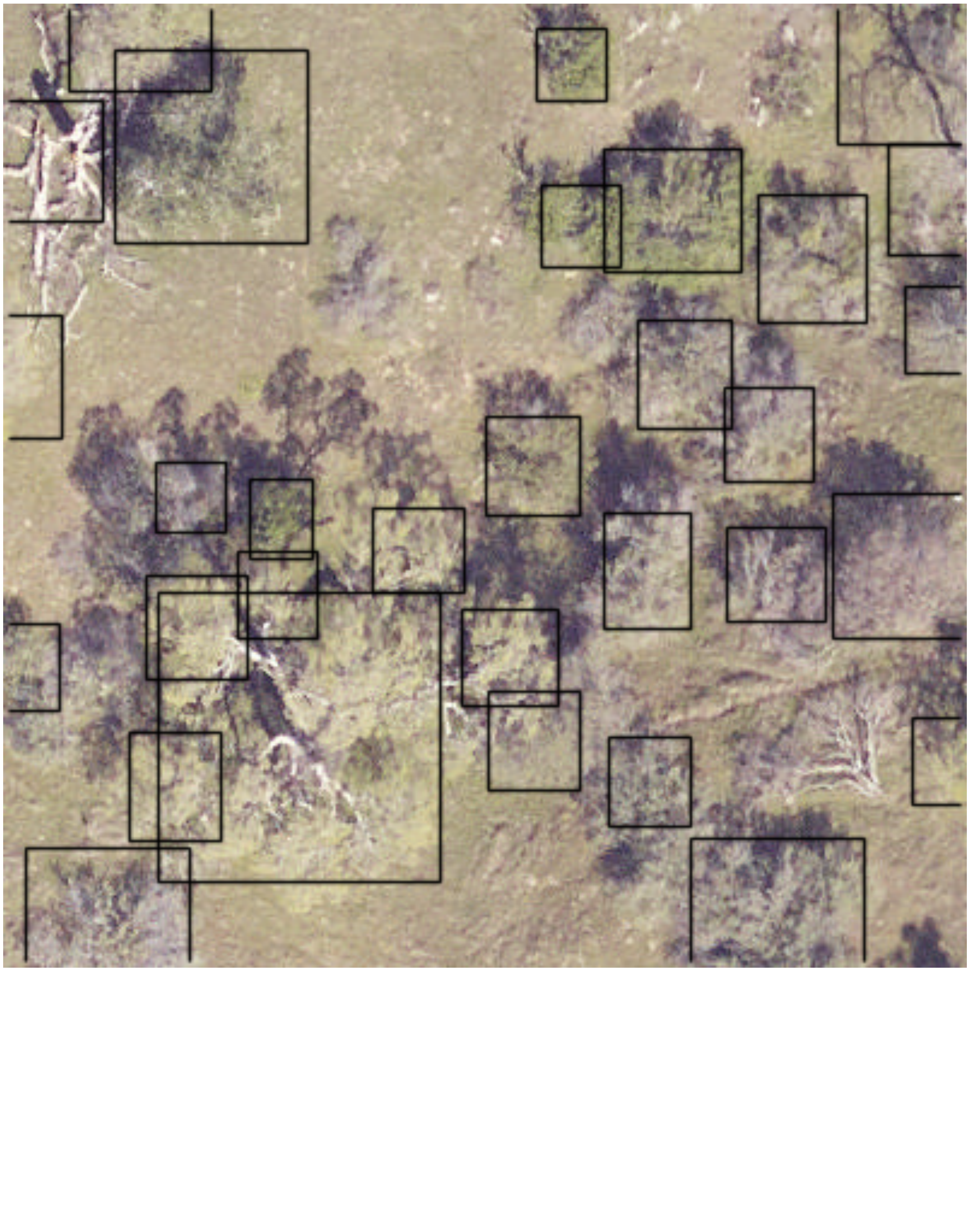

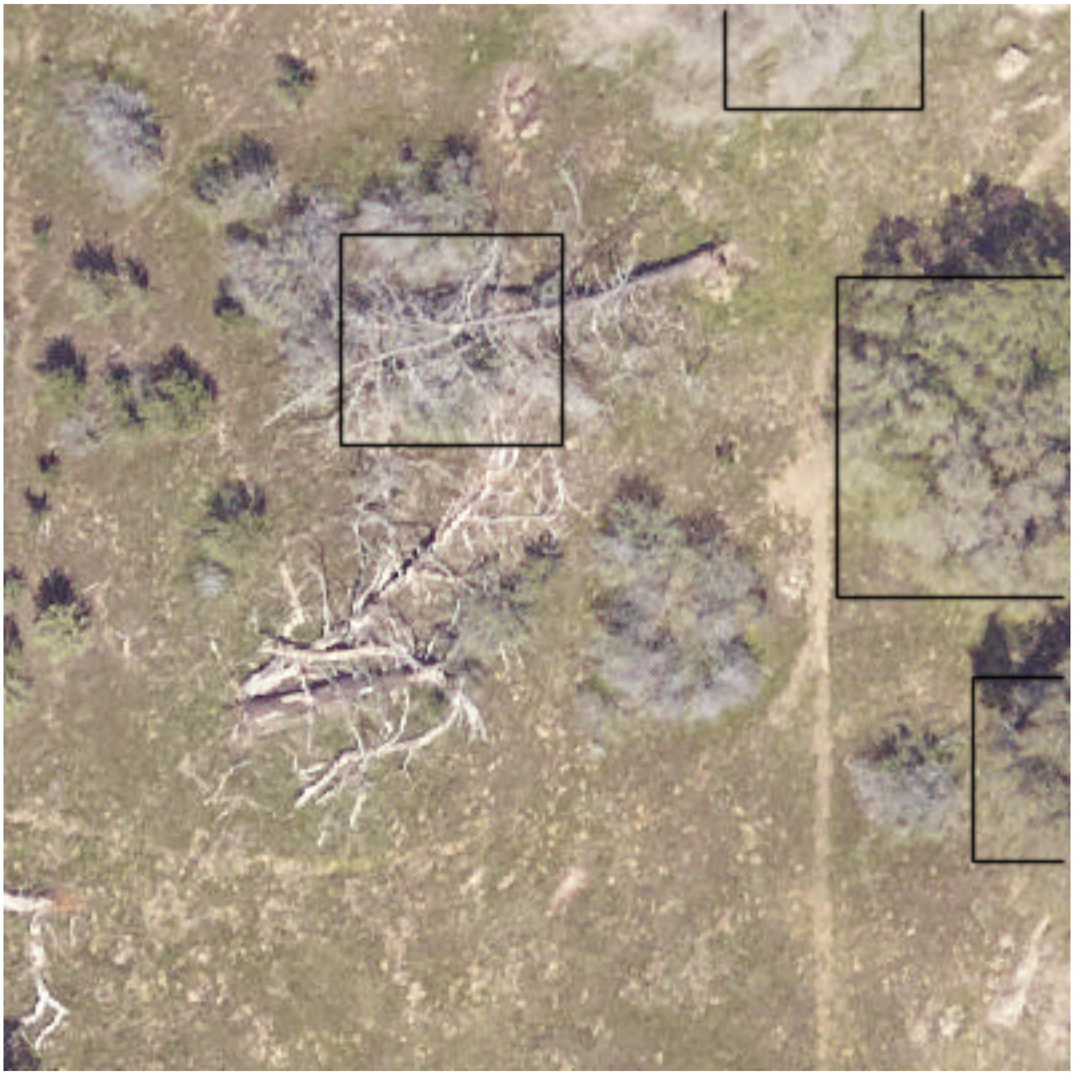

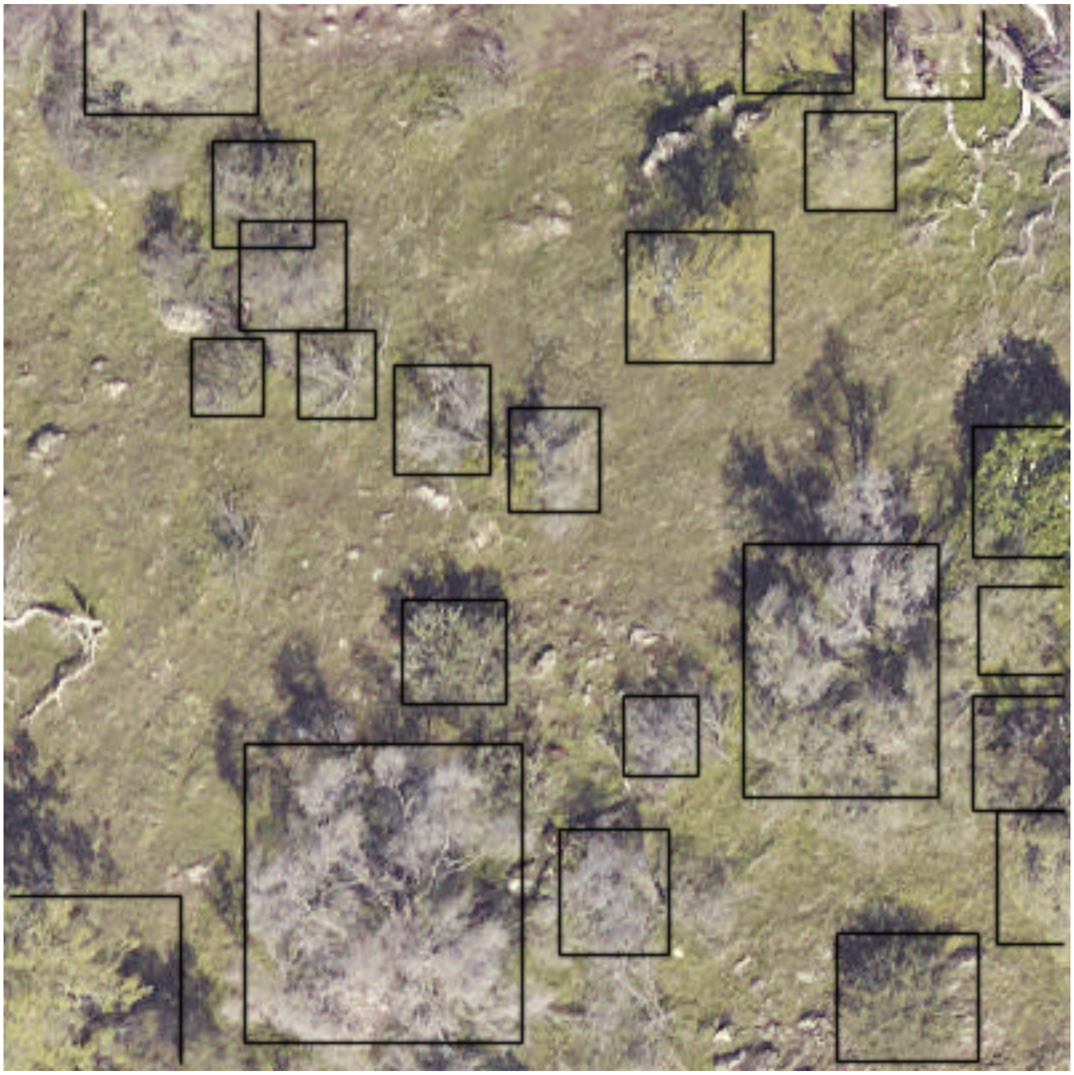

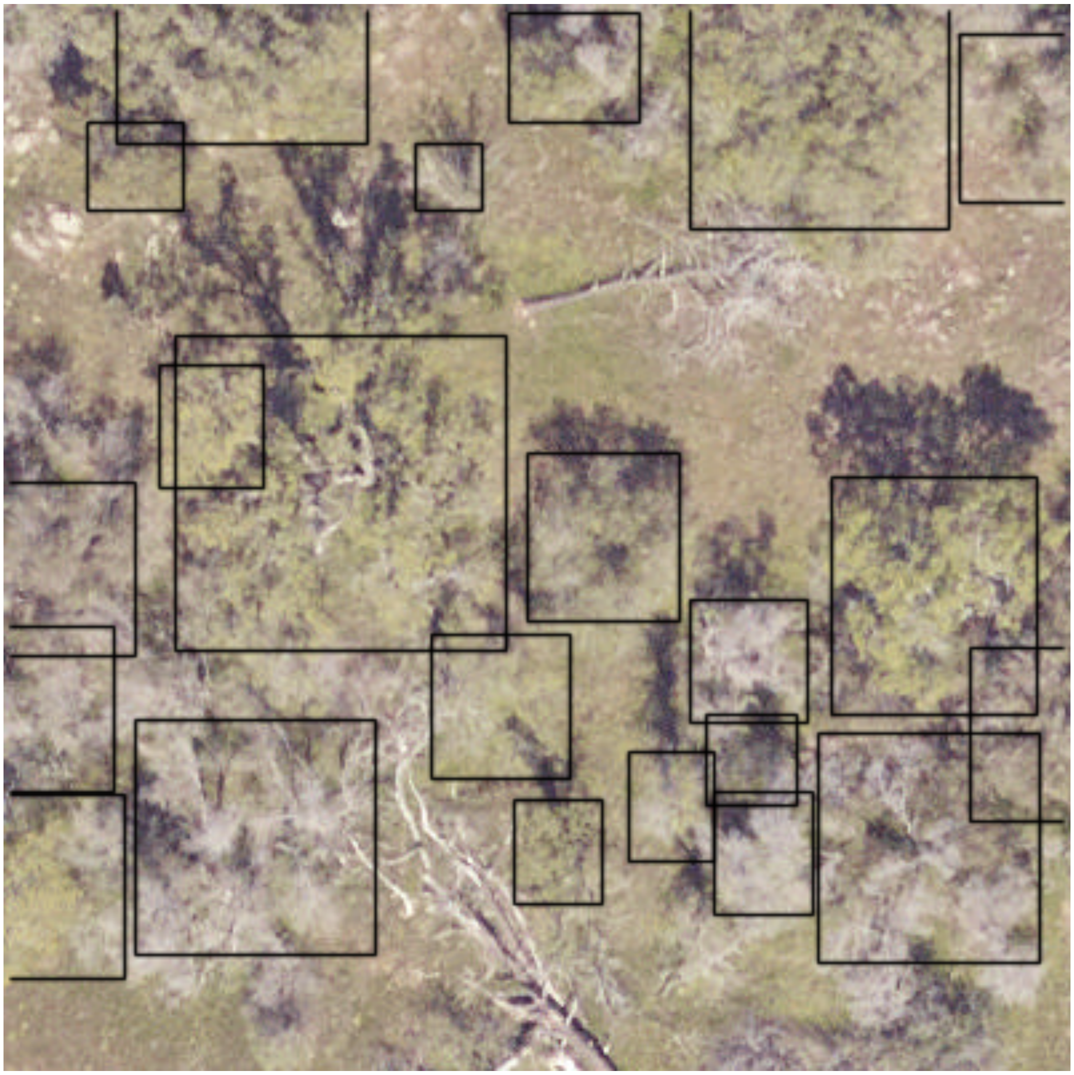

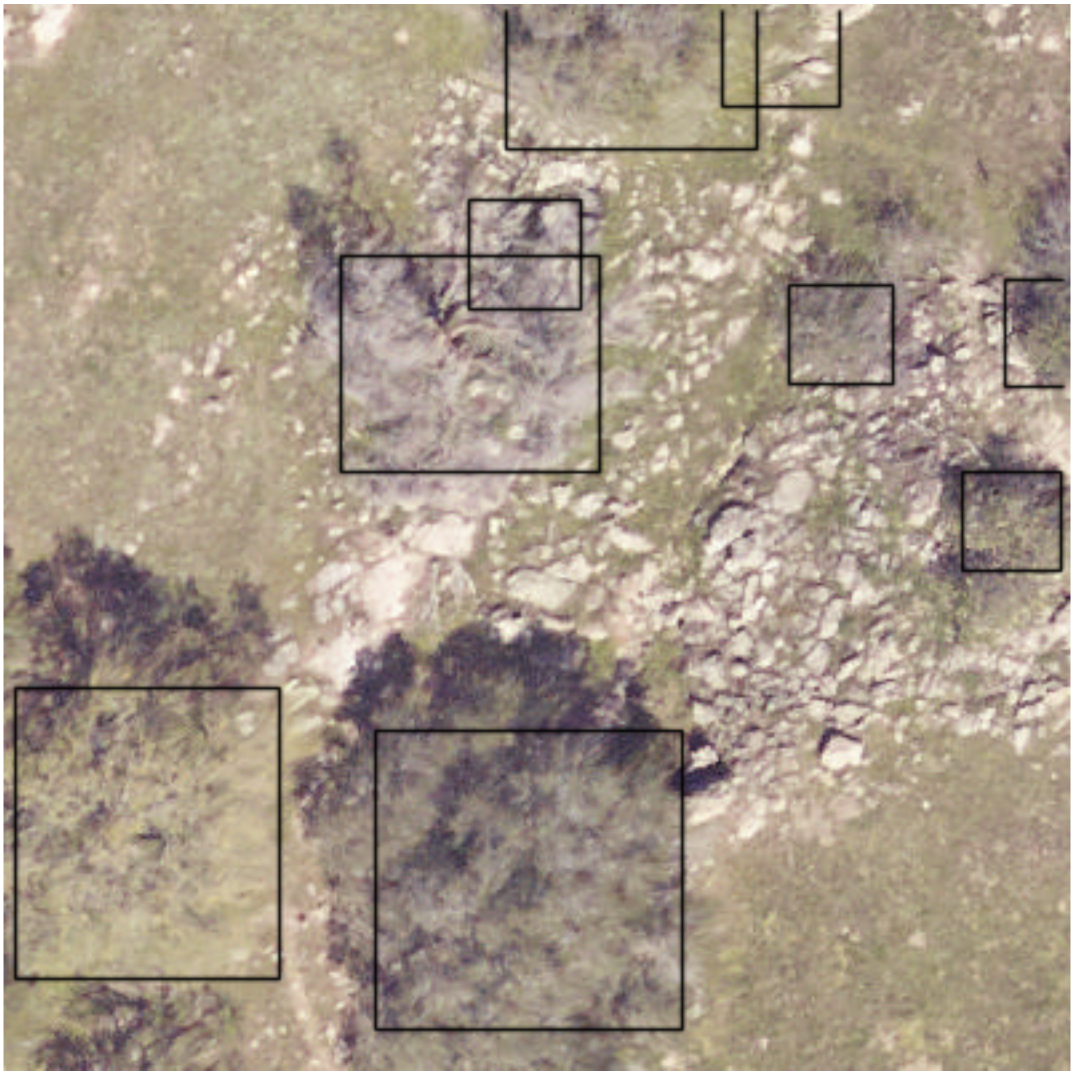

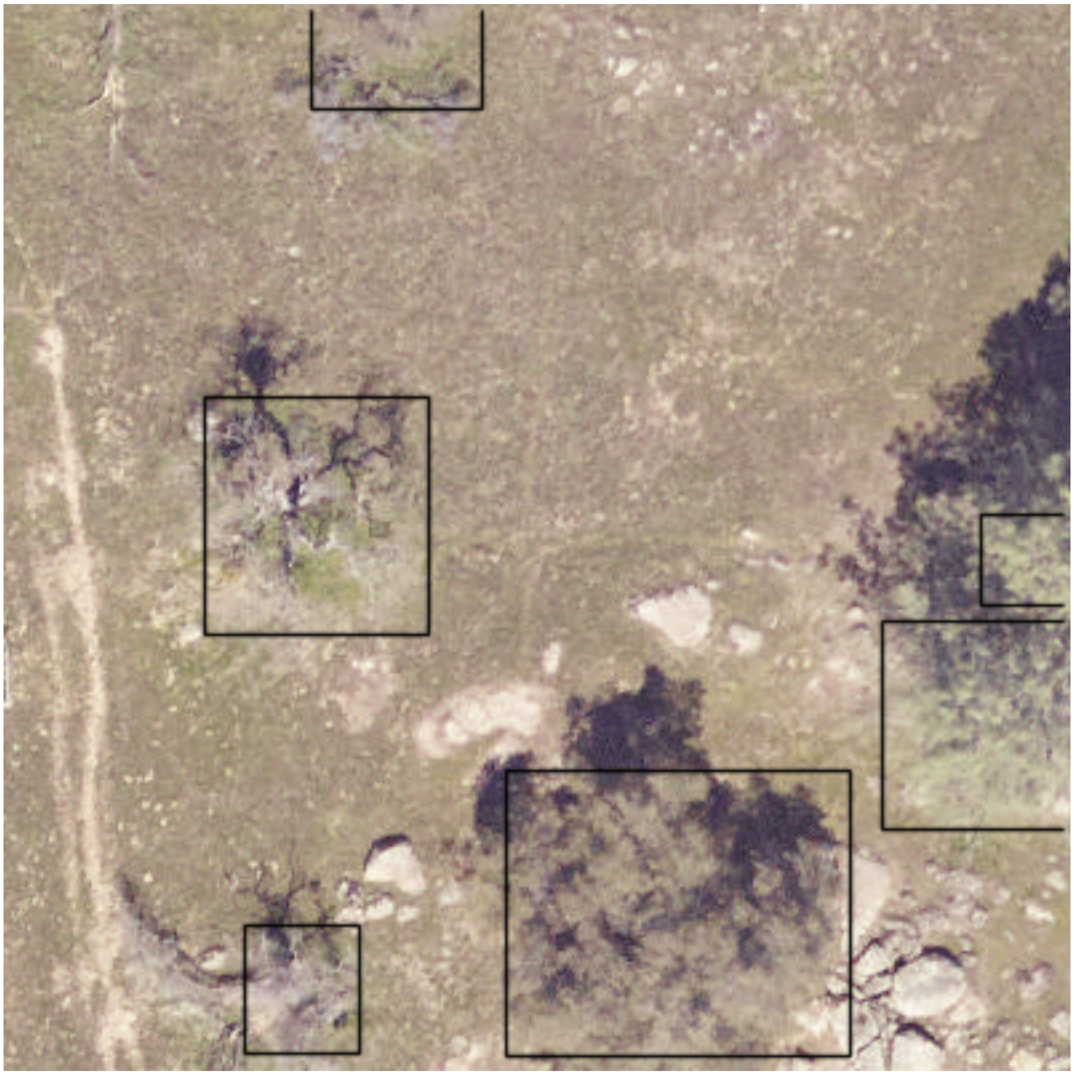

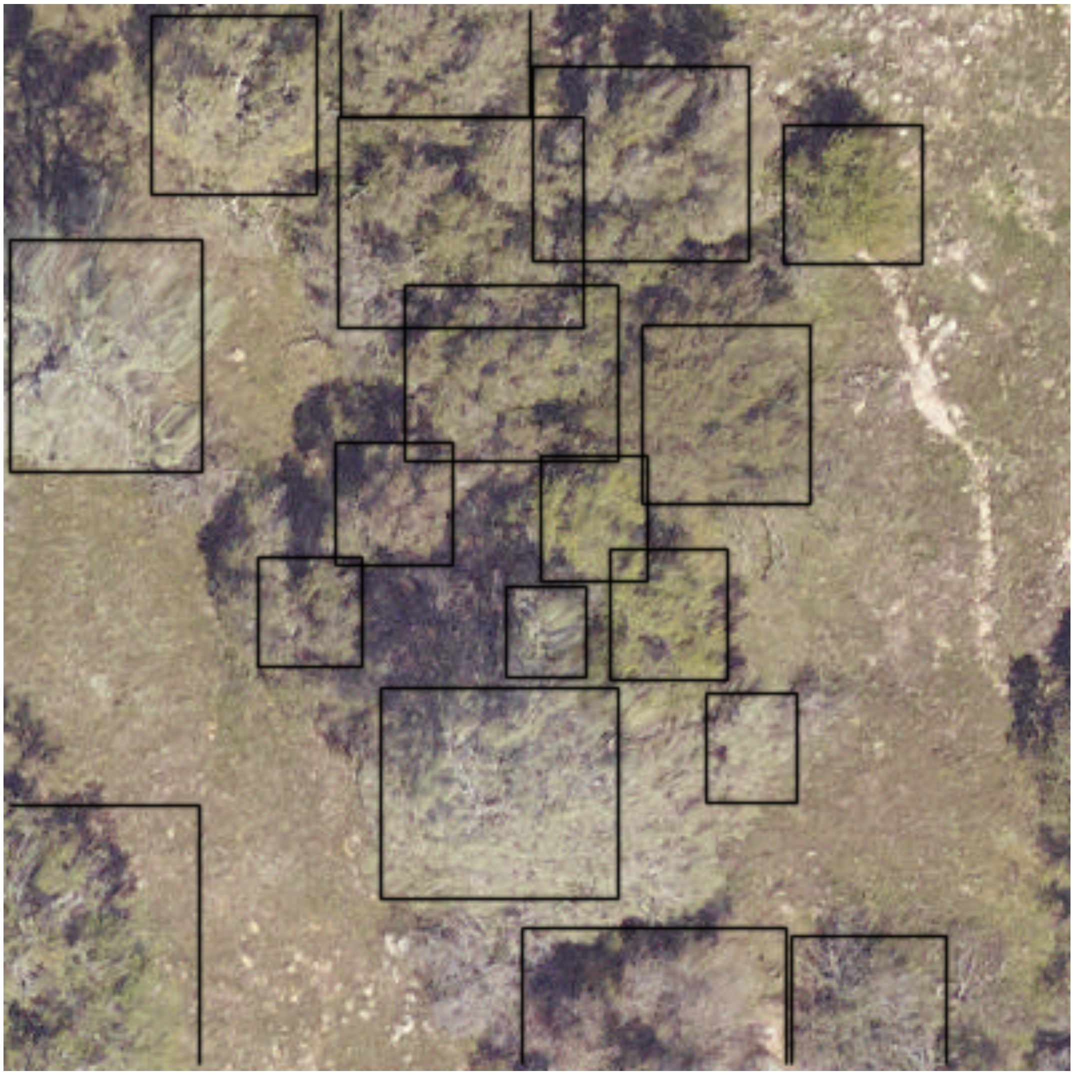

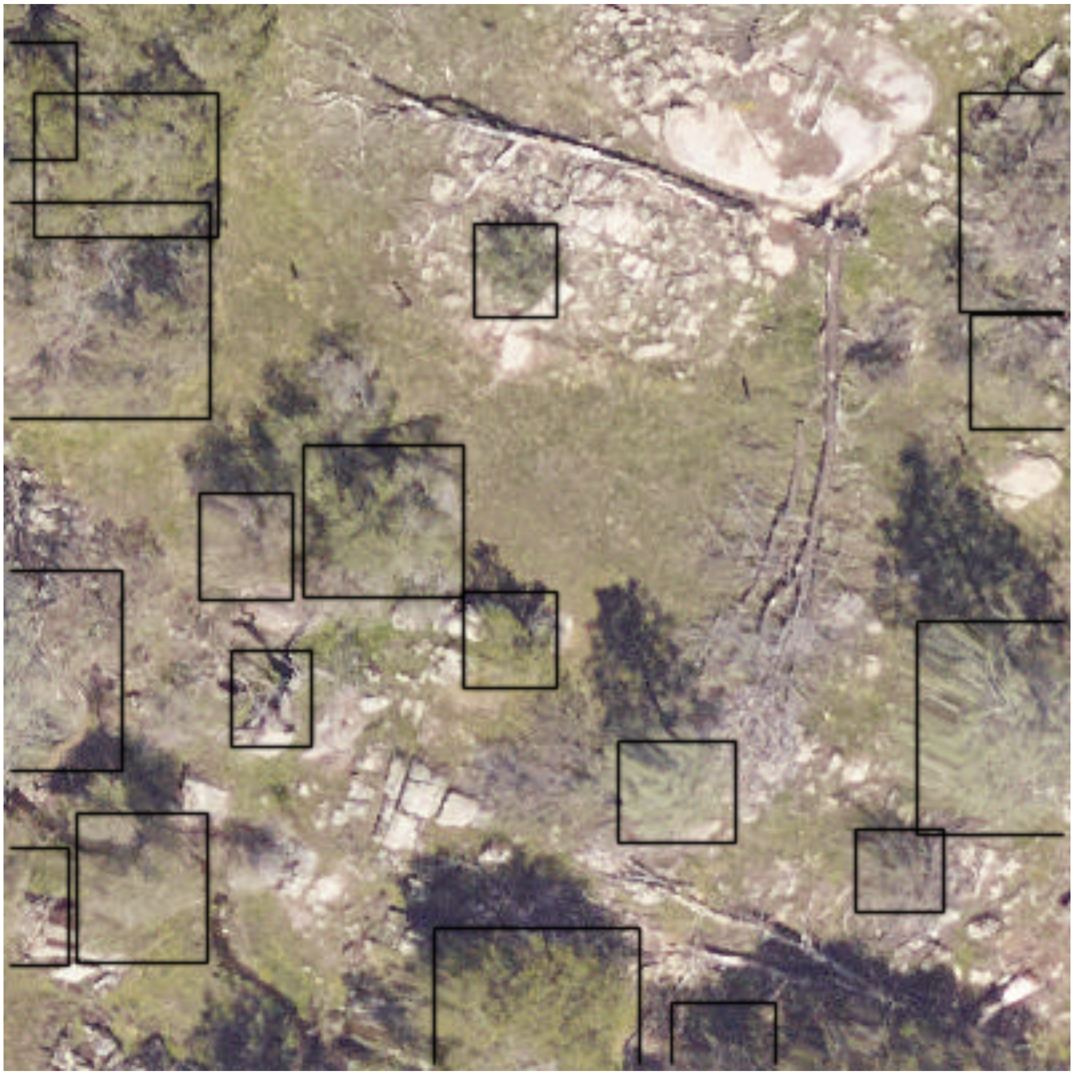

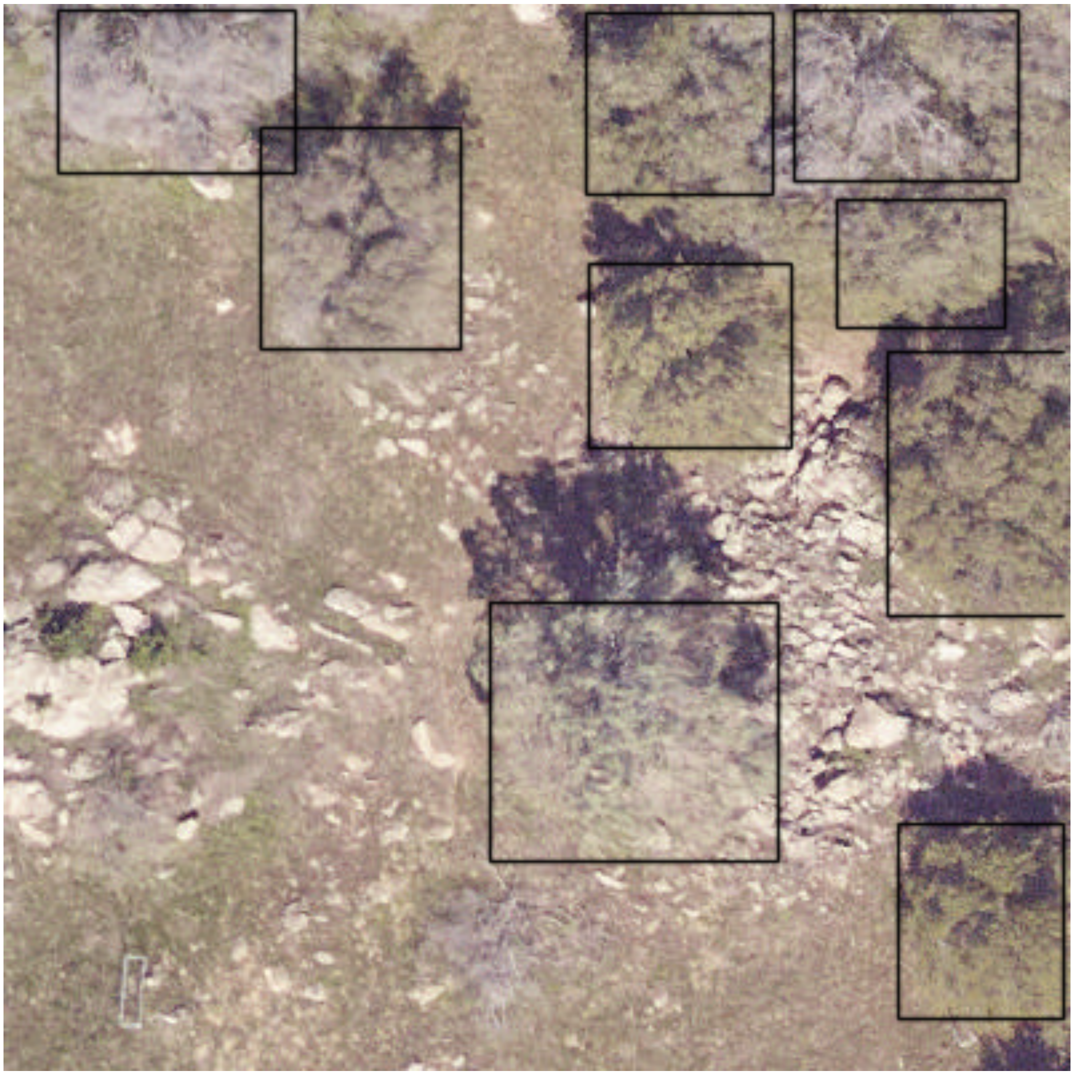

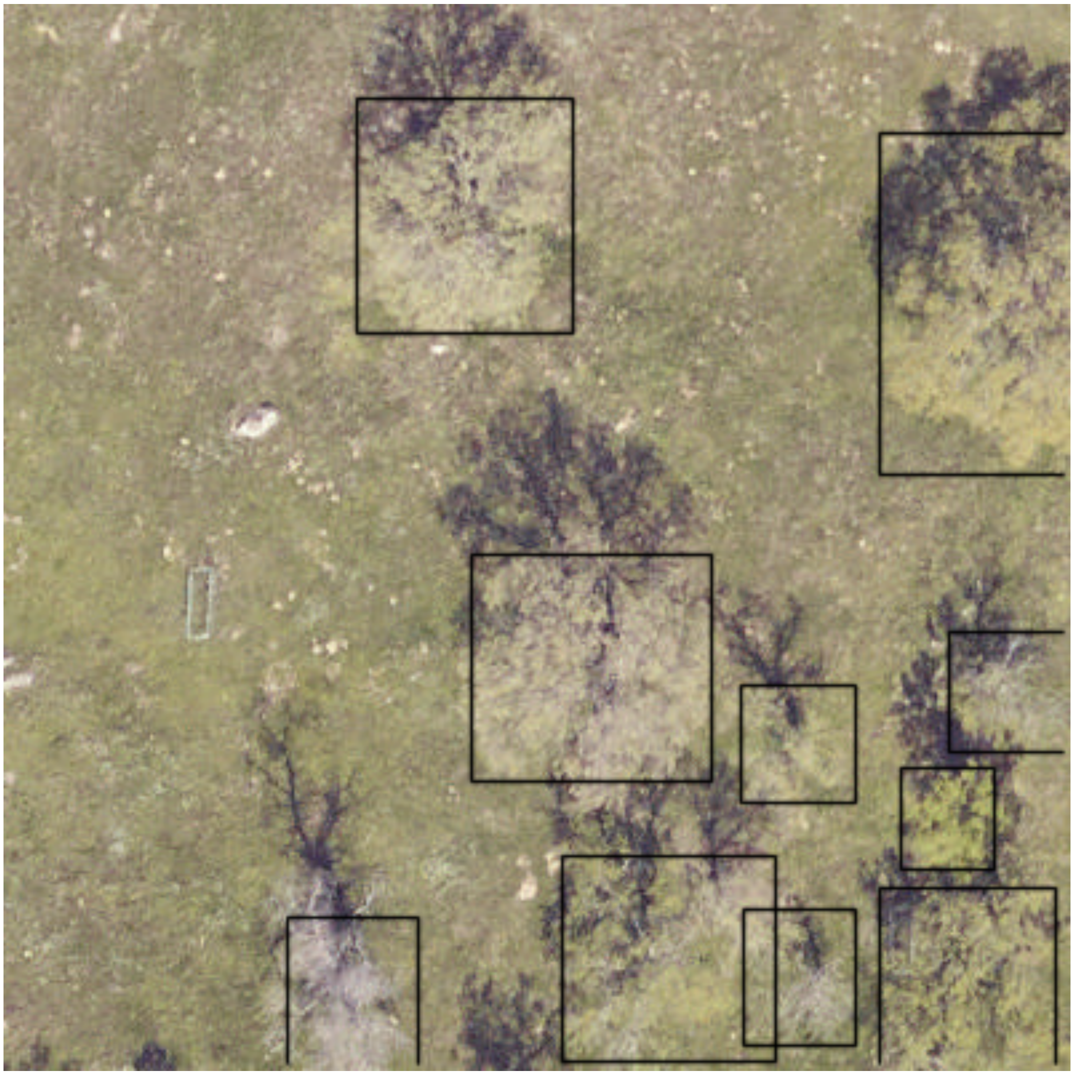

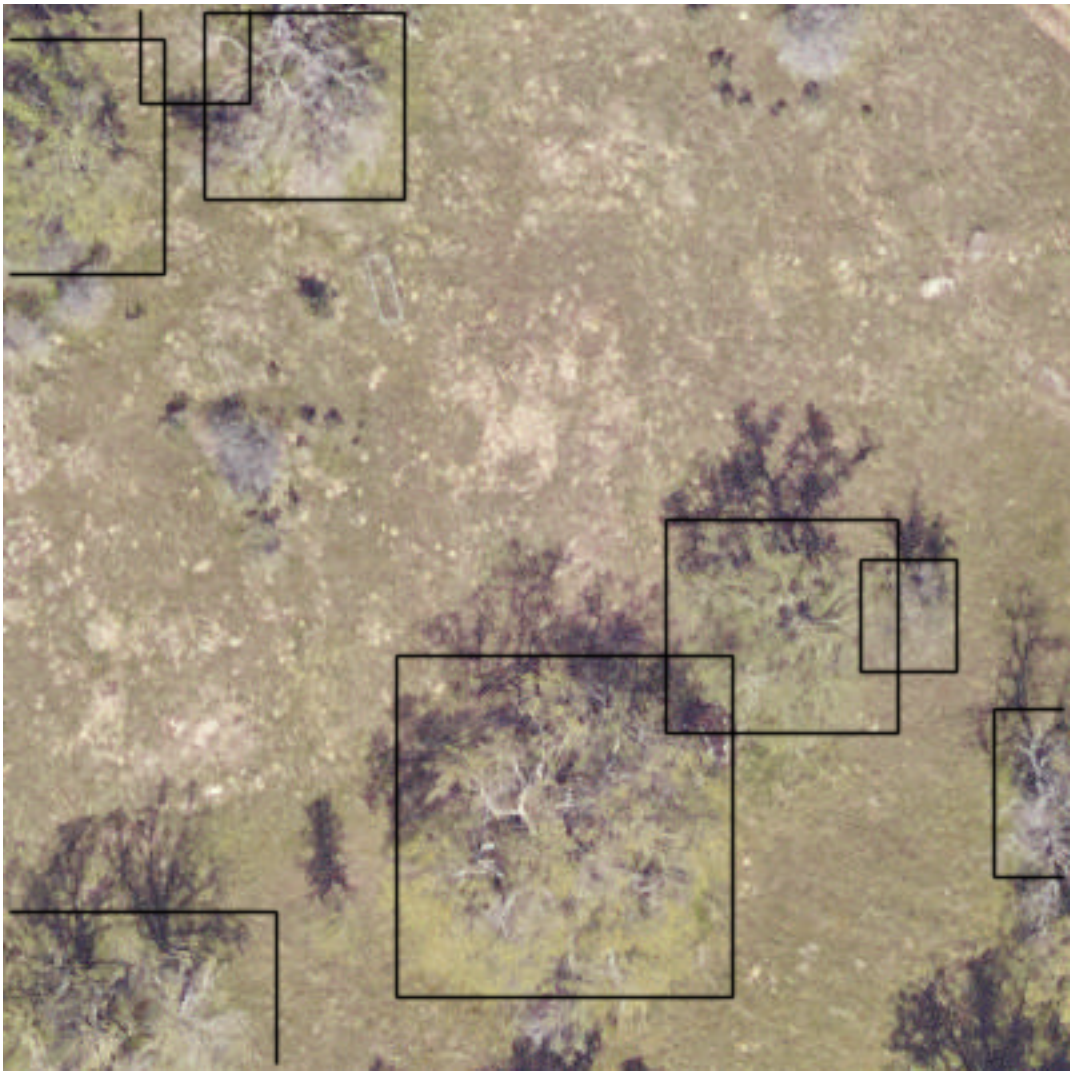

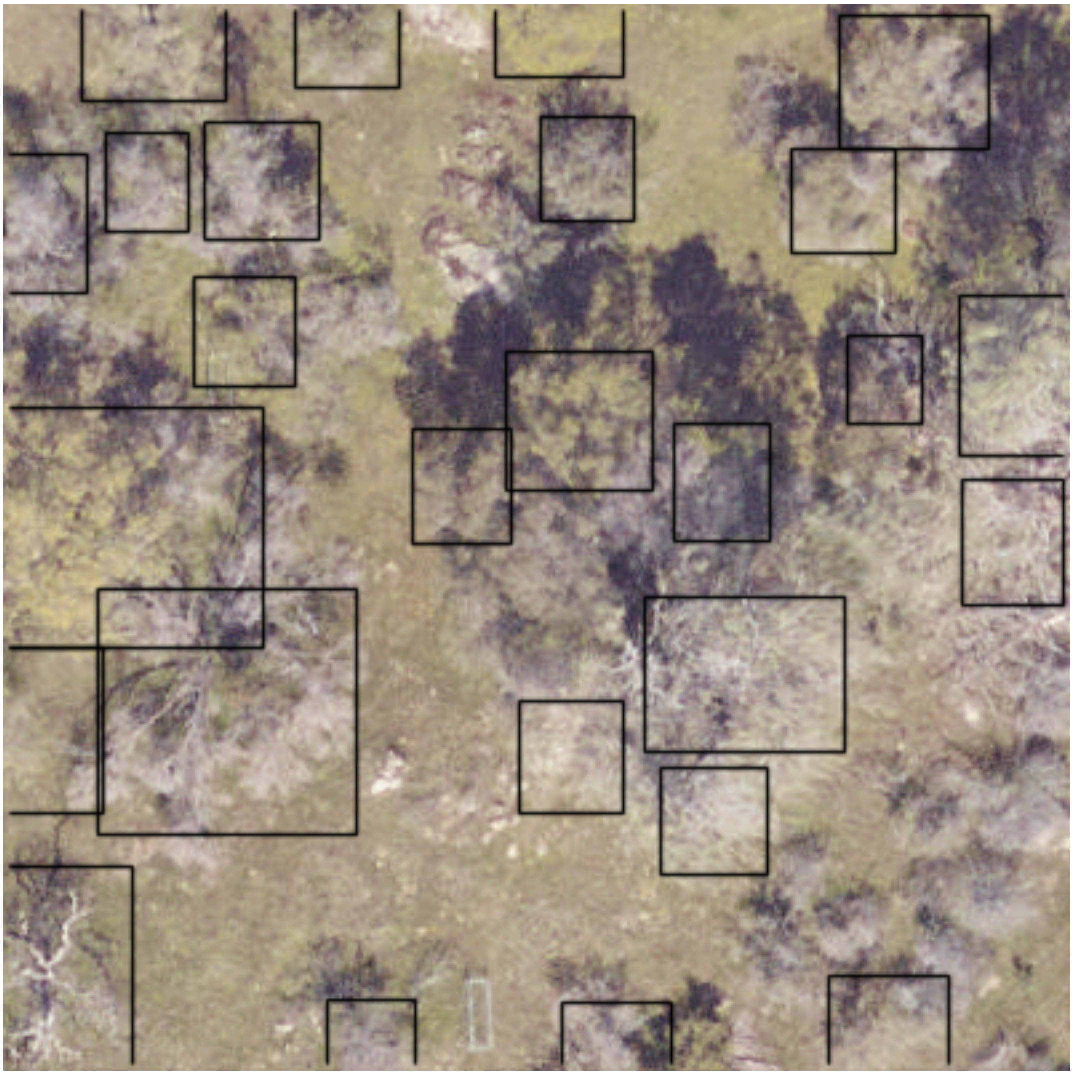

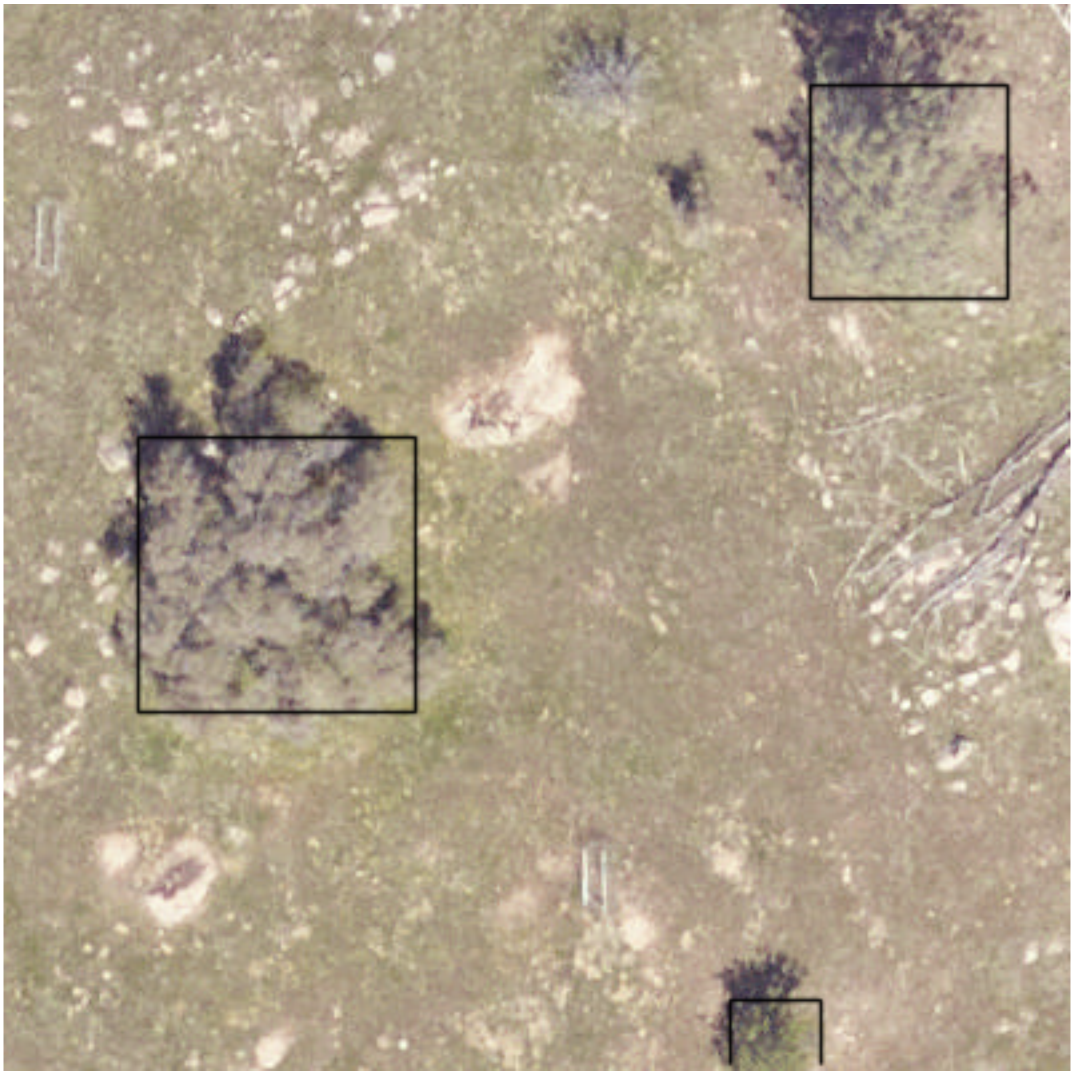

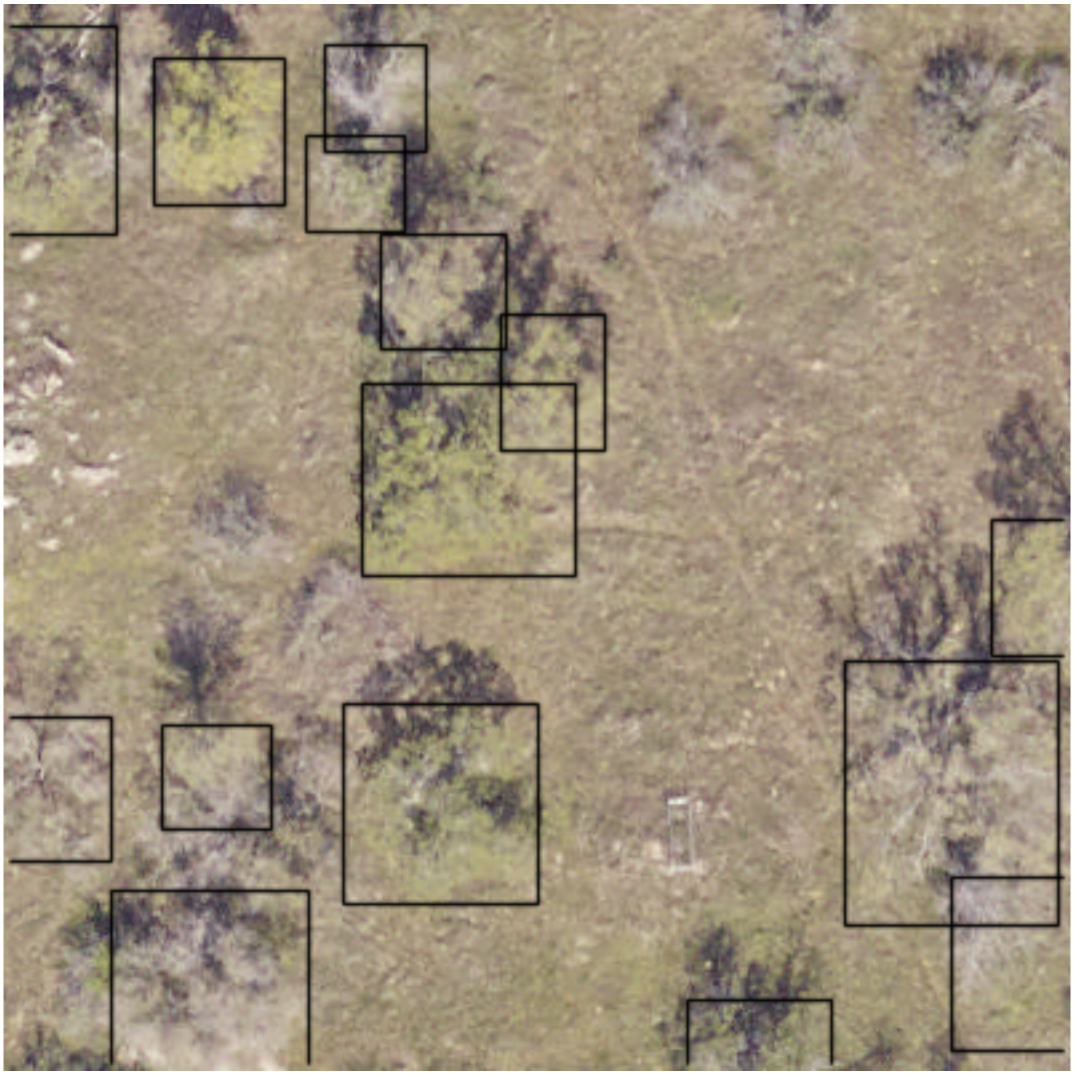

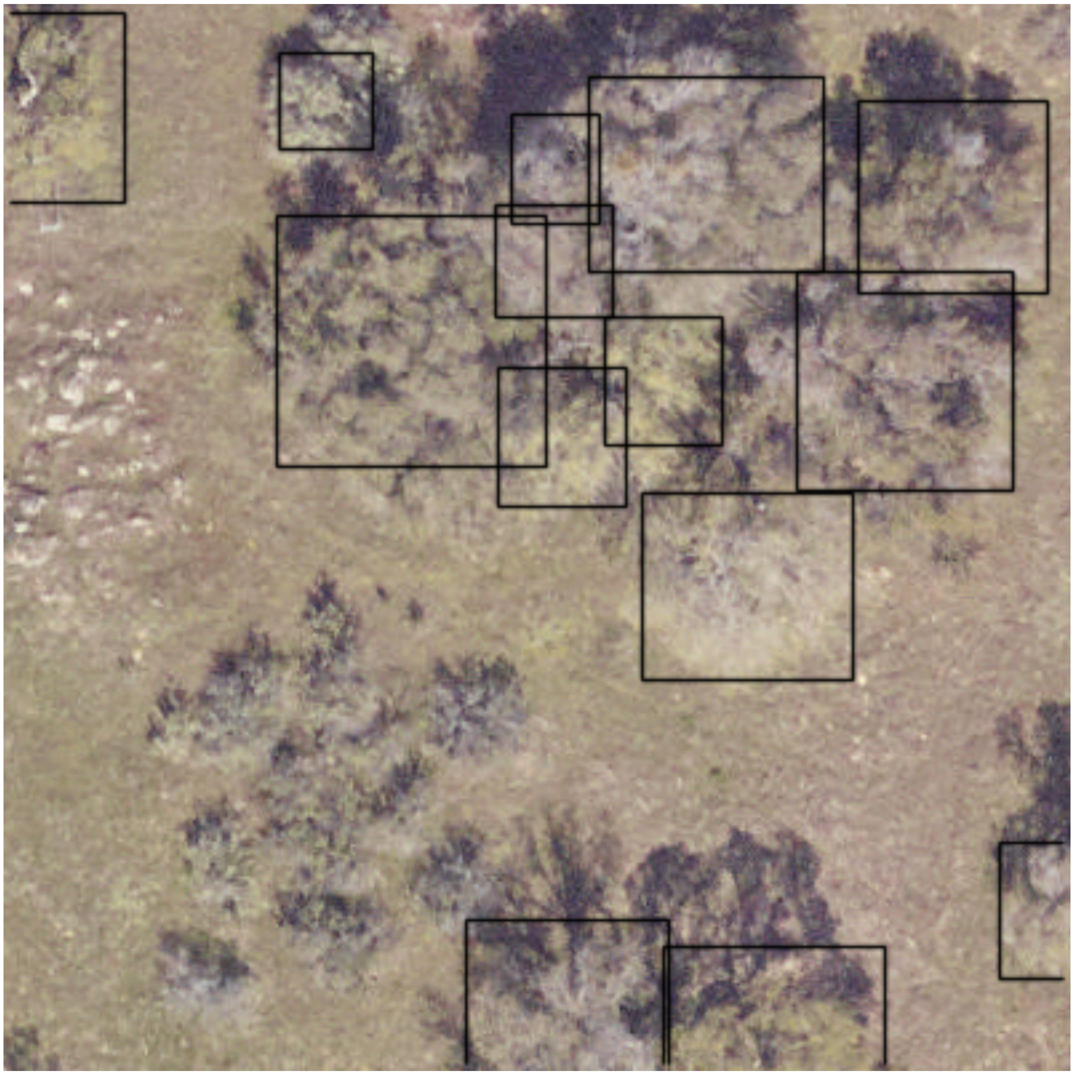

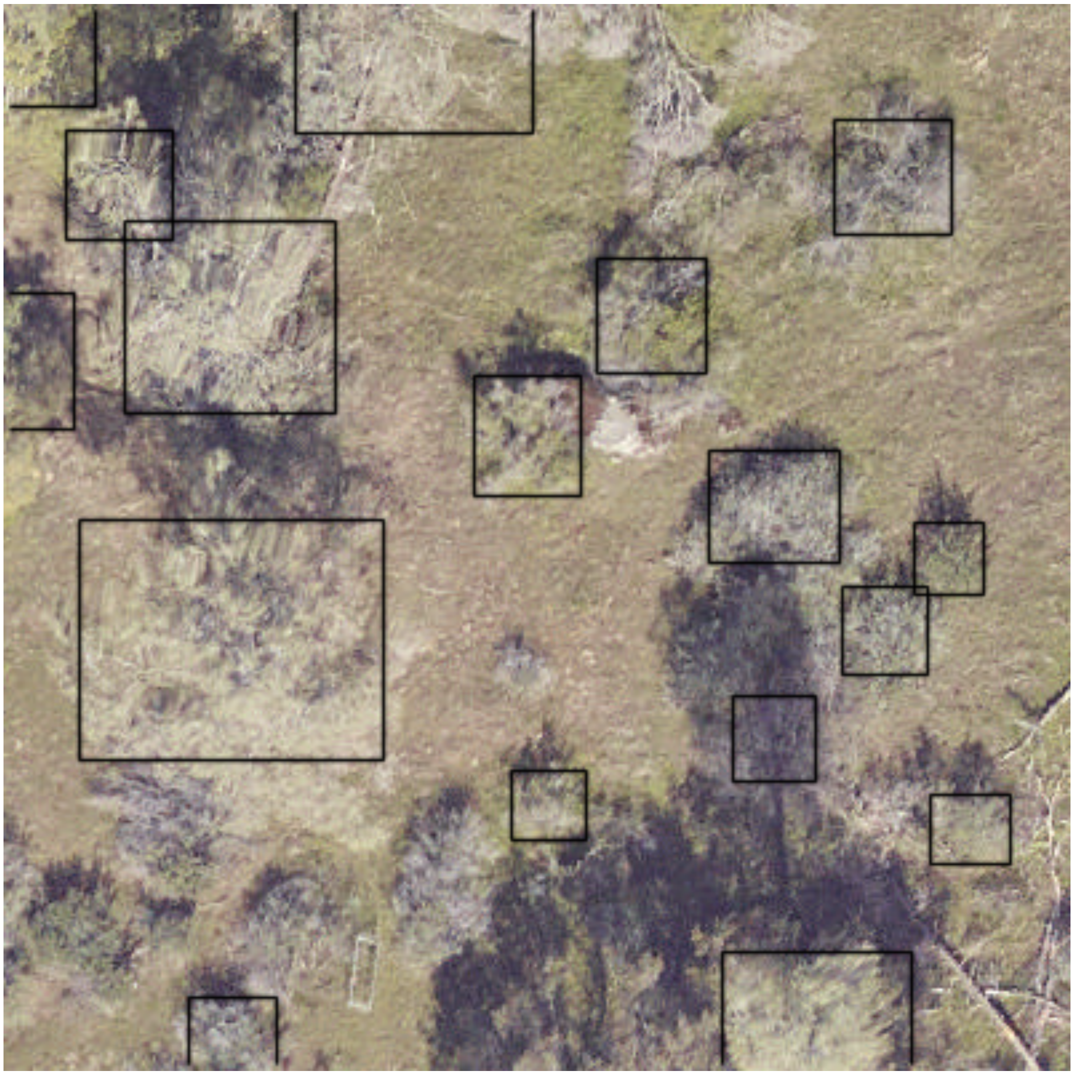

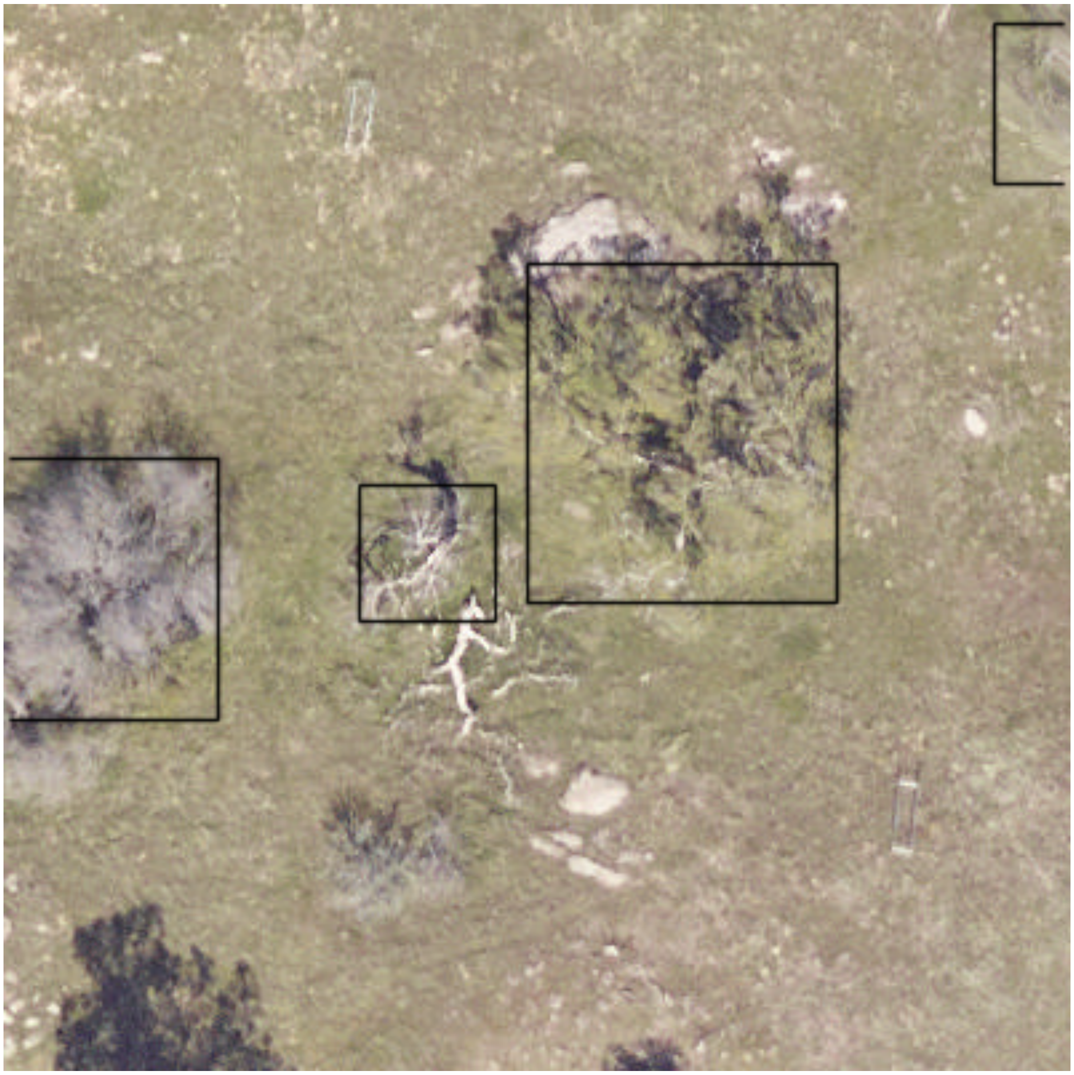

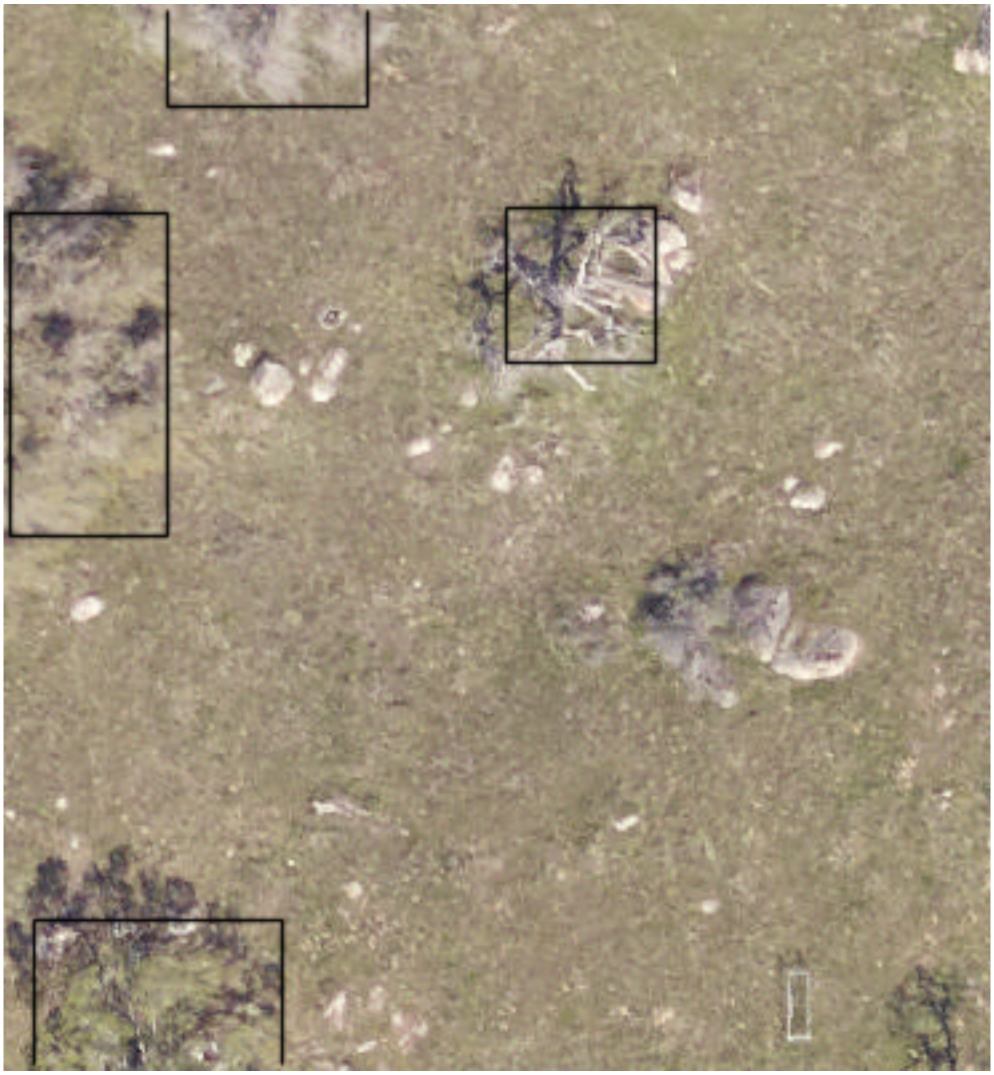

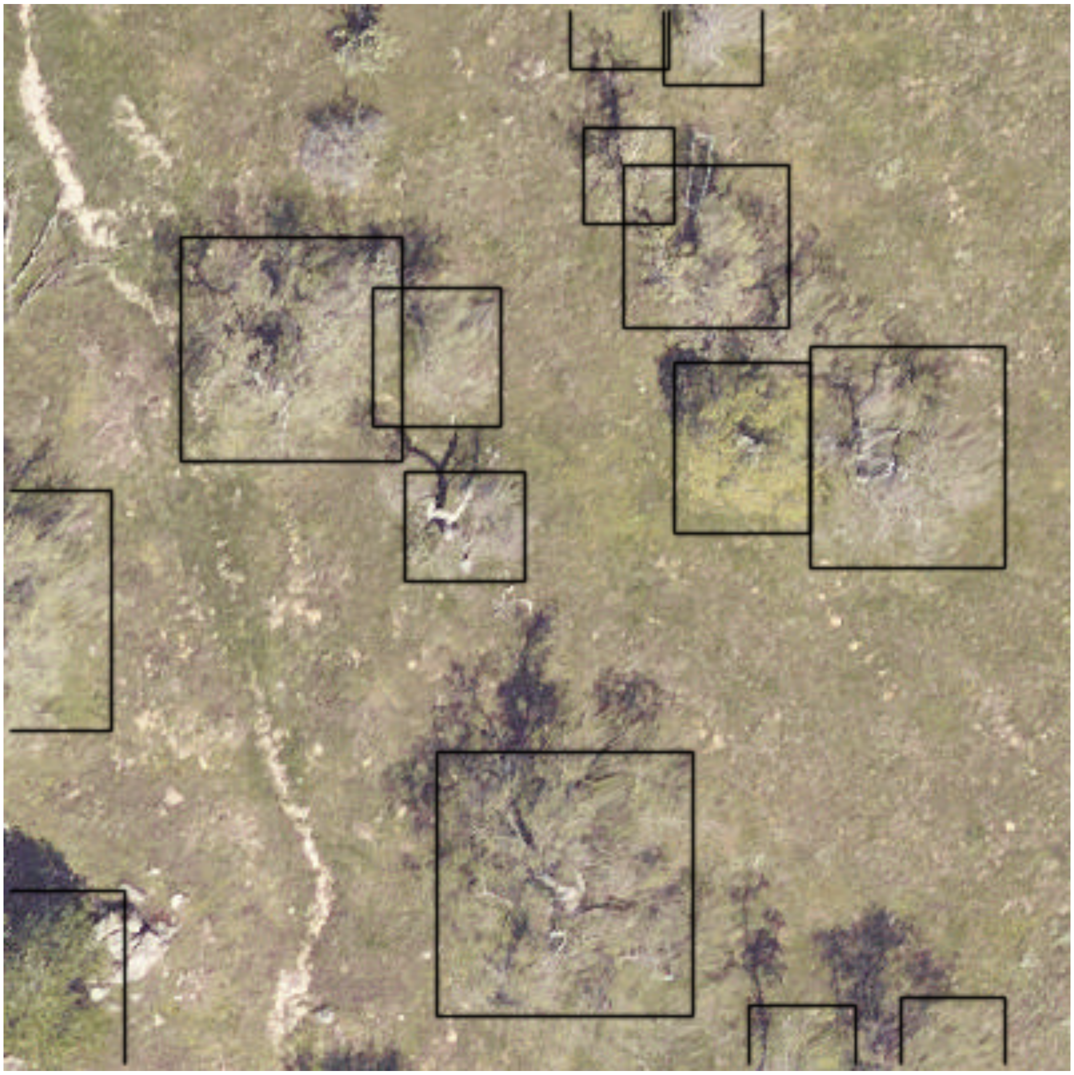

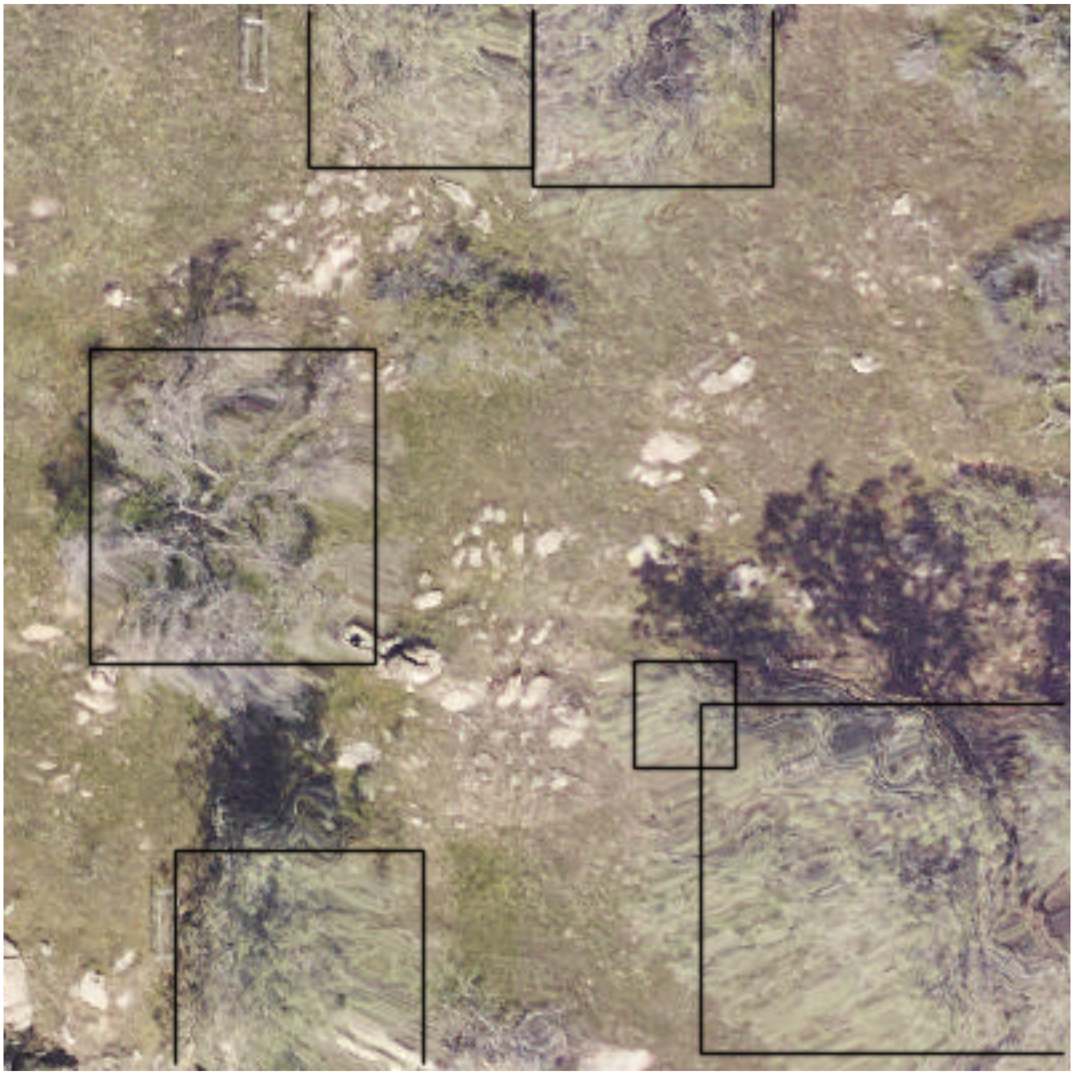

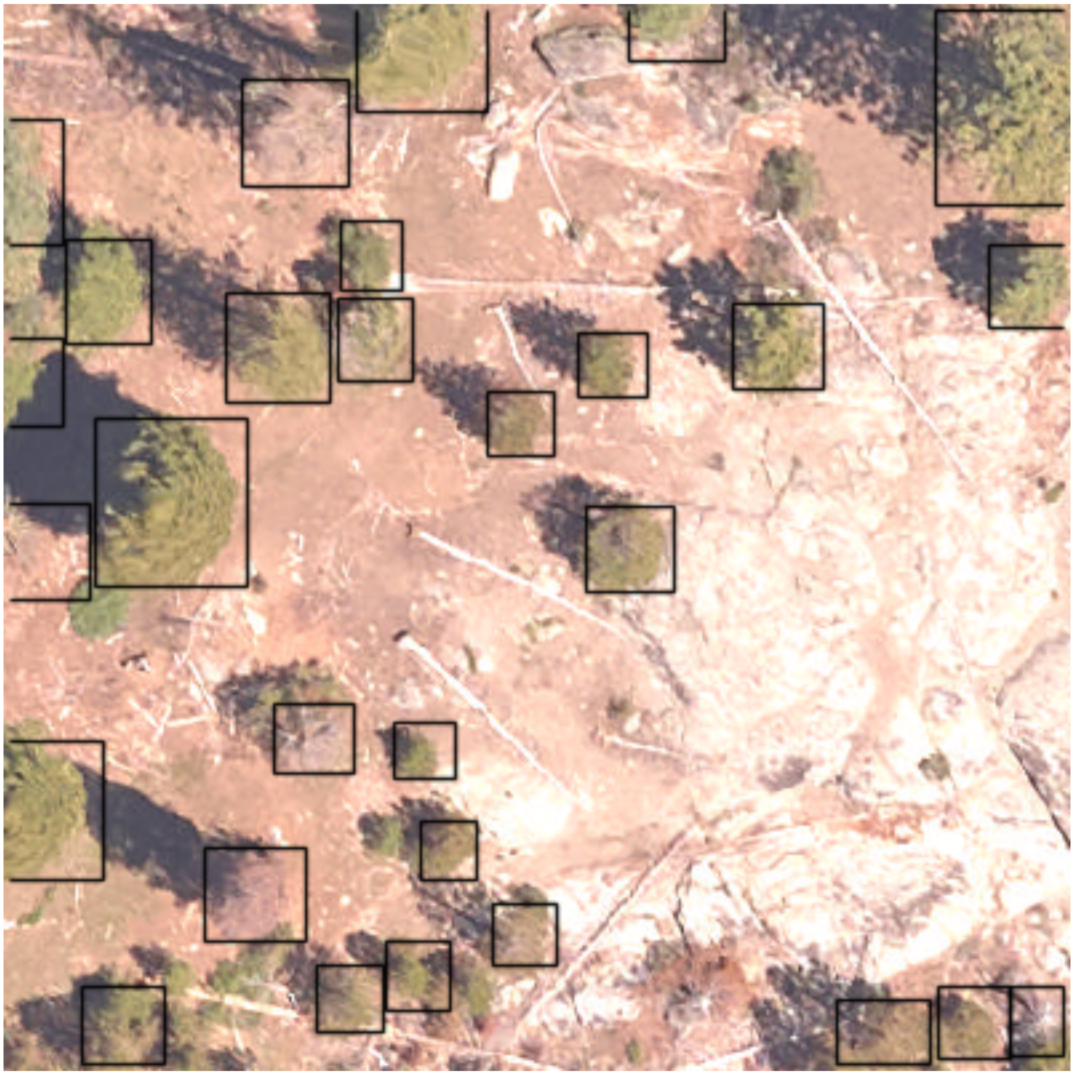

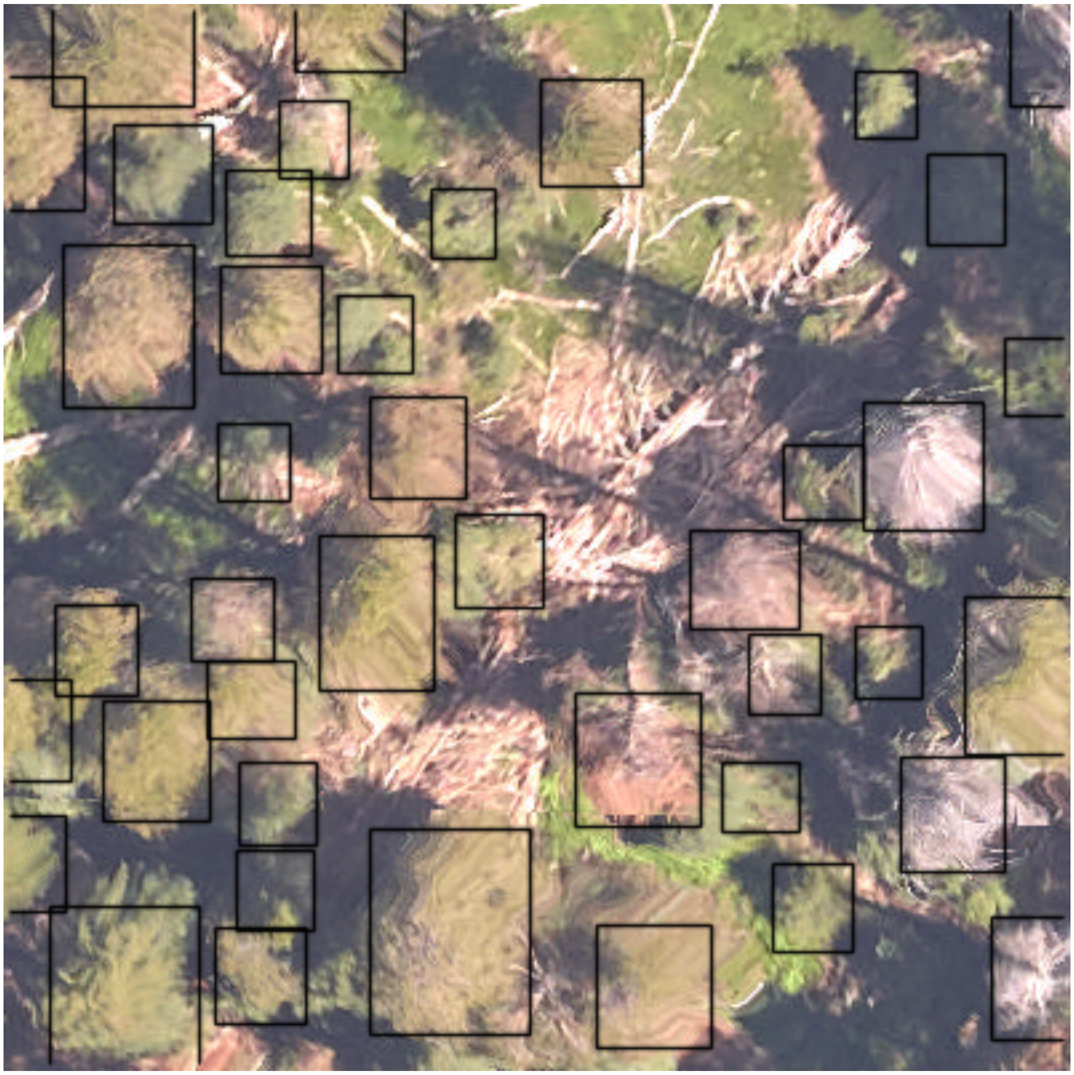

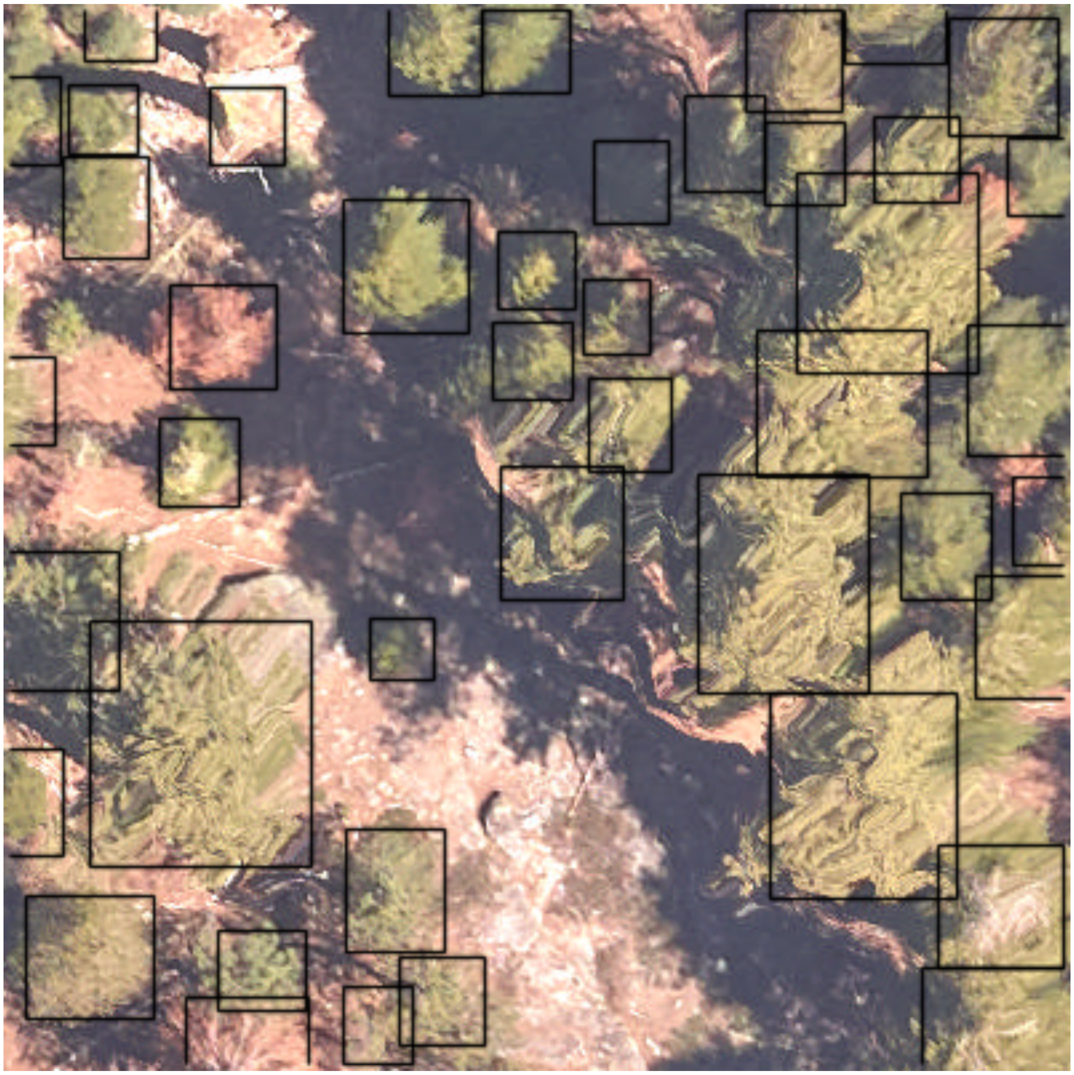

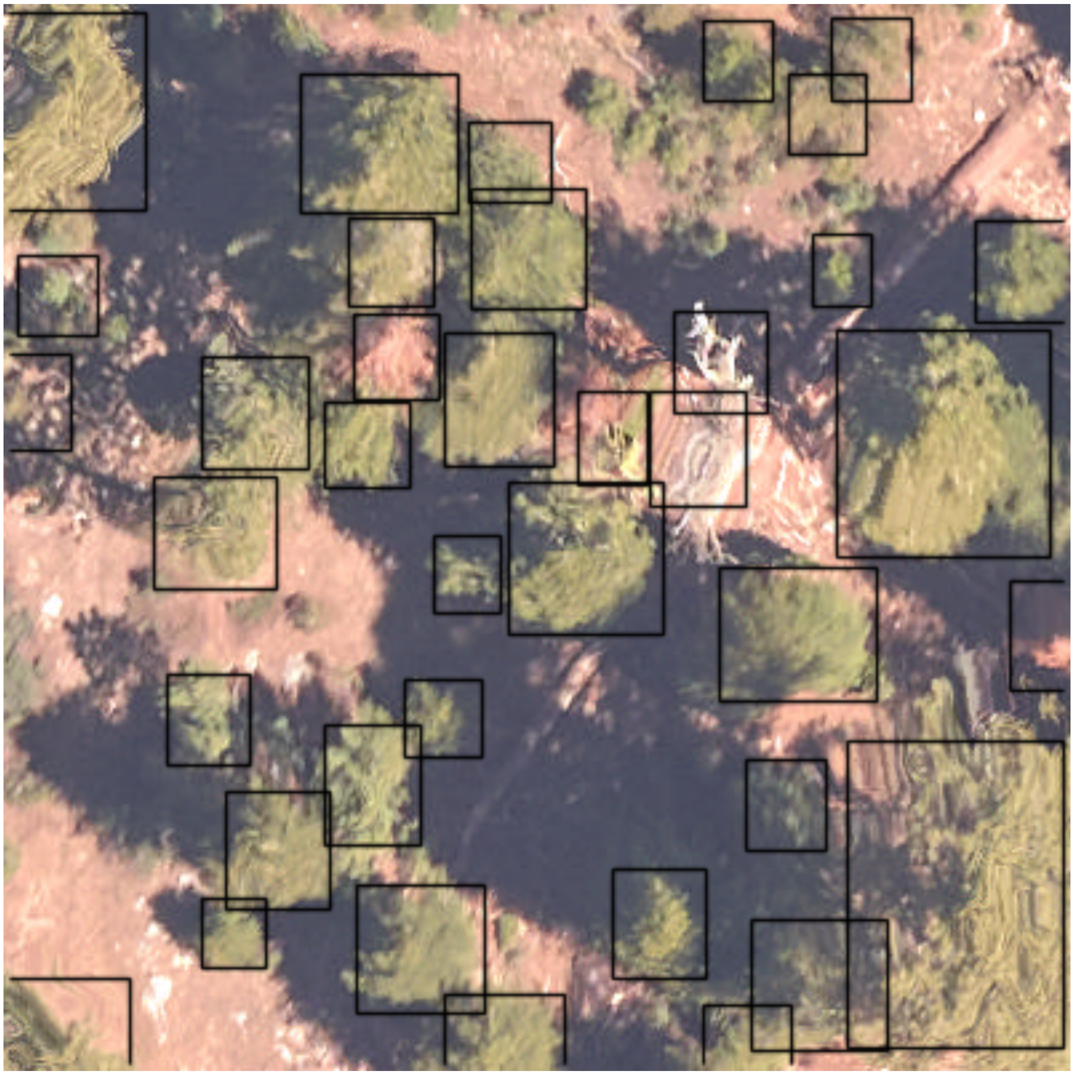

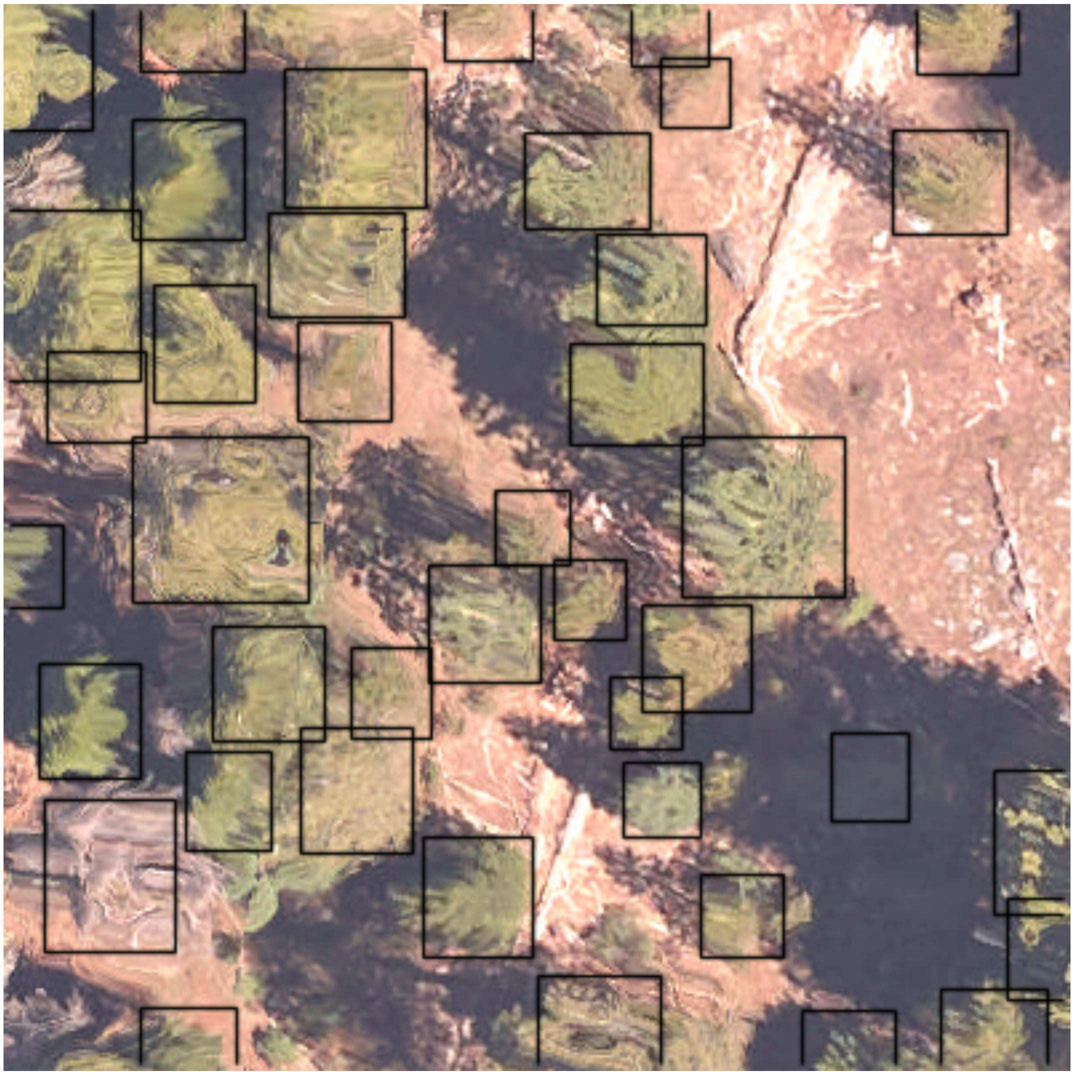

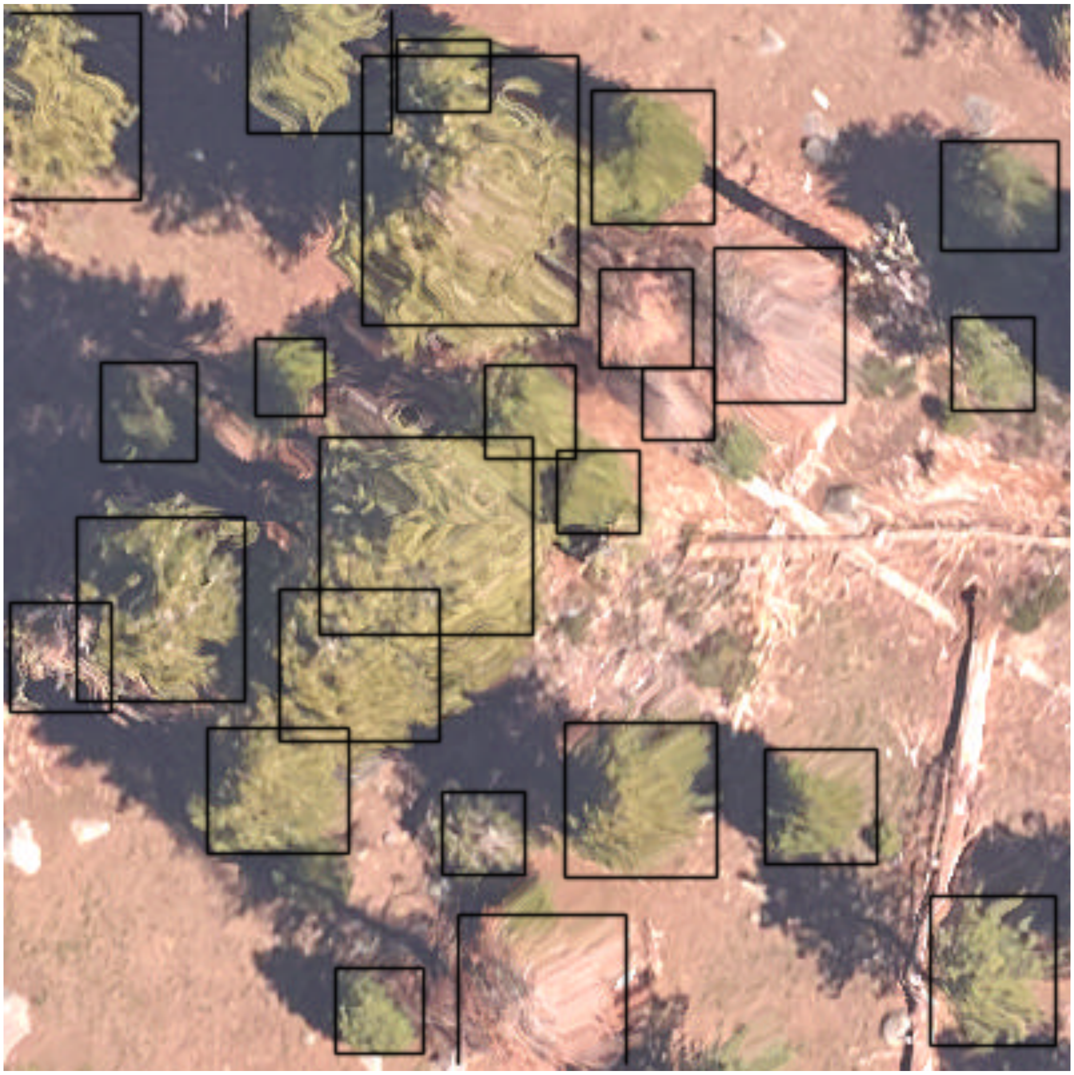

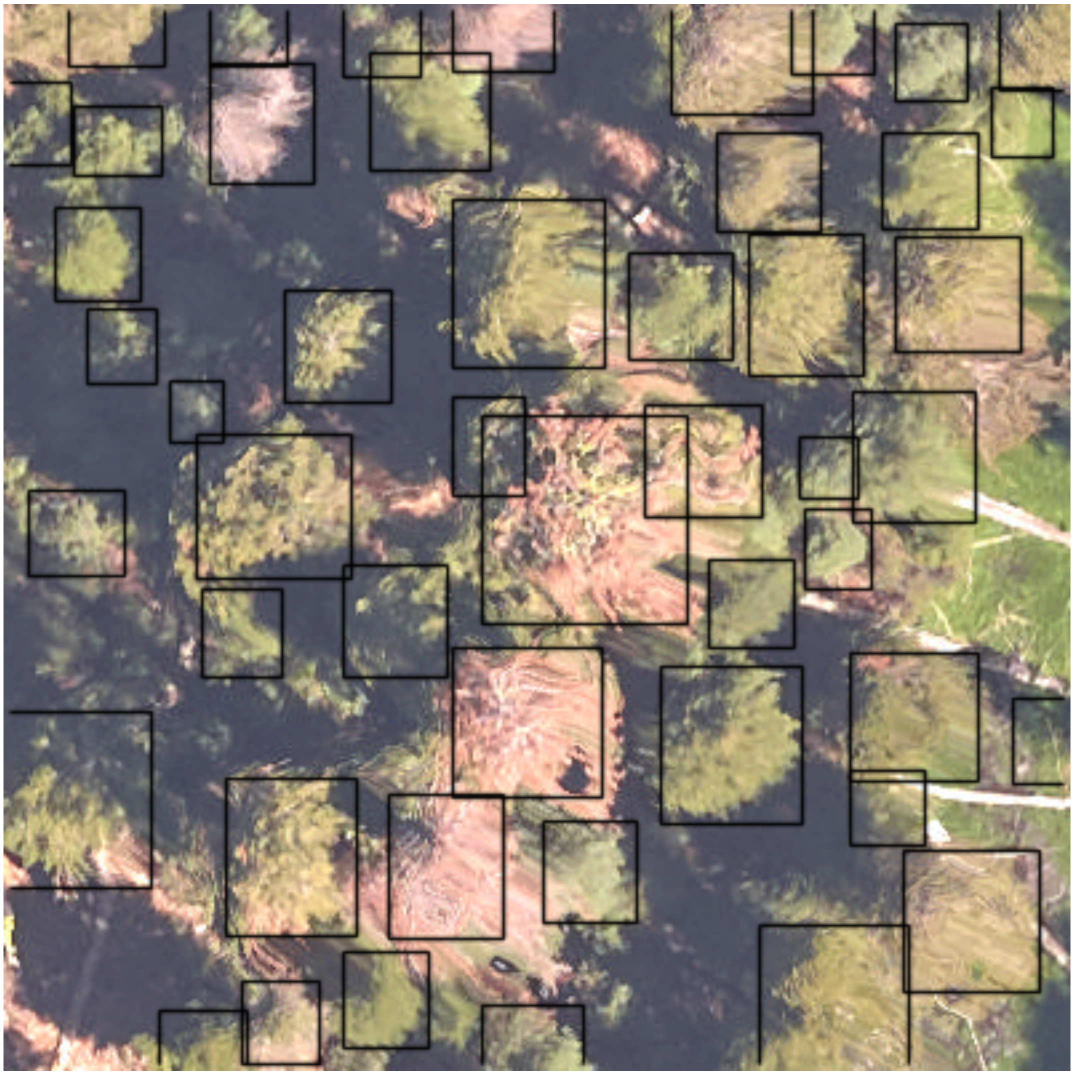

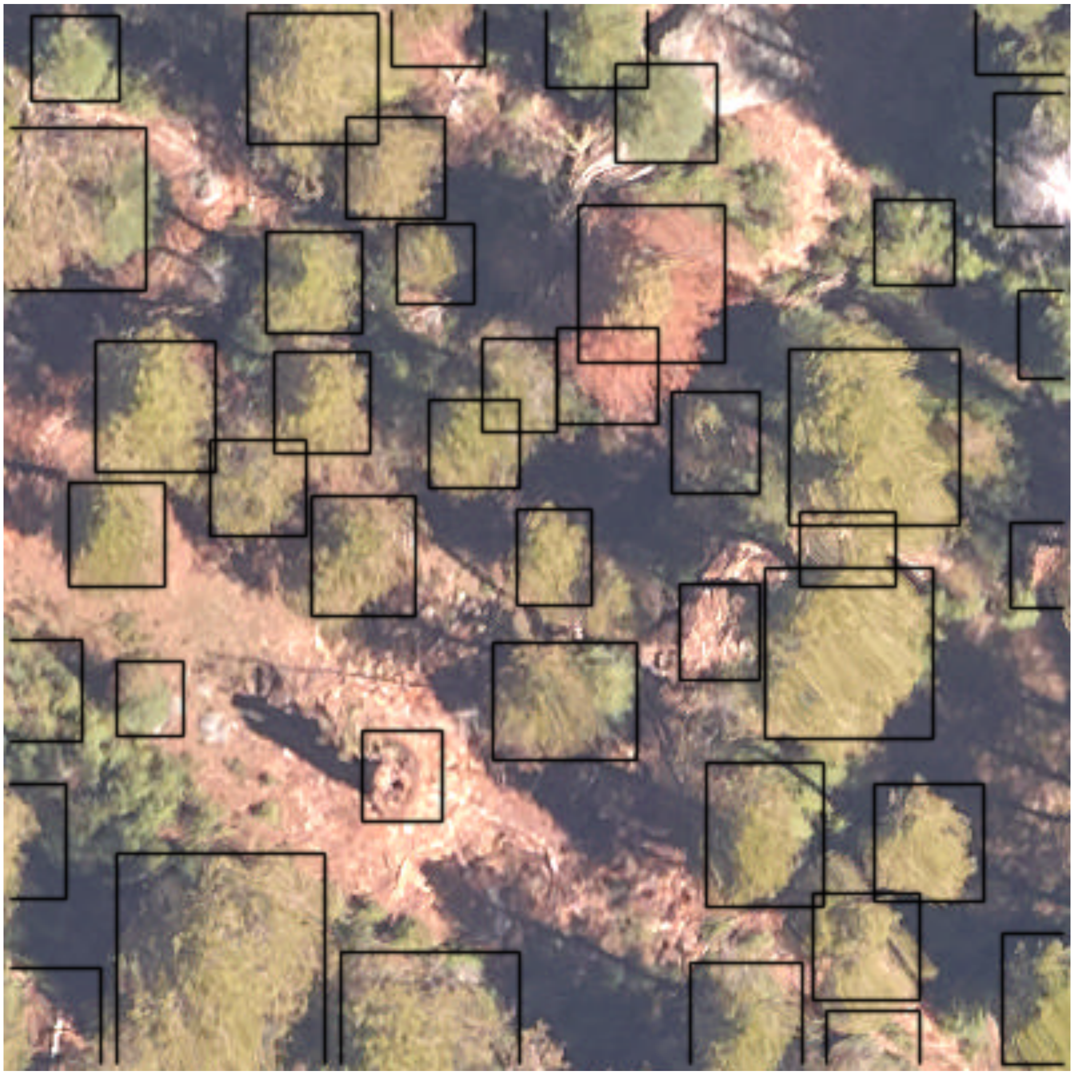

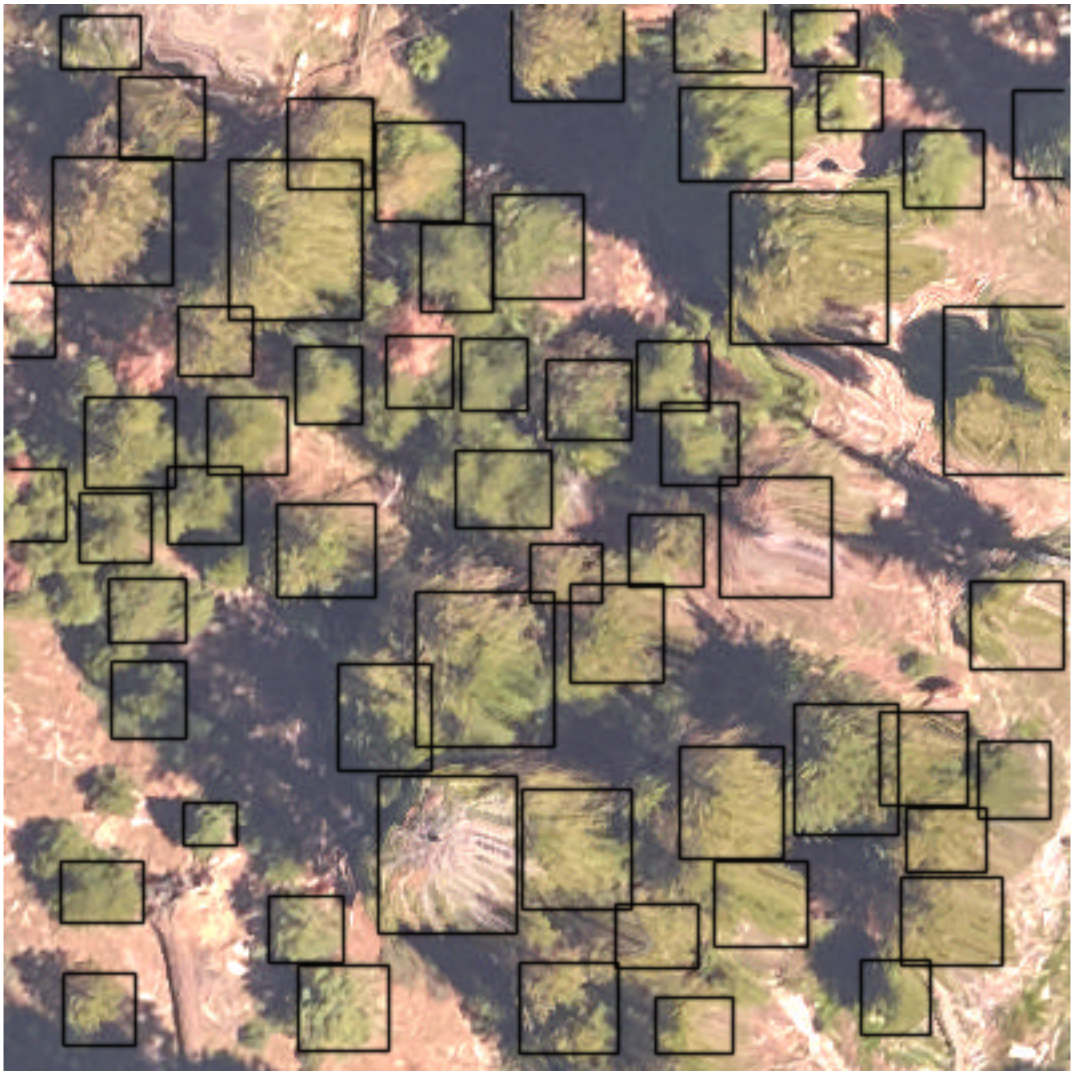

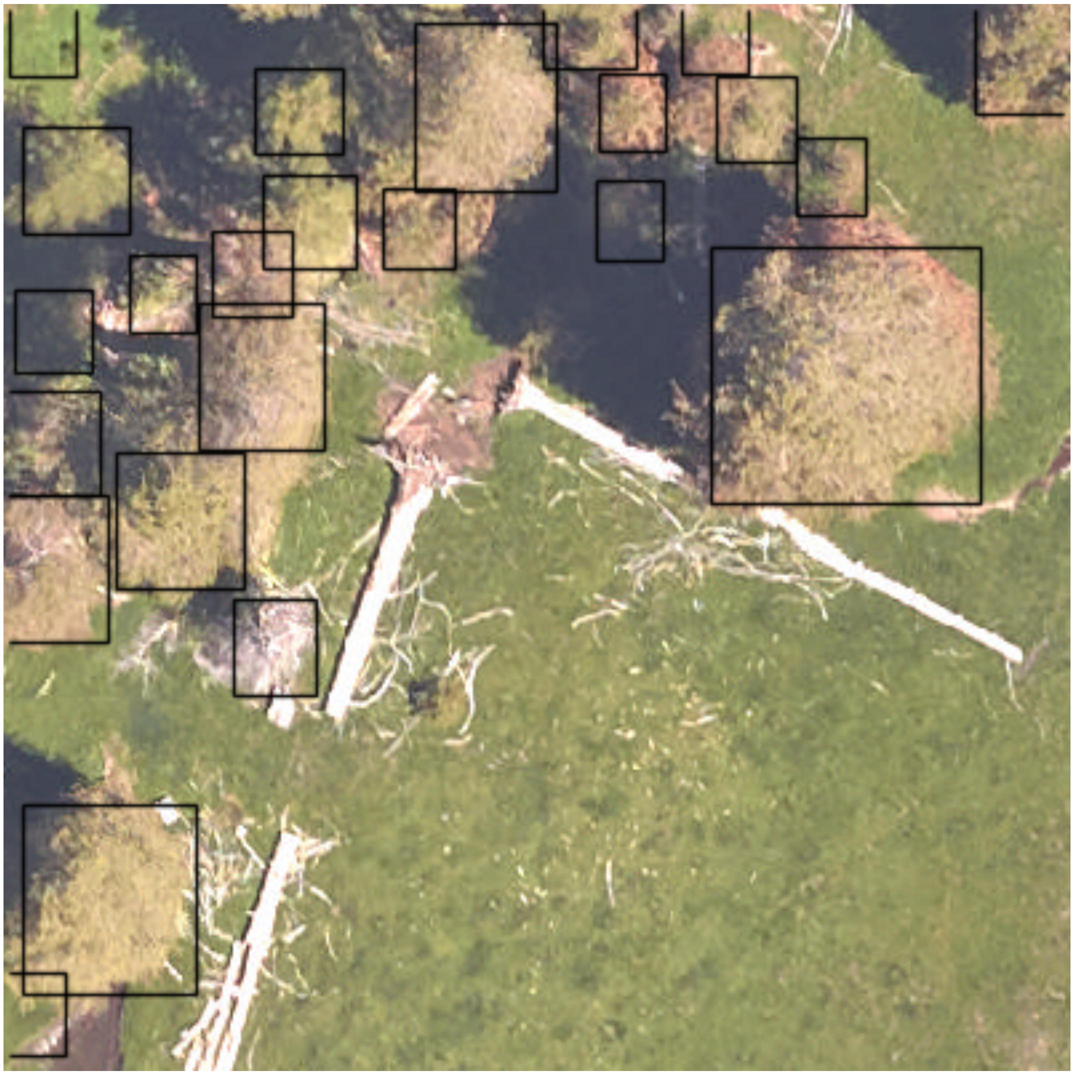

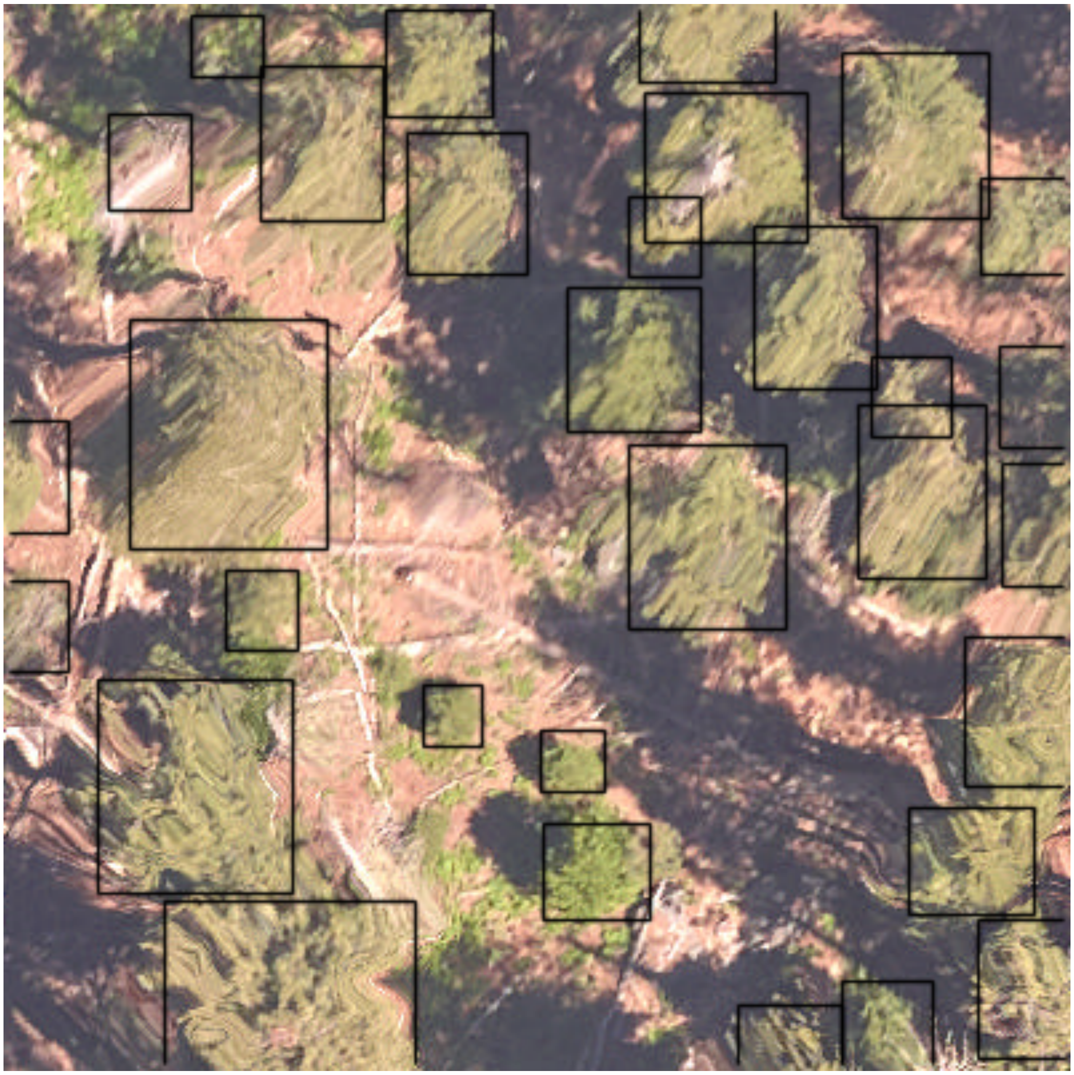

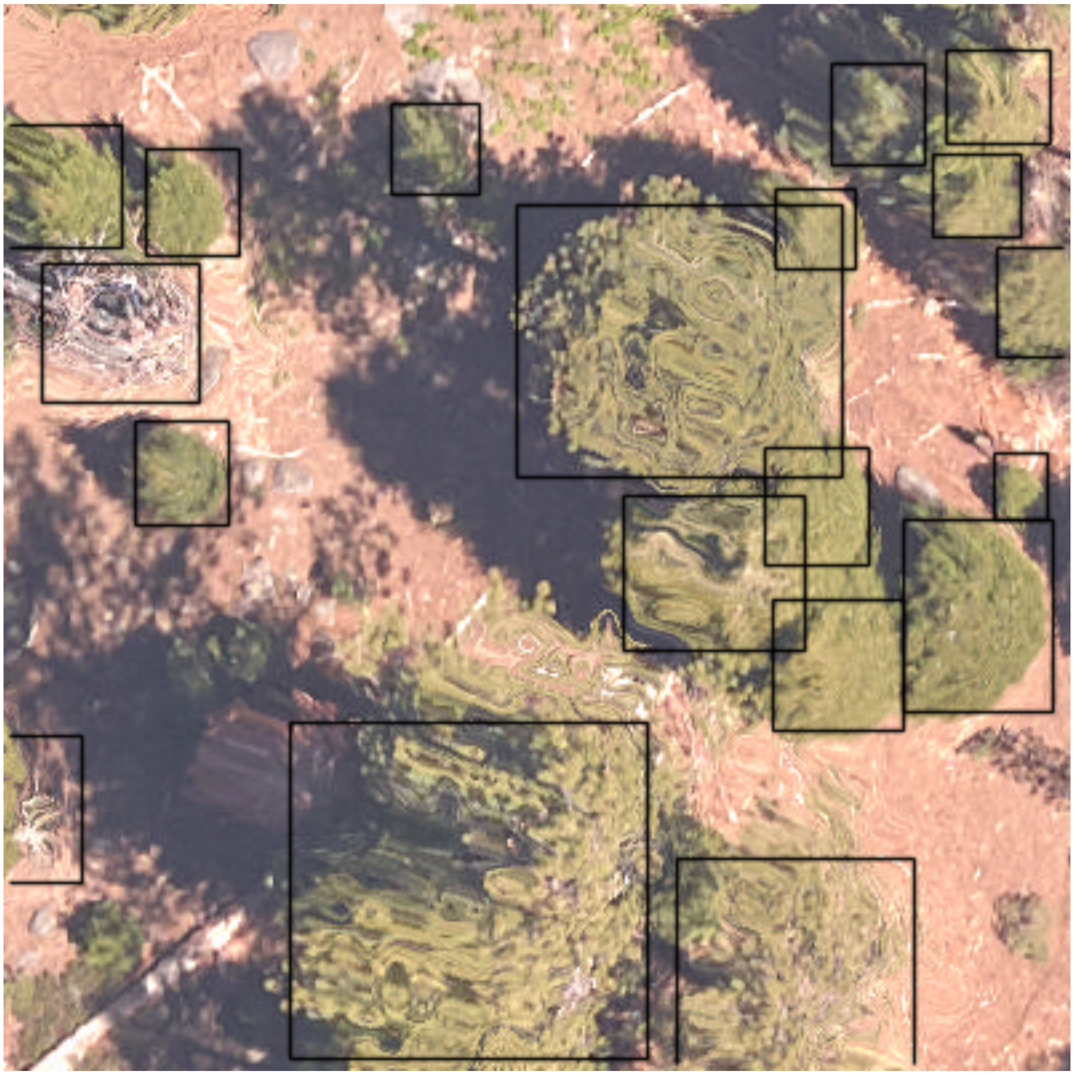

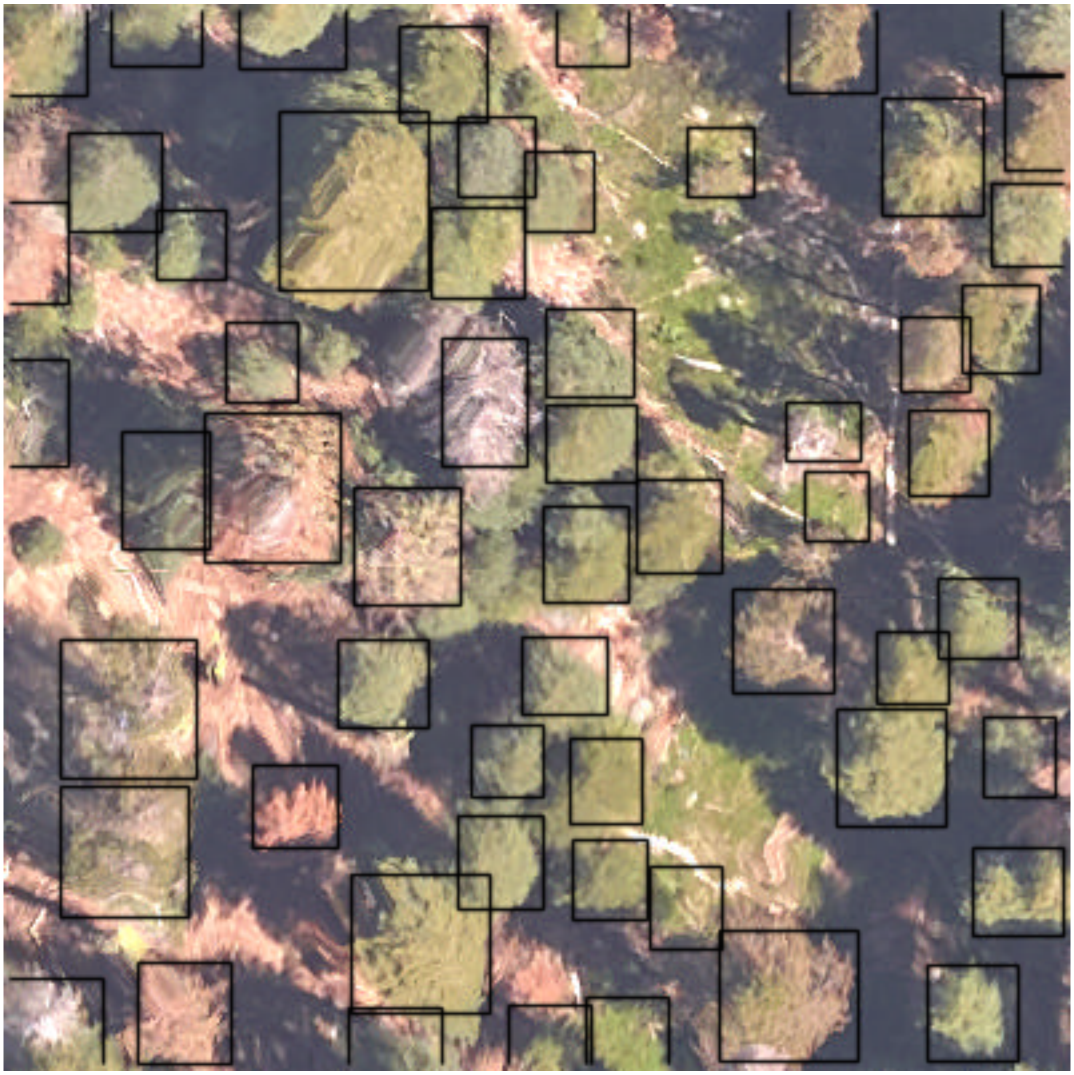

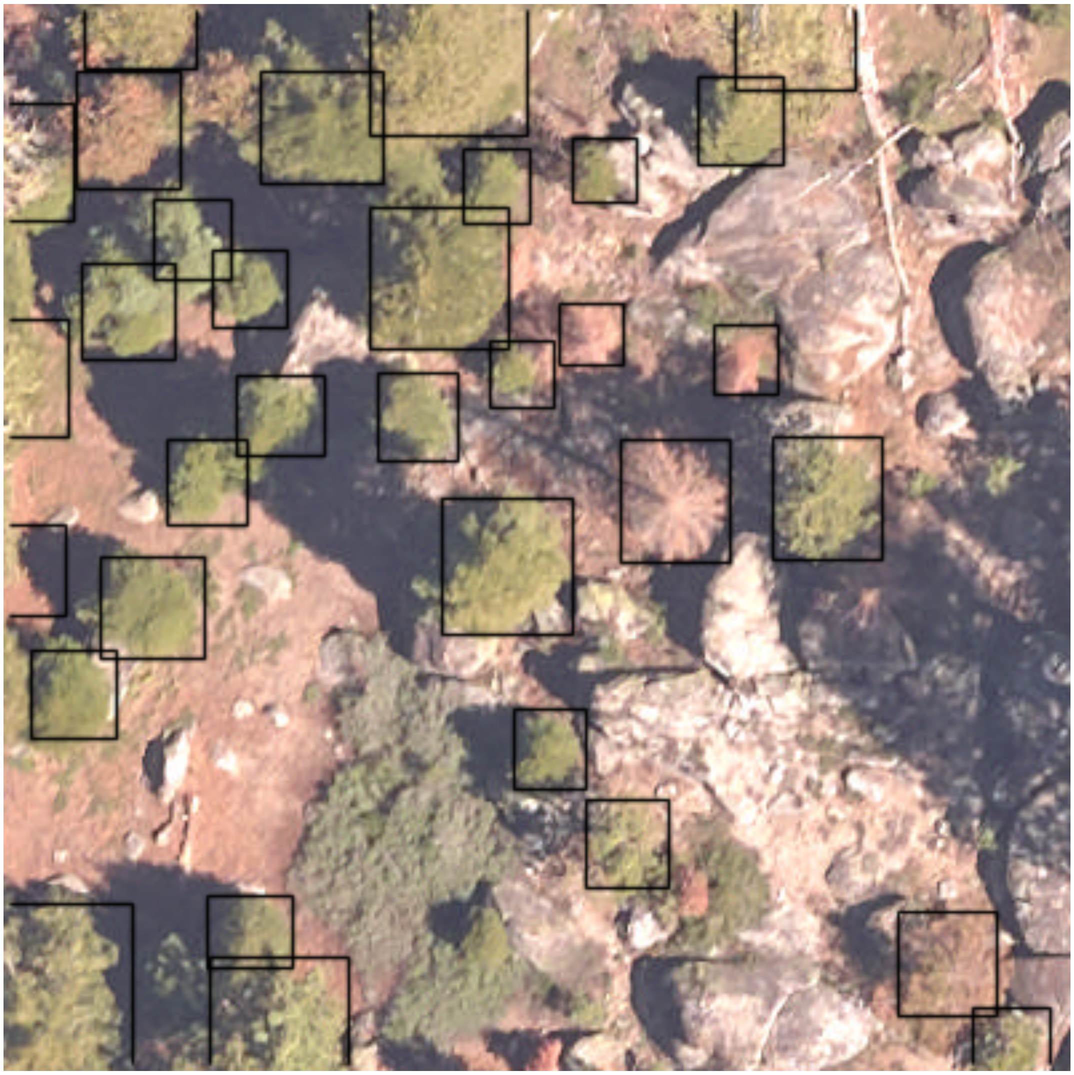

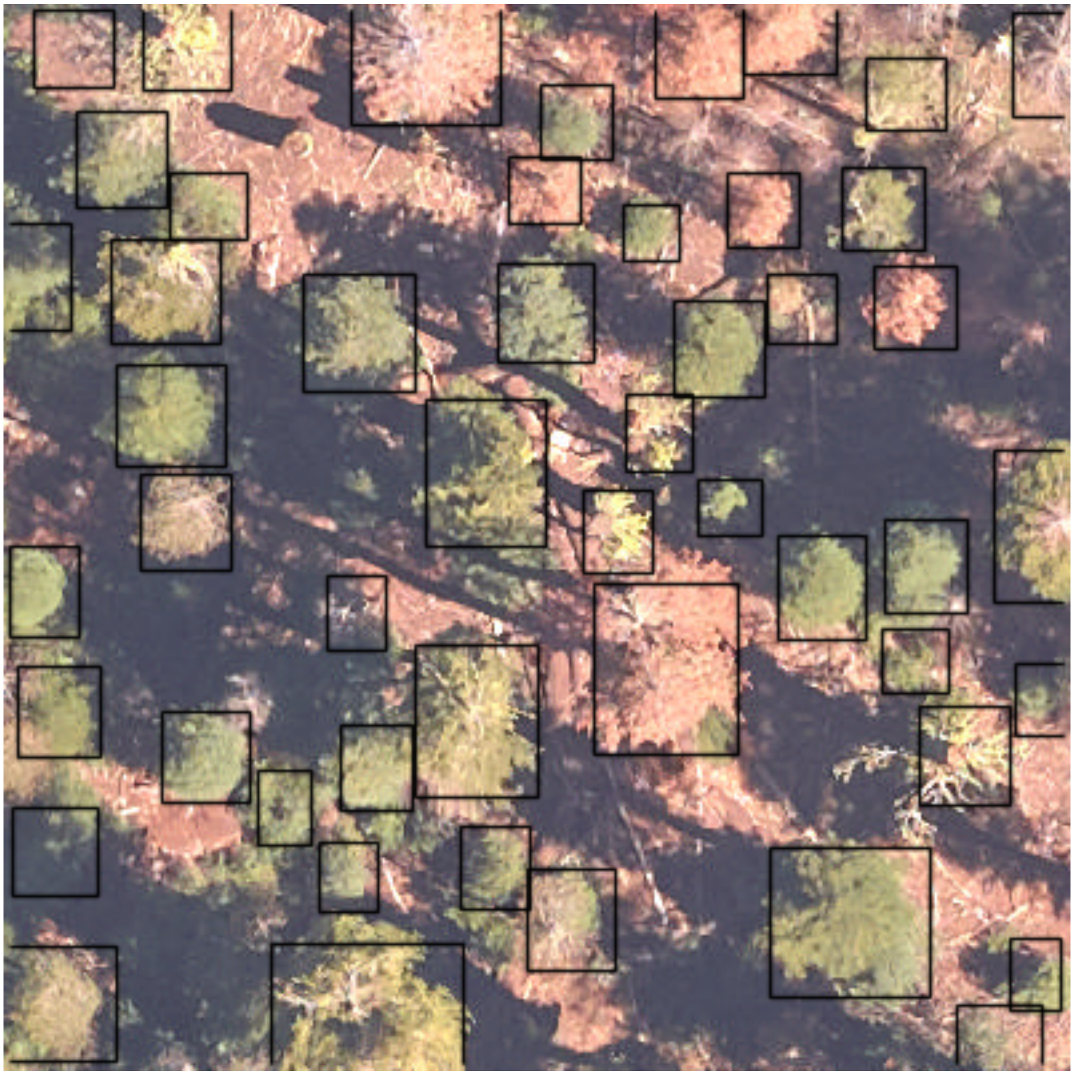

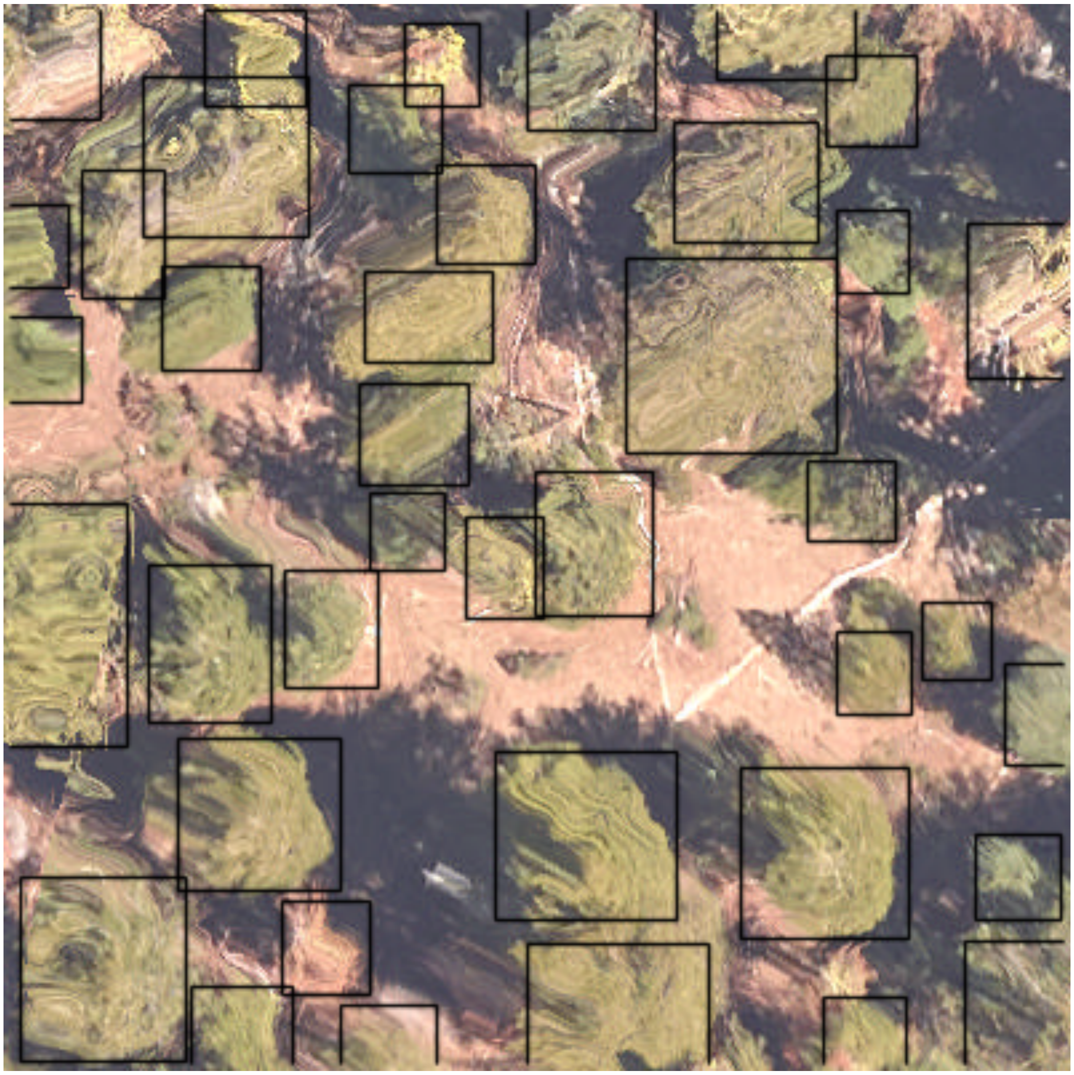

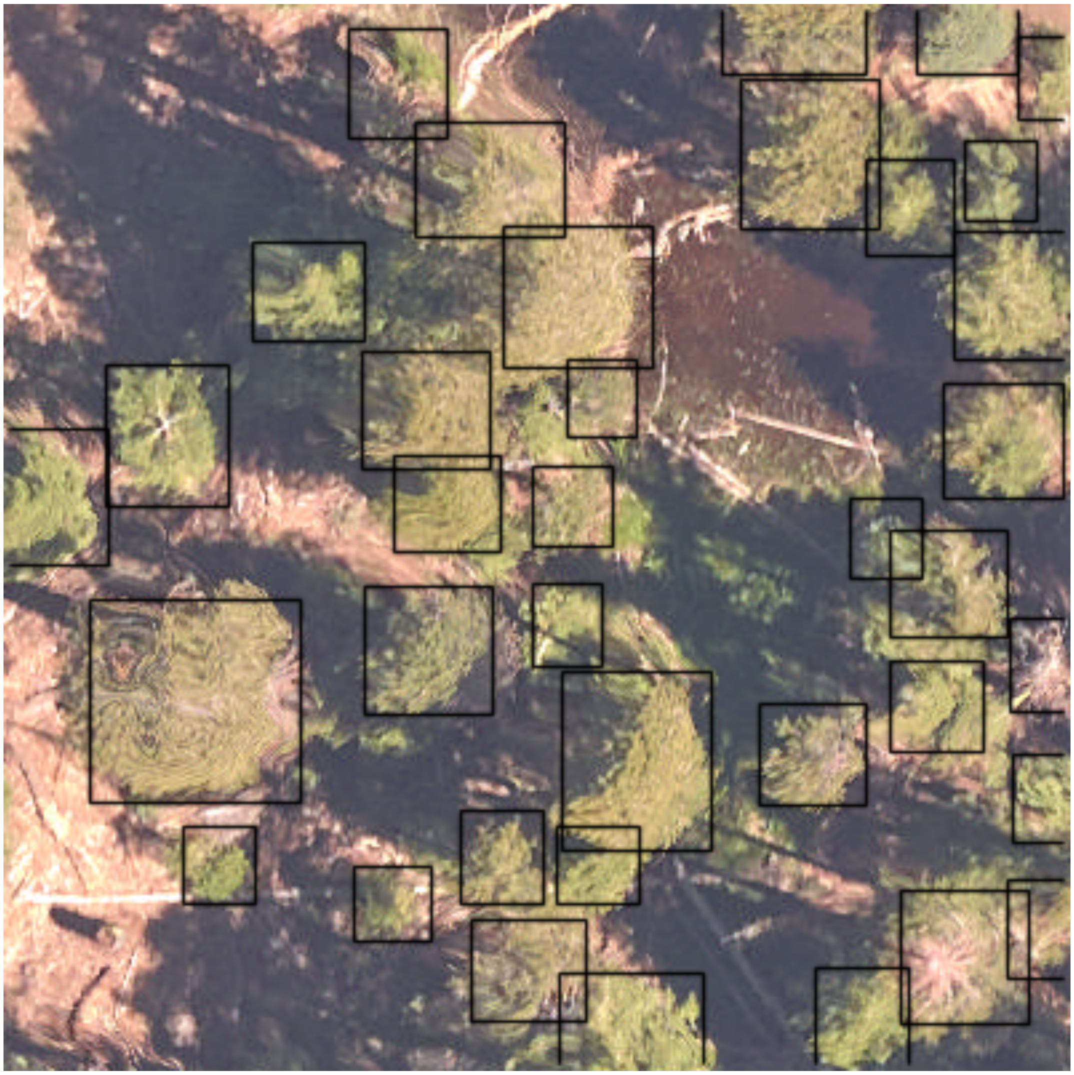

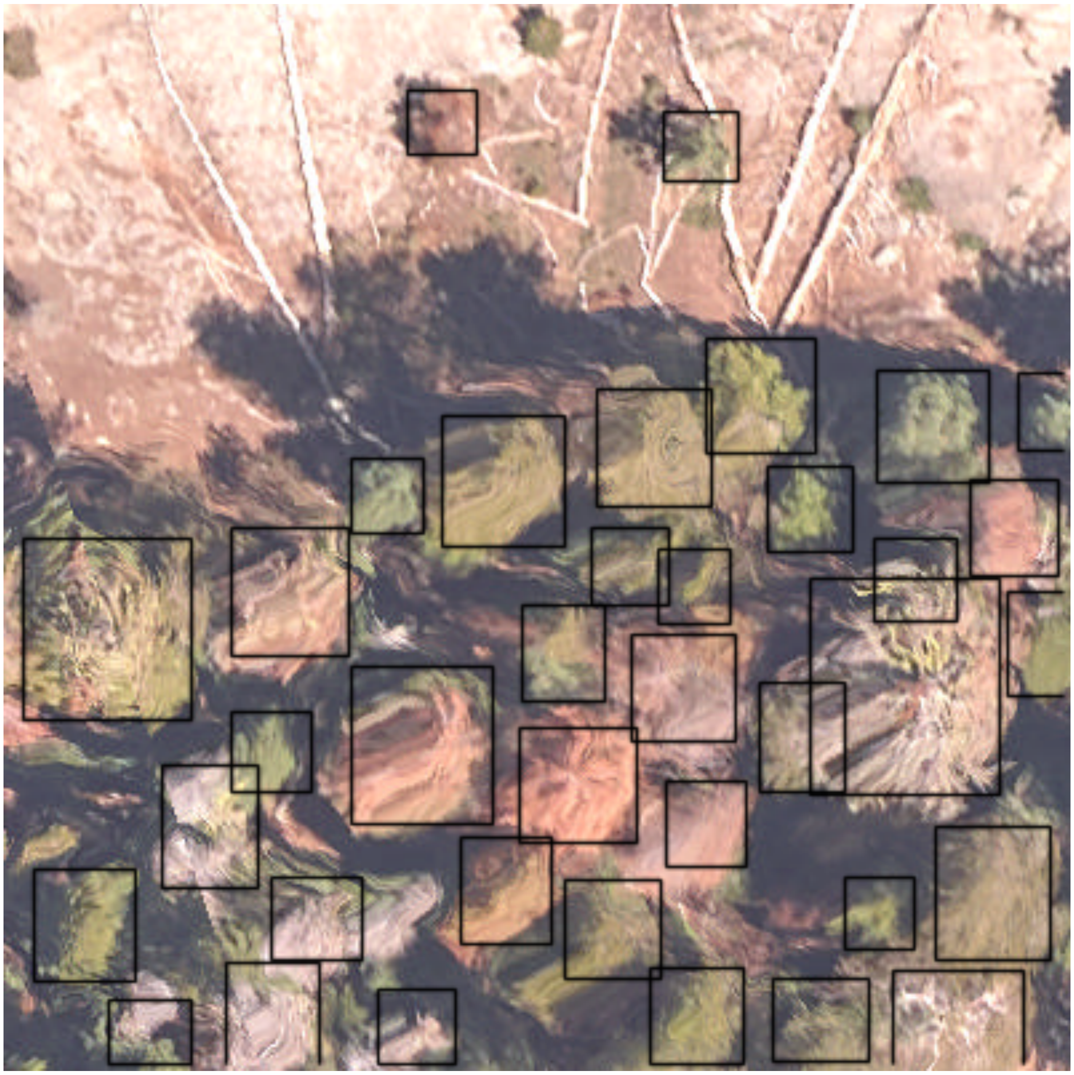
